# Isolation and In vitro Characterization of BchE, the Cobalamin-Dependent Anaerobic Magnesium Protoporphyrin IX Monomethylester Cyclase

**DOI:** 10.64898/2026.04.29.721654

**Authors:** Nicholas J. York, Xuekai Zhang, Squire J. Booker

**Affiliations:** Department of Chemistry, The Pennsylvania State University, University Park, PA, USA; Department of Chemistry, School of Arts and Sciences, University of Pennsylvania, Philadelphia, PA, USA; Department of Biochemistry & Molecular Biology, The Pennsylvania State University, University Park, PA, USA; Department of Biochemistry and Biophysics, Perelman School of Medicine, University of Pennsylvania, Philadelphia, PA, USA; Howard Hughes Medical Institute, Chevy Chase, MD, USA; Department of Biological Engineering, College of Chemical and Biological Engineering, Shandong University of Science and Technology, Qingdao 266590, China

**Keywords:** Radical SAM, Cobalamin, Bacteriochlorophyll, S-adenosylmethionine, Magnesium protoporphyrin IX monomethyl ester

## Abstract

The radical *S*-adenosylmethionine (SAM) superfamily comprises more than 800,000 enzymes that use [Fe_4_S_4_] clusters to initiate radical chemistry that mediates an exceptionally broad range of chemical transformations. Within this superfamily, cobalamin (Cbl)-dependent radical SAM enzymes constitute a major subclass predominantly associated with methylation reactions. However, several notable members catalyze non-methylase reactions, for which the mechanistic role of Cbl is poorly understood. Bacteriochlorophyll biosynthesis enzyme BchE is a Cbl-dependent radical SAM enzyme that catalyzes a six-electron oxidation of Mg-protoporphyrin IX monomethylester (MPE) to protochlorophyllide (PChlide), installing a ketone and forming the fifth ring of bacteriochlorophyll under anaerobic conditions. Although prior in vivo and in vitro studies have demonstrated a requirement for Cbl, SAM, and a low-potential reductant, detailed mechanistic analysis has been impeded by the inability to obtain soluble, catalytically active enzyme. Here, we report the successful isolation and spectroscopic characterization of BchE, enabling the first *in vitro* reconstitution of its enzymatic activity. Using both chemical and biological reducing systems, we observe the formation of PChlide along with proposed reaction intermediates and several off-pathway products. These results provide new insight into the oxidative chemistry mediated by Cbl in non-methylase radical SAM enzymes and establish BchE as a tractable model for elucidating how cobalamin is deployed in this understudied subclass.

**Significance Statement:** Bacteriochlorophyll biosynthesis enzyme BchE is a cobalamin-dependent radical SAM enzyme that catalyzes a six-electron oxidation of Mg-protoporphyrin IX monomethylester (MPE) in the absence of molecular oxygen. This reaction entails the oxidation of carbon 13^1^ from a methylene to a ketone and the formation of a carbon-carbon bond between carbons 13^2^ and 15 to generate bacteriochlorophyll’s fifth ring. How BchE performs this reaction has remained mysterious since its identification in the early 2000s, largely because it has been refractory to purification and in vitro characterization. This work describes the first purification of BchE, its spectroscopic characterization, and biochemical studies, providing key mechanistic insight and information on the likely state of cobalamin during the reaction.

## Introduction

The radical S-adenosylmethionine (SAM) superfamily of enzymes catalyzes a wide assortment of reactions through the intermediacy of organic radicals. To date, the superfamily (PF04055) comprises more than 800,000 unique sequences of enzymes that catalyze more than 100 distinct reactions.(1–3) These reactions include the generation of glycyl radical cofactors, 1,2-cross-migrations of hydrogen atoms and functional groups, thiolation and methylthiolation, oxidation and oxidative decarboxylation, complex rearrangements, epimerization, methylation, the formation of C–S and C–N crosslinks, and the formation of C(sp^3^)–C(sp^3^) and C(sp^2^)–C(sp^3^) bonds, to name a few. All radical SAM (RS) enzymes contain at least one redox-active [Fe_4_S_4_] cluster, which is required for the reductive cleavage of SAM to generate, most often, a 5’-deoxyadenosyl 5’-radical (5’-dA•) (**Figure 1A**).

**Figure 1.**
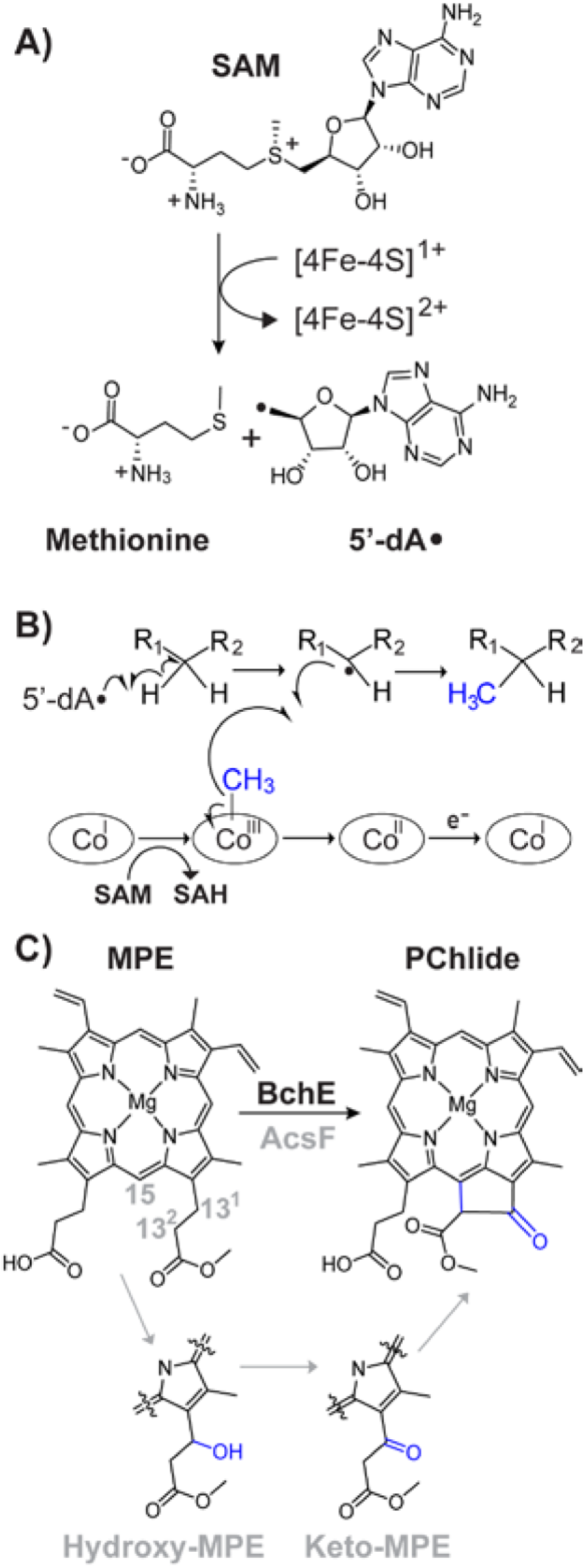
**A**) The [4Fe-4S] cluster-dependent reductive cleavage of *S*-adenosylmethionine (SAM) to generate methionine and a 5’-deoxyadenosyl radical (5’-dA·). **B**) Representative mechanism for the methylation of an inert carbon center by cobalamin-dependent radical SAM enzymes. **C**) Overall reaction catalyzed by BchE, converting Mg-protoporphyrin IX monomethylester (MPE) to protochlorophyllide (PChlide). Also depicted are the hydroxy and keto intermediates observed during the AcsF-catalyzed reaction.(9)

The expansive repertoire of reactions catalyzed by RS enzymes is made possible by additional cofactors that some RS enzymes employ, including auxiliary iron-sulfur (Fe-S) clusters, pyridoxal 5’-phosphate, and cobalamin (Cbl).(4–7) Cbl-containing RS enzymes represent one of the largest subclasses within the superfamily, currently with over 80,000 predicted sequences.(3, 4) Most of these enzymes are presumed to be methylases that add methyl groups to unactivated carbon or phosphinate phosphorus centers, as depicted in **Figure 1B**. In Cbl-dependent RS methylases, a cob(I)alamin species acquires a methyl moiety from SAM, forming methylcobalamin (MeCbl). In a second step, the 5’-dA• abstracts a target hydrogen atom from the substrate. The resulting substrate radical performs an S_H_2 reaction on the methyl moiety of MeCbl, inducing a homolytic cleavage of the cobalt-carbon bond to afford the methylated species and cob(II)alamin. Upon the reduction of cob(II)alamin, the resulting cob(I)alamin species is remethylated by another molecule of SAM.(4, 8)

A few Cbl-dependent RS enzymes that do not perform methylation reactions have been identified. These include the structurally characterized OxsB, which catalyzes a contraction of the five-membered ring of 2′-deoxyadenosine-5′-monophosphate to form oxetanocin A.(10, 11) Others include HpnJ, involved in hopanoid biosynthesis, and PbsB, which modifies a ribosomally synthesized and posttranslationally modified peptide (RiPP) by crosslinking two *ortho*-tyrosines to form a macrocycle.(12, 13) Cbl-dependent RS enzymes are also proposed to form the cyclobutane rings of ladderane lipids formed in anammox bacteria and the cyclopentane rings of glycerol tetraether lipids found in archaea.(14, 15)

BchE, a Cbl-dependent RS enzyme involved in the biosynthesis of bacteriochlorophyll, also does not catalyze a methylation reaction. Rather, this enzyme catalyzes the 6e-oxidation of Mg-protoporphyrin IX monomethylester (MPE) to protochlorophyllide (PChlide), a transformation that entails ketone formation on carbon 13^1^ and ring formation between carbons 13^2^ and 15, as shown in **Figure 1C**. This reaction has been purported to involve hydroxy- and keto-MPE intermediates on carbon 13^1^ (**Figure 1C)**.(16, 17) These species were observed in the reaction of *Rubrivivax gelatinosus* AcsF, the aerobic counterpart to BchE that utilizes a diiron active site to insert oxygen from O_2_.(9) Based on isotopic studies, BchE is proposed to function in anaerobic conditions and utilize oxygen from water to generate the 13^1^ ketone.(18, 19) In 2000, Gough et al. determined that Cbl was necessary for this reaction, and the following year, Sofia et al. annotated BchE as an RS enzyme in their pivotal bioinformatics study.(20, 21)

The characterization of BchE and its reaction has been limited almost exclusively to *in vivo* studies, largely due to the enzyme’s historical insolubility. Ouchane et al. isolated urea-solubilized BchE but it lacked evidence of activity or cobalamin incorporation.(22) Purifications were also attempted by Wiesselmann et al., but were ultimately unsuccessful.(23) In lieu of an isolated enzyme, the authors developed an *in vitro* assay to study the BchE reaction using the cell lysate of a *bchH^−^*mutant strain of *Rhodobacter capsulatus*, which natively expresses BchE. The *bchH* gene encodes part of the BchIDH magnesium chelatase complex, which is responsible for inserting a magnesium cation into protoporphyrin IX (PP IX). After magnesium insertion, BchM methylates the carboxyl group at C13, forming MPE. With the pathway for MPE (and ultimately bacteriochlorophyll *a*) production disrupted, the *bchH^−^* mutant strain expresses native BchE but lacks its substrate. Upon incubating the lysate of these cells with MPE, PChlide formation was observed, consistent with active BchE. The authors determined that the reaction was dependent on both SAM and reductant, while being inhibited by SAH, O_2_, and iron-chelating agents. Furthermore, an unknown soluble component native to *R. capsulatus* was required for enzyme activity. The authors did not identify the component but speculated that it was likely a redox protein specific to BchE.

To date, OxsB is the only Cbl-dependent RS enzyme that is not a methylase to be structurally characterized and mechanistically investigated.(10, 11) The ability to study BchE therefore represents a significant advance toward understanding the role of cobalamin in this understudied class of enzymes. Here, we report the isolation and spectroscopic characterization of BchE and demonstrate its in vitro catalytic activity using multiple reducing systems. Under these conditions, we observe formation of the PChlide product, along with proposed reaction intermediates and several off-pathway species.

## Results

### Rubrivivax pictus BchE purification and spectroscopic characterization

The gene for *Rubrivivax pictus* BchE was synthesized to encode a fusion protein with a TEV protease-cleavable C-terminal maltose-binding protein (MBP) tag and a His_10_ affinity tag (*i.e*., *Rp*BchE-TEV-MBP-His_10_). The full peptide and DNA sequence are given in the *Supporting Information*. *Rp*BchE was heterologously overexpressed in an *E. coli* BL21(DE3) strain containing the pDB1282 and pBAD42-BtuCEDFB plasmids, which facilitate iron-sulfur (FeS) cluster biosynthesis and cobalamin uptake and transport, respectively.(24, 25) Production of *Rp*BchE-MPB (≈100 kDa) was observed by SDS-PAGE after an overnight incubation at 18 °C (**Fig. S1**). The bacteria were harvested and lysed, and the protein was purified in an anaerobic chamber by immobilized metal affinity chromatography (IMAC) using Ni-NTA resin. The eluted protein was blackish-brown, consistent with incorporation of an FeS cluster. We have found that it is common for co-expressed *isc* proteins from the pDB1282 plasmid to bind to the Ni-NTA column and co-elute with the target protein. Therefore, we opted to use a cleavable affinity tag for further purification. Incubation with TEV protease was performed overnight, along with chemical reconstitution of the FeS cluster and cobalamin. The reconstituted protein mixture was then passed through the Ni-NTA column a second time, where the cleaved MBP-free BchE would flow through for collection. Meanwhile the contaminate (presumably *isc*-related) proteins and cleaved MBP-His_10_ fragments were retained on the resin. SDS-PAGE analysis of a typical purification is shown in **Fig. S1**. Additional purification was achieved by size-exclusion chromatography.

Bound Cbl was quantified by UV-vis spectroscopy after BchE treatment with potassium cyanide; however, mass spectrometry analysis shows that the bound Cbl is primarily adenosylcobalamin (AdoCbl) (*vide infra*). Given that AdoCbl was not predicted to be the active form of the Cbl cofactor in the reaction, BchE was treated in several ways to eliminate it. We initially attempted to photolyze AdoCbl-bound BchE to cleave the Co-C_Ado_ bond, but were unable to fully convert the cofactor to a uniform state. Efforts to isolate BchE lacking Cbl were unsuccessful because overexpression without the pBAD42-BtuCEDFB plasmid resulted in no soluble protein. Incubation of AdoCbl-bound BchE with excess OHCbl followed by gel-filtration chromatography resulted in no change in the AdoCbl content. Therefore, we overproduced BchE in a *ΔbtuR* knockout *E. coli* BL21(DE3) strain, which is unable to produce AdoCbl.(24) The *Rp*BchE-MBP construct was cloned into a pRham vector (Lucigen), which was used, along with the pDB1282 and pBAD42-BtuCEDFB plasmids, to transform this new strain. Overexpression and purification were repeated with the new strain.

The MBP-free *Rp*BchE was characterized by SDS-PAGE (**Figure 2A**) and quantified by the Bradford assay, yielding 0.9 mg of pure protein per liter of culture. UV-vis spectroscopy of the purified protein shows shoulders at 405 nm, consistent with an FeS cluster, and 473 nm, indicative of Cbl (**Figure 2B**). Treatment with potassium cyanide reveals a cyanocobalamin UV-vis spectrum (**Figure 2C**). Cbl and iron were quantified as 0.66 ± 0.02 and 2.9 ± 0.3 per protein, respectively, suggesting incomplete cofactor incorporation. However, normalization of the Fe concentration to that of Cbl results in 4.2 Fe/Cbl, which agrees with expectations of one [Fe_4_S_4_] cluster and one Cbl per protein. Due to this discrepancy, protein concentrations are normalized to Cbl concentration throughout this work. No AdoCbl or MeCbl is detected by mass spectrometry from *Rp*BchE produced in the *ΔbtuR* knockout strain.

**Figure 2.**
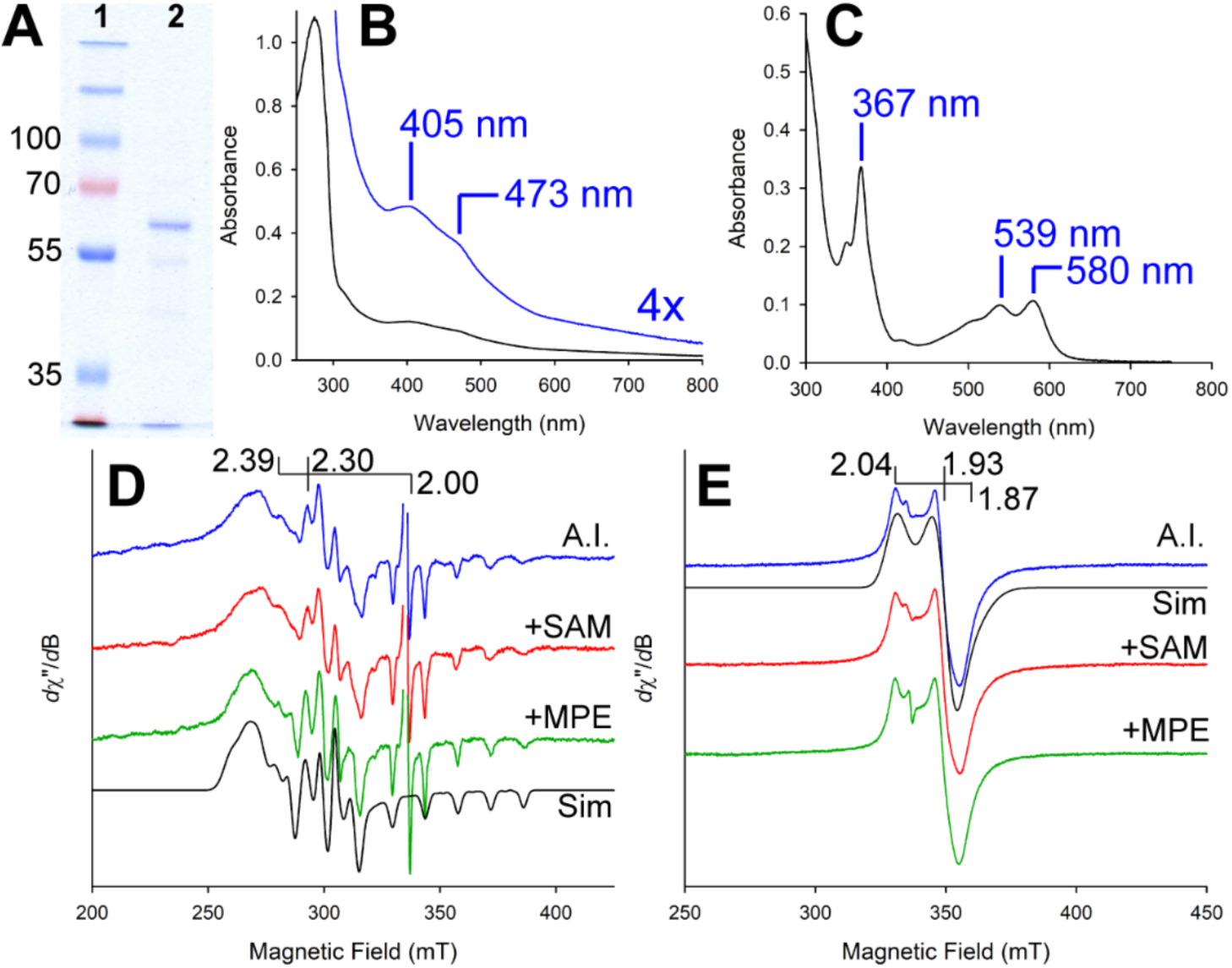
**A**) SDS-PAGE analysis of *Rp*BchE. Lane 1, molecular mass markers; lane 2, purified *Rp*BchE after removal of the MBP-tag. (**B**) UV-vis spectrum of purified *Rp*BchE showing absorption features consistent with an FeS cluster and OHCbl. (**C**) UV-vis spectrum of purified *Rp*BchE following treatment with potassium cyanide, demonstrating conversion to cyanocobalamin. (**D**) EPR spectra of *Rp*BchE in the absence of reductant, revealing a cob(II)alamin signal. (**E**) EPR spectra of *Rp*BchE following dithionite reduction, showing the [4Fe-4S] cluster. Traces correspond to as-isolated *Rp*BchE (blue), *Rp*BchE + SAM (red), and *Rp*BchE + MPE (green). Simulated spectra are shown in black with extrapolated *g*-values.

Electron paramagnetic resonance (EPR) spectroscopy was employed to analyze the FeS cluster and Cbl cofactors and their interactions with SAM and MPE. The spectrum in **Figure 2D** (*blue trace)* shows the EPR spectrum of as-isolated RpBchE (300 µM) recorded at 70 K. The addition of 1 mM SAM (**Fig. 2D**, *red trace*) results in no spectral change, while the addition of 0.6 mM MPE (**Fig. 2D**, *green*) results in spectral line sharpening. This latter spectrum was used for simulations because the improved peak resolution allowed for more confident fitting. The simulated parameters (**Table S1**) include a hyperfine splitting for an *I* = 7/2 nucleus, as observed with cobalt. The *g*-values and hyperfine coupling constants are consistent with a base-off four-coordinate Cbl similar to that observed in the Cbl-dependent RS enzymes as-isolated TokK and TsrM in the presence of SAM.(26, 27) Spin quantification reveals that the cob(II)alamin signal accounts for 56% of the total Cbl concentration. While no spectral changes are observed when BchE is incubated with SAM, peak sharpening is observed with the addition of MPE, with *g*-values and hyperfine couplings remaining unchanged. These observations suggest that neither SAM nor MPE binds to the Cbl. However, MPE likely binds in close proximity, creating a more rigid active site and thereby reducing heterogeneity, as evidenced by the sharper peaks in the associated EPR spectrum.

The EPR spectrum of BchE reduced with dithionite and recorded at 10 K is shown in **Figure 2E**. The spectrum, consistent with that of an [Fe_4_S_4_] cluster, is axial, exhibiting principal *g*-tensor values of 2.04, 1.93, and 1.87. Increasing the pH to 8.5 was necessary to record signals with adequate intensity, suggesting that the reduction potential of the FeS cluster is relatively low. In RS enzymes, the binding of SAM often induces an upfield shift in *g*-values; however, no shift is observed upon incubating the protein with SAM or MPE. Similarly, neither aza-SAM nor *S*-adenosylhomocysteine (SAH) induces a spectral shift. Simultaneous addition of MPE and SAM was attempted to test if an obligate ordered addition of substrates is required for the cluster to bind SAM. However, the resulting spectrum was indistinguishable from that of reduced BchE lacking substrate (**Fig. S2**).

### BchE Enzymatic activity

BchE time-dependent activity was measured (when applicable) by LC-HRMS after quenching with a final concentration of 150 mM H_2_SO_4_ in isopropyl alcohol, conditions that result in the dissociation of the magnesium ion from the bacteriochlorophyll precursors. Initially, we focused on retaining the magnesium in the porphyrin products but found that detection and quantification were improved when using an acidic quench, which ultimately removes the bound magnesium. When 1 mM Ti(III)citrate is used as the low-potential reductant for initiating the radical chemistry, PChlide is produced, as shown in **Figure 3A (***trace I*). The mass spectra for the peaks attributed to MPE and PChlide are displayed in **Figure 3B and C**. Figure 3A (*traces II-IV*) shows that the reaction is dependent on reductant, enzyme, SAM, and MPE.

**Figure 3.**
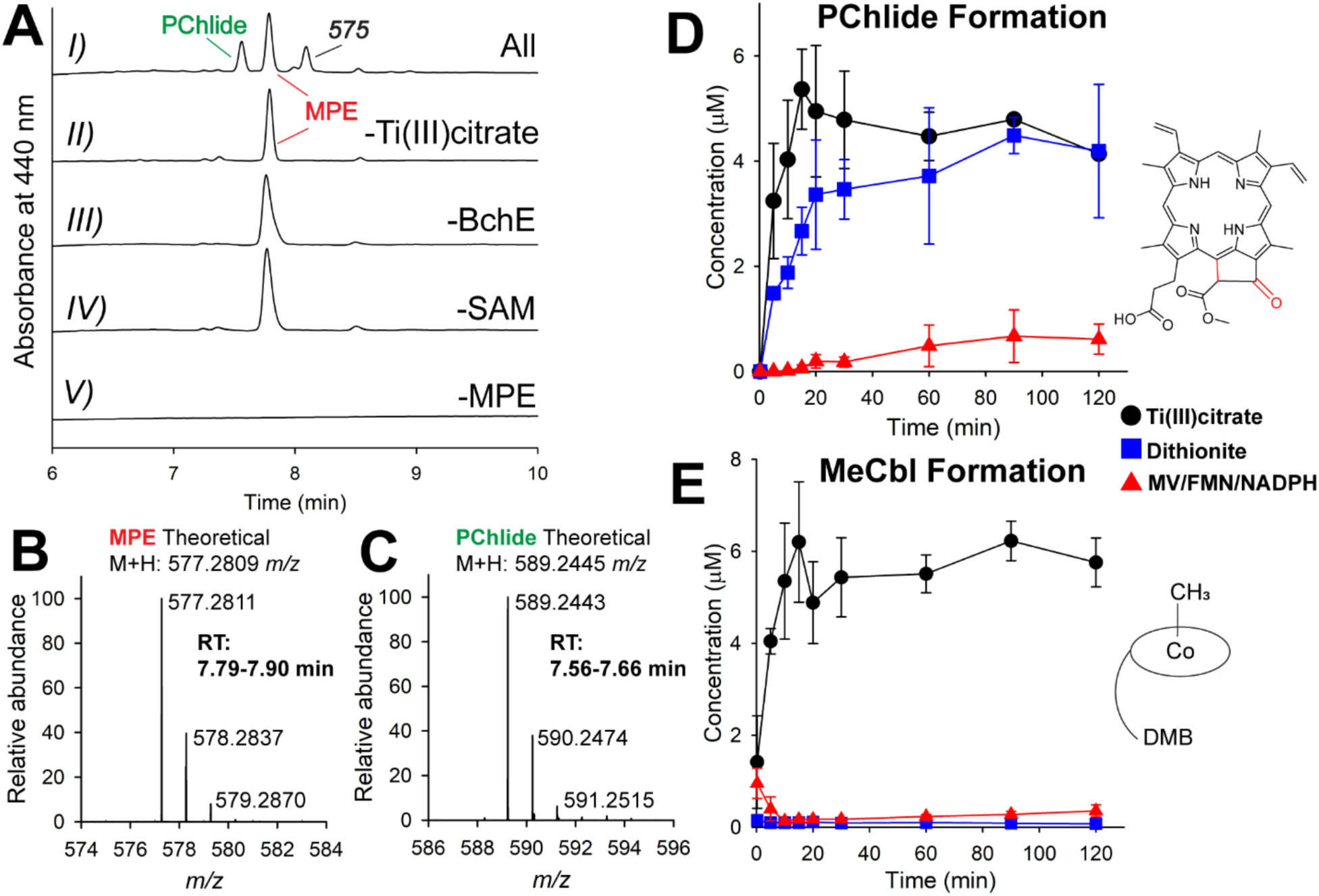
**A**) HPLC chromatograms of quenched BchE reactions under various conditions. Shown are assays containing all components and quenched after 30 min *(I),* as well as control reactions lacking reductant *(II)*, enzyme *(III)*, MPE *(V)*, or SAM *(VI)*. **B**) and **C**) Mass spectra of HPLC peaks corresponding to MPE (B) and PChlide (C), together with the theoretical *m/z* values for the [M+H]^+^ ions of each species. D, E) Quantification of PChlide (D) and MeCbl (E) formation from assays performed in triplicate using Ti(III)citrate (*black circles*), dithionite (*blue squares*), or MV/FMN/NADPH (*red triangles*) as reductants.

BchE contains two redox-active cofactors, Cbl and an [Fe_4_S_4_] cluster. Reduction of the [Fe_4_S_4_] cluster to the +1 oxidation state is needed for the reductive cleavage of SAM, but the active oxidation state of Cbl is unclear. To identify optimal parameters for BchE activity, assays were repeated with alternative chemical reductants, including 1 mM dithionite (DT) and a mixture of 500 µM methyl viologen (MV), 50 µM FMN, and 2 mM NADPH. In **Figure 3D**, the time-dependent formation of PChlide is shown when using these reductants and compared to that when using Ti(III)citrate. The rate of PChlide formation is initially higher with Ti(III)citrate. It is slower in the presence of dithionite, but the magnitude of PChlide formed is ultimately similar. The reaction in the presence of the MV/FMN/NADPH reducing system yields the least activity, with approximately 7-fold lower PChlide accumulation and the slowest observed rate of formation. The formation of methionine and 5’-dAH (**Fig. S3**) with various reductants mimics PChlide formation. Together, these data confirm that purified *Rp*BchE is necessary and sufficient to catalyze the 6 e-oxidation of MPE to PChlide *in vitro*.

Cbl-dependent RS methylases can access cob(I)alamin, which performs a nucleophilic attack on the methyl group of SAM, resulting in the formation of MeCbl and SAH. Though BchE is not a methylase, we also monitored the production of MeCbl and SAH to determine if using chemical reductants could result in aberrant chemistry that inhibits the native reaction. When using Ti(III)citrate as the reductant, MeCbl production is prevalent, accounting for ∼60% of the total Cbl concentration (**Figure 3E**). DT and the MV/FMN/NADPH reducing system support negligible MeCbl accumulation. This trend is also reflected in the production of SAH (see **Fig. S3**). Ti(III)citrate is commonly used for *in vitro* studies of Cbl-dependent RS enzymes due to its ability to reduce both the [Fe_4_S_4_] cluster and Cbl to their +1 oxidation states. While it is not surprising that Ti(III)citrate supports more MeCbl production as compared to the other “weaker” reductants, it is perplexing that Ti(III)citrate also supports the highest production of PChlide, given that the substrate is not observed to undergo methylation and, therefore, MeCbl is not expected to be an intermediate.

Given that BchE is not a methylase, we predicted that MeCbl might be an inactive form of the Cbl cofactor in the BchE reaction. While optimizing reaction conditions, we found that incubating BchE with SAM and Ti(III)citrate before initiating the reaction by adding MPE results in much lower activity compared to when initiating with SAM or Ti(III)citrate (**Fig. S4**). We attributed the lower activity to MeCbl formation; however, reaction optimization was not performed with the goal of detecting MeCbl. Therefore, another assay was performed in the dark, where BchE was incubated with SAM and Ti(III)citrate to premethylate Cbl before initiating the reaction with MPE. This order of addition was compared to incubating with SAM and MPE and then initiating with Ti(III)citrate. **Figure 4A** shows that the final concentration of MeCbl in the premethylated reaction is near stoichiometric with total Cbl (10 µM) and remains constant throughout the reaction. By contrast, initiating with reductant builds up MeCbl to ∼55% of the total Cbl. The SAH concentration is marginally higher but still correlates with MeCbl (**Fig. S5**). When Cbl is premethylated, PChlide formation is greatly inhibited (**Figure 4B**), as is the formation of methionine and 5’-dAH (**Fig. S5**). When reactions are initiated with Ti(III)citrate, PChlide accumulates to ∼40% of a full turnover, while MeCbl accumulates to ∼55% of the total enzyme. These data suggest that MeCbl is not the active form of the Cbl cofactor and that the reaction requires that Cbl not be methylated.

**Figure 4.**
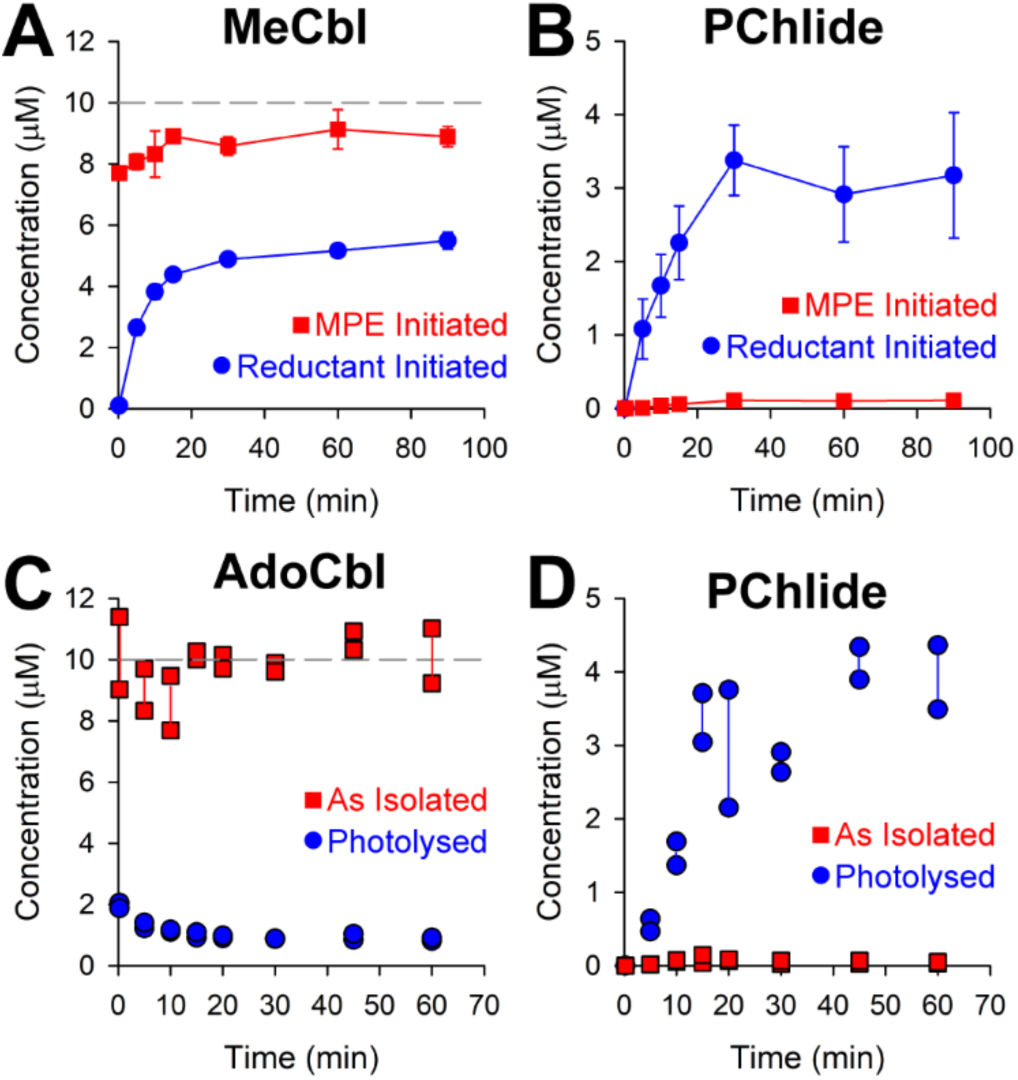
BchE assays comparing MeCbl (**A**) and PChlide (**B**) formation when initiated with substrate, MPE, or reductant. BchE assays using protein purified with AdoCbl (as isolated) compared to photolyzing the protein with 385-395 nm UV light before initiating the reaction. AdoCbl content is shown in **C**) and PChlide formation is shown in **D**). Premethylation reactions were run in triplicate and AdoCbl reactions were run in duplicate. The gray dashed line indicates enzyme concentration.

Upon discovering the Cbl dependence of the BchE reaction, Gough et al. first proposed a mechanism with AdoCbl as the active cofactor and the source of the 5’-dA•. Since BchE’s annotation as an RS enzyme, other mechanisms have been proposed with SAM serving as the source of the 5’-dA•.(16, 17) However, AdoCbl has never been excluded as an active cofactor. Our current preparations of BchE are derived from an *E. coli* strain unable to produce AdoCbl, and only OHCbl is added during expression and purification. Furthermore, no AdoCbl is detected by LC-MS after denaturation of BchE. These observations strongly suggest that AdoCbl is not needed for BchE catalysis. Nevertheless, given the concern that AdoCbl might be present at trace levels in BchE preparations, additional experiments were conducted using BchE purified from *E. coli* BL21(DE3) (i.e., not the *ΔbtuR* knockout), in which AdoCbl is the predominant form of Cbl in the isolated enzyme. This enzyme was used in assays initiated with Ti(III)citrate to determine if AdoCbl enhances BchE activity. One assay was conducted with the AdoCbl-bound enzyme. In a second assay, AdoCbl bound to BchE was photolyzed to cob(II)alamin for 45 min on ice with 385-395 nm UV light. As shown in **Figure 4C**, AdoCbl accounts for almost all Cbl forms throughout the reaction. Upon photolysis, AdoCbl constitutes 20% of the Cbl forms, which further drops to ∼10% as the BchE reaction progresses. The effect of photolysis on activity is substantial, as seen in **Figure 4D**. PChlide formation is greatly enhanced when there is less AdoCbl, signifying that AdoCbl is not the active cofactor. These findings are replicated with methionine and 5’-dAH formation (**Fig. S6**).

### Activity with Ferredoxins

Among the tested reductants, Ti(III)citrate yields the most product and promotes the formation of MeCbl, which we show is inhibitory. Ti(III)citrate’s lower reduction potential likely affords more efficient reduction of the FeS cluster for the reductive cleavage of SAM and the reduction of Cbl to cob(I)alamin, leading to methyl transfer from SAM to Cbl. We surmise that these are competing reactions, in which PChlide forms faster than MeCbl. Given that Ti(III)citrate is both nonideal and nonphysiological as a reducing system for the BchE reaction, biological reductants that are more native to BchE’s cellular environment were tested (See Supporting Information for a full list). We found two ferredoxin systems that were able to support turnover by BchE: FdxA from *R. capsulatus* (Uniprot D5AP15) and a ferredoxin from *R. gelatinosus* (Uniprot I0HRP6). *Rc*FdxA is a [Fe_7_S_8_] ferredoxin containing both [Fe_3_S_4_] and [Fe_4_S_4_] clusters. The [Fe_4_S_4_] cluster has been shown to have a low reduction potential.(28) The ferredoxin from *R. gelatinosus* is also annotated as a potential [Fe_7_S_8_] ferredoxin and exhibits 51% sequence identity to *Rc*FdxA. Both proteins were overexpressed in *E. coli* BL21(DE3) along with the genes on plasmid pDB1282 prior to anaerobic purification by Ni-IMAC. The EPR spectra for both as-isolated proteins show signals consistent with a [Fe_3_S_4_] cluster (**Fig. S7**). However, neither protein shows the emergence of an EPR spectrum consistent with a [Fe_4_S_4_]^+^ cluster upon reduction with dithionite, only the attenuation of the signal associated with the [Fe_3_S_4_] cluster. In previous studies of this protein, a similar behavior was reported. Reduction of the [Fe_4_S_4_] cluster to the +1 oxidation state required photoreduction with 5-deazaflavin.(28)

Activity assays were conducted using BchE isolated from the *ΔbtuR* strain of *E. coli* with either *Rc*FdxA or *Rg*FdxA, along with a purified ferredoxin reductase (Uniprot I0HR46) and 2 mM NADPH. Compared to assays using Ti(III)citrate, both ferredoxins support a slower formation of PChlide, as shown in **Figure 5A**. The reaction in the presence of Ti(III)citrate plateaus after 1 h, allowing the reaction in the presence of *Rc*FdxA to ultimately reach the same quantity of PChlide after 4 h. Notably, *Rg*FdxA outperforms these two reductants, allowing BchE to undergo multiple turnovers. The formation of methionine (**Figure 5B**) and 5’-dAH (**Fig. S8**) is greater in the reaction with Ti(III)citrate, but less product is formed. This behavior is likely due to increased abortive cleavage of SAM. Furthermore, MeCbl and SAH formation is negligible for both the ferredoxin systems (**Fig. S8**). The lack of MeCbl formation suggests that the ferredoxins do not readily reduce Cbl to cob(I)alamin. The quantification of both SAH and 5’-dAH for the *Rc*FdxA reaction is systematically low. We suspected this is due to an SAH nucleosidase contamination from the *Rc*FdxA purification. The *Rc*FdxA reaction shows a spike in adenine concentration at early time points, followed by a decline (**Fig. S9**). This decline is also correlated with an accumulation of hypoxanthine, suggesting the presence of both SAH nucleosidase and adenine deaminase. The Ti(III)citrate and *Rg*FdxA reactions both show linear adenine formation at an initial rate that is ∼3 times slower. Hypoxanthine formation is also negligible in these reactions.

**Figure 5.**
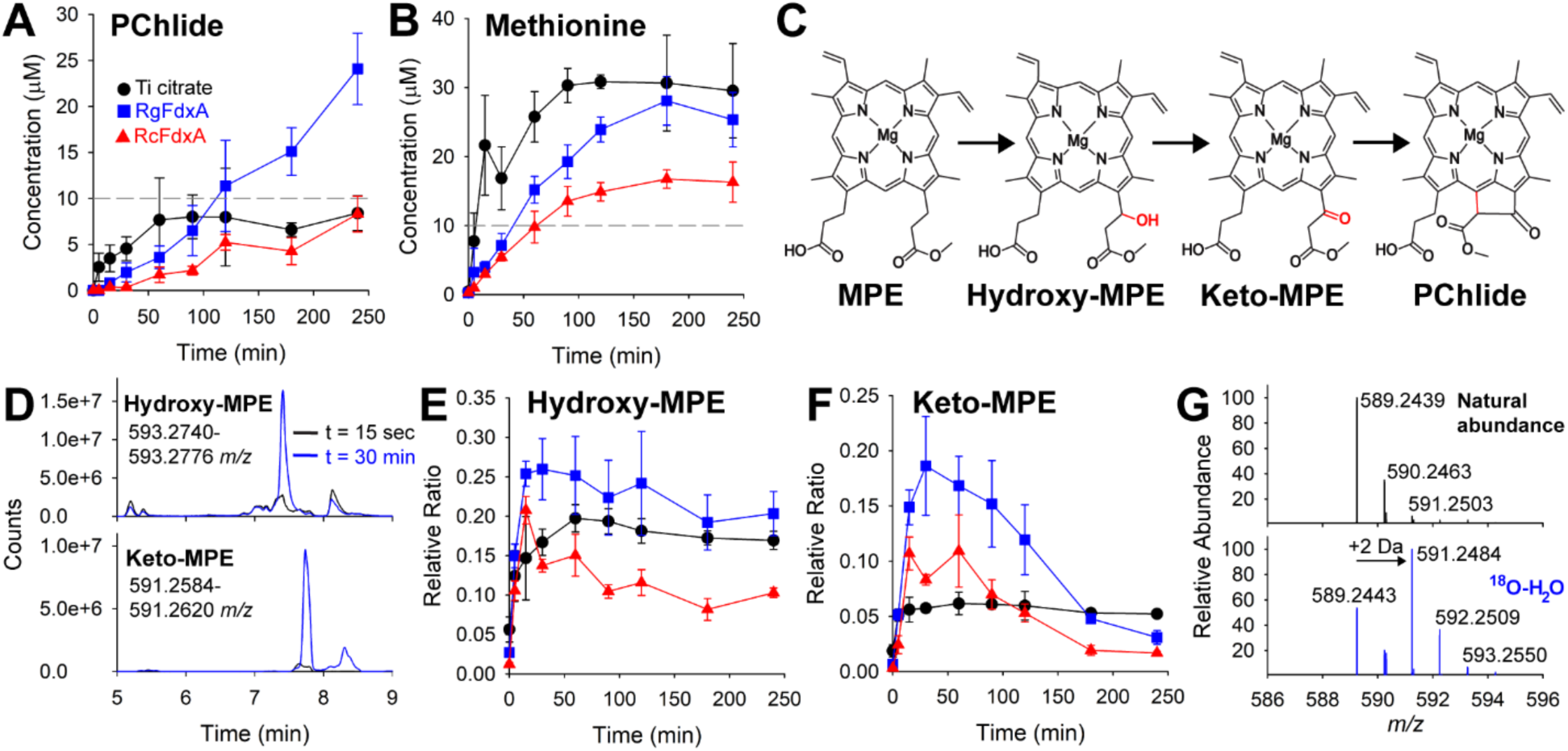
BchE assays utilizing ferredoxins *Rg*FdxA and *Rc*FdxA compared to Ti(III)citrate as reducing systems. PChlide (**A**) and methionine (**B**) formation are shown for these reactions. The gray dashed line indicates enzyme concentration. **C**) shows the scheme of proposed intermediates for the BchE reaction. The extracted ion chromatograms for proposed intermediates, hydroxy-MPE and keto-MPE are shown in **D**) from quenched BchE reactions using *Rg*FdxA at t = 15 sec and 30 min. Formation of Hydroxy-MPE (**E**) and Keto-MPE (**F**) with time is shown for the BchE reactions shown in **A**) and **B**). **G**) displays the mass spectrum of PChlide produced from BchE reactions performed anaerobically in buffer made with natural abundance versus ^18^O enriched water. Error bars display standard deviation from reactions run in triplicate.

### Detection of proposed intermediates

Multiple mechanisms have been proposed for the BchE reaction, all of which invoke the order of intermediates shown in **Figure 5C**. Hydroxy-MPE is formed by oxidation of the 13^1^-carbon, followed by a further oxidation to keto-MPE. Lastly, cyclization yields PChlide by forming a bond between carbons 13^2^ and 15. Chen et al. detected the hydroxy and keto intermediates for the paralogous aerobic enzyme, AcsF.(9) However, to our knowledge, these intermediates have not been reported for the BchE reaction. **Figure 5D** shows extracted ion chromatograms from t = 30 min of the BchE reaction utilizing *Rg*FdxA as reductant. The top (593.2758 *m/z)* and bottom (591.2602 *m/z*) traces correlate to the [M+H]^+^ for hydroxy-MPE and keto-MPE, respectively (both without bound magnesium). The MS/MS fragmentations of both hydroxy-MPE and keto-MPE are consistent with those shown by Chen et al. for AcsF (**Fig S10&11**).(9) The time-dependent formation of both species is shown in **Figure 5E&F** with Ti(III)citrate and the ferredoxin reductant systems. Quantification of these intermediates was not possible due to the lack of standards. Instead, formation is represented by the response ratio (e.g., area of hydroxy-MPE/area of internal std). In each system, hydroxy-MPE accumulates at early timepoints. Only the *Rc*FdxA reaction shows a clear decrease in abundance of this species at longer times. By contrast, the formation and decay of keto-MPE is dramatic with both ferredoxin systems, consistent with it being an intermediate. **Fig. S12** shows an overlay of normalized PChlide, hydroxy-MPE, and keto-MPE formation for both of the BchE reactions with ferredoxin reducing systems. In both cases, PChlide accumulation clearly correlates with keto-MPE attenuation. The hydroxy- and keto-MPE species are also observed when using dithionite and MV-FMN-NADPH chemical reductants (**Fig. S13**). Interestingly, while the MV-FMN-NADPH reducing system affords the lowest PChlide formation, it produces the highest accumulation of the proposed intermediates. This behavior suggests that the MV-FMN-NADPH reducing system is adequate for initiating the BchE reaction but struggles to reduce the system for the final cyclization. Given that this behavior is observed only for the weakest reducing system, we interpret it as the enzyme having a lower reduction potential for the last step, perhaps requiring reduction to cob(I)alamin (*vide infra*).

Both intermediates appear to plateau without a decay phase when using Ti(III)citrate, consistent with Ti(III)citrate supporting the generation of a MeCbl species, which inhibits the reaction. This is also supported by the observation that the final PChlide concentration is consistently lower than the enzyme concentration when using Ti(III)citrate. The initial rate of formation of the hydroxy-MPE and keto-MPE intermediates appears comparable between the Ti(III)citrate and ferredoxin reducing systems. However, formation of PChlide is slower with the ferredoxin reducing systems compared to Ti(III)citrate. These discrepancies in rates suggest that the enzyme must be reduced by an external electron at least two times: once to produce the hydroxy-MPE intermediate and once to produce PChlide. It remains to be seen if BchE requires an external electron to form keto-MPE or if this step is similar to that proposed for BtrN, where the auxiliary FeS cluster can transfer an electron back to the RS cluster.(29, 30) In the case of BchE, the role of the auxiliary cluster would be replaced by Cbl.

As shown above, methylating the Cbl of BchE by incubating with Ti(III)citrate and SAM results in a drastic drop in PChlide production (∼30× less than when initiating with reductant after incubation with SAM and MPE). In these conditions, the accumulation of hydroxy-MPE and keto-MPE only decreased by approximately 2× and 3×, respectively (**Fig. S14**). This observation suggests that cobalamin is required for the final cyclization, whereas hydroxy-MPE and keto-MPE formation are independent of the state of Cbl but influenced by the steric hindrance of the methyl ligand.

### Origin of oxygen in the BchE reaction

AcsF converts MPE to PChlide utilizing a diiron active site, with the oxygen atoms in the hydroxy and keto intermediates stemming from O_2_. For BchE, both *in vivo* and *in vitro* studies have shown that water is the source of the oxygen atom in the 13^1^ ketone of PChlide.(22, 23, 31) RS enzymes are notoriously sensitive to O_2_, which degrades their [Fe_4_S_4_] clusters, suggesting that O_2_ is unlikely to serve as a substrate in the BchE reaction. However, DarE, a RS enzyme that catalyzes a key step in the biosynthesis of darobactin, requires O_2_ as a substrate.(32) To show that O_2_ inhibits the BchE reaction, we performed reactions outside an anaerobic chamber rather than inside, as in all previous BchE reactions in this study. As shown in **Fig. S15,** O_2_ exposure severely diminishes BchE activity, consistent with the study of Wiesselmann et al.(23) A second assay was conducted anaerobically in the presence of ∼70% H_2_^18^O. PChlide features a carboxylate, a methyl ester, and the newly formed ketone group as oxygen-containing moieties. To ensure that ^18^O incorporation depends on BchE activity and does not result from oxygen exchange in the acidic quench solution, the reaction was quenched with methanol after 30 min and immediately frozen in liquid N_2_. The sample was completely dried using a SpeedVac concentrator and resuspended in a 50/50 solution of 150 mM H_2_SO_4_/isopropanol (all at natural abundance) before analysis by HRMS. The resulting mass spectrum of PChlide from the reaction conducted in ∼70% H_2_^18^O is shown in **Figure 5G** and compared to PChlide dissolved in buffer containing H_2_O at natural abundance. A shift of +2 *m/z* is observed, denoting the presence of ^18^O. The mass spectra of hydroxy-MPE and keto-MPE also show similar shifts (**Fig. S16**). Notably, the +2 *m/z* shifted signal observed for hydroxy-MPE appears at a lower-than-expected intensity relative to natural abundance, especially when compared with the corresponding signals for PChlide and keto-MPE. It is possible that the first oxidation step, which forms hydroxy-MPE, uses a water molecule that is already tightly bound to the active site via hydrogen bonding or bound to cobalamin. Subsequent steps will then allow exchange with the solvent and increase the likelihood of ^18^O incorporation. MPE shows no shift, indicating that ^18^O incorporation into PChlide does not stem from oxygen exchange into either the carboxylate or ester (**Fig. S16**).

### Characterization of an unsaturated species

The chromatogram of a quenched BchE reaction (**Figure 3A**, *trace II*) shows the formation of the final product, PChlide, as well as another species exhibiting a retention time (RT) of 8.08 min. The mass spectrum corresponding to this peak (termed the *575* species) is shown in **Figure 6A** and displays an [M+H]^+^ of 575.2652 *m/z*. We observe time-dependent formation of the *575* species when using chemical reductants (**Fig. S17**) and both ferredoxin-reducing systems (**Figure 6B**). Formation of the *575* species shows the same MPE, SAM, and enzyme dependence as does PChlide formation (**Figure 3A**). We initially suspected that the *575* species is an intermediate in the reaction. Yokoyama and Lilla proposed a mechanism for BchE featuring an olefin intermediate, which agrees with the observed mass of the *575* species.(17) This mechanism had precedent from other RS enzymes that generated olefin products, but there are also examples of olefin shunt products in RS reactions.(29, 32–34) Therefore, we set out to test the chemical and kinetic competence of the *575* species. A BchE reaction was scaled up to 10 mL and quenched with 10 mL of 0.1% ammonium hydroxide in methanol. Preparative HPLC was then used to isolate the *575* species using a pH-neutral buffer to retain the central Mg ion. Fractions containing the *575* species were dried down and desalted. The isolated *575* species was incubated with BchE under typical reaction conditions using either Ti(III)citrate or the *Rg*FdxA reducing system. Quenched reactions were analyzed by HRMS, but ultimately, no PChlide or intermediate formation was observed. These results suggest that the *575* species is likely not an intermediate, but an off-pathway product. It is not surprising that an off-pathway product would accumulate in the presence of chemical reductants such as Ti(III)citrate. The overall reaction is proposed to involve multiple steps, with the oxidation state of the cofactors likely playing a critical role. Having excess reductant could disrupt the active cofactor oxidation state at any given step. Interestingly, the formation of the *575* species with the ferredoxin reducing is comparable to that with Ti(III)citrate (**Figure 6B**). We anticipated that biological reductants would yield more “controlled” enzymatic reactions, because they are expected to be more selective in the cofactors they reduce.

**Figure 6.**
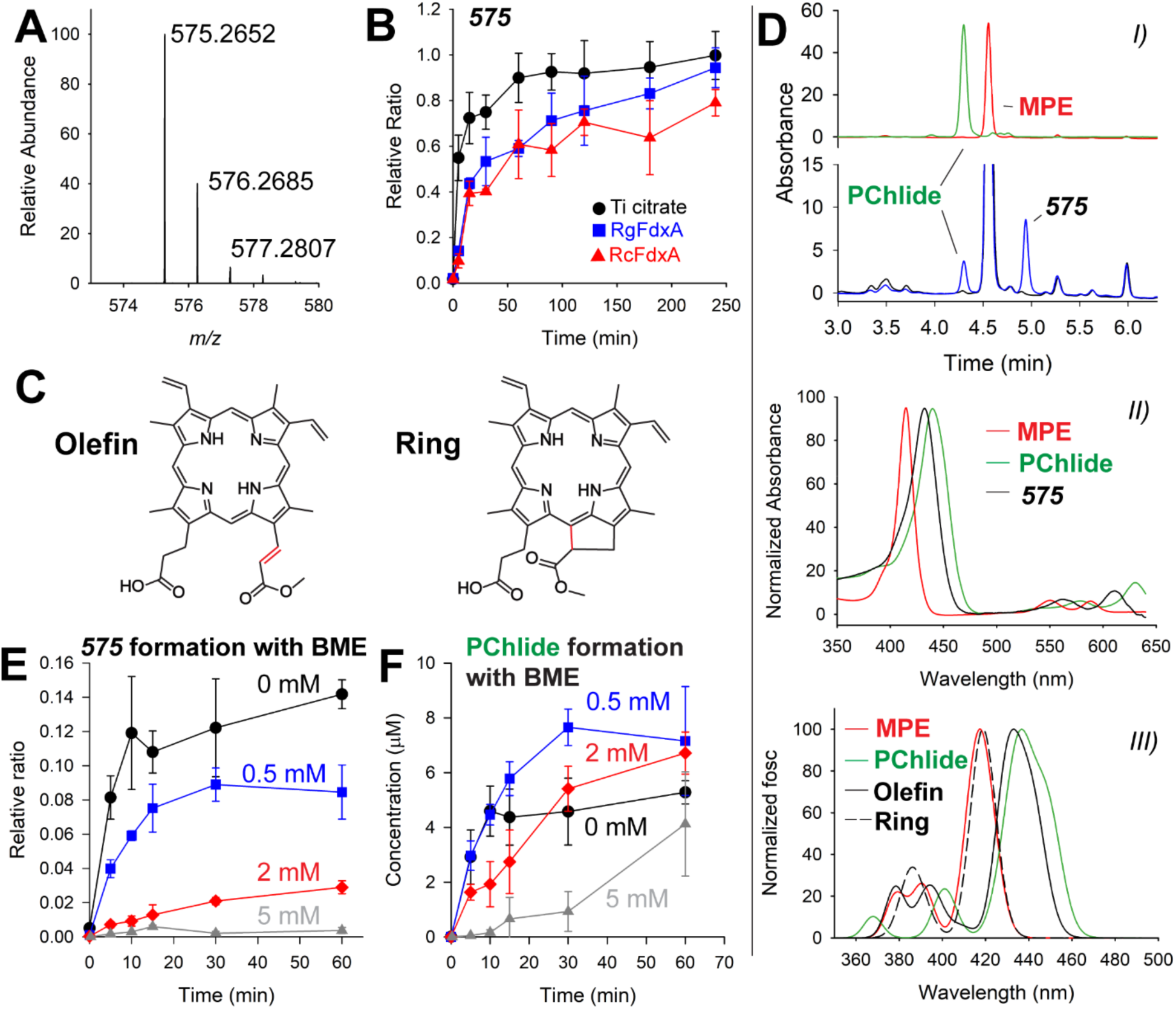
Mass spectrum of the observed *575* species (**A**) and formation (**B**) with time using Ti(III)citrate, *Rg*FdxA, and *Rc*FdxA reducing systems. **C**) shows the two proposed structures of *575*. **D**) details the UV-vis characterization of *575*. The HPLC chromatogram *I)* shows time-dependent formation of the *575* species compared to standards of MPE and PChlide. The UV-vis spectra from each species is shown in II). TD-DFT-calculated spectra are shown in *III)* for the optimized structures of MPE, PChlide, olefin-*575*, and ring-*575*. E) and F) show the effects of BME on the production of *575* and PChlide, respectively. BME was added at concentrations of 0 mM (*black circles*), 0.5 mM (*blue squares*), 2 mM (*red diamonds*), and 5 mM (*gray triangles*). B, E, and F error bars represent the standard deviation from triplicate reactions.

Although the *575* species is most likely an off-pathway product, its structure may contain relevant insight into the BchE mechanism. The mass correlates with that of the substrate, MPE, with the loss of two protons, most likely through one degree of unsaturation. Considering that the porphyrin substrate is already highly unsaturated, we suggest two possibilities. The first is that an olefin is formed, such as that proposed by Yokoyama and Lilla (**Figure 6C**, Olefin). The second is that cyclization occurs before oxygen addition, affording the product without oxygen incorporation (**Figure 6C**, Ring).

When isolating the *575* species to test for chemical and kinetic competence, we were unable to obtain sufficient yields and/or purities for NMR analysis. MS/MS fragmentation was performed by HRMS, but it was unable to exclude either the olefin or the cyclized product (**Fig. S18**). The *575* species was also generated from a BchE reaction containing MPE with a deuterated (*d_3_*) methyl group on the ester moiety. The *d_3_*-containing *575* species was also fragmented by MS/MS to compare with the natural abundance *575* species. This data suggests that the unsaturation is occurring on the ester chain of MPE rather than the carboxylate chain. A detailed analysis leading to this conclusion is provided in *Supporting Information*.

UV-vis spectroscopy was used to further characterize the *575* species. MPE and PChlide show distinct UV-vis features in their Soret and Q-bands. Removing the magnesium under acidic conditions affects the UV-vis features. Therefore, a BchE reaction was quenched with basic 0.2% ammonia in methanol to retain the magnesium in the porphyrins. Spectra were recorded via HPLC with a photodiode array (PDA) detector using a gradient with pH-neutral (typically pH 6.9) aqueous (40 mM ammonium acetate) and methanol mobile phases. The resulting chromatogram is shown in **Figure 6D**, *I*, with MPE and PChlide standards for comparison. The peak corresponding to the *575* species was verified by HRMS (**Fig. S19**). The UV-vis spectra for MPE, PChlide and the *575* species are shown in **Figure 6D**, *II*. The spectra for MPE and PChlide agree with those previously recorded.(35–37) Notably, the shifts seen when comparing MPE to PChlide for the Soret bands are 415 nm to 440 nm, while the Q-bands shift from 550 and 588 nm to 578 and 630 nm. The spectrum of the *575* species is also red-shifted from that of MPE, but to a lesser extent, with a Soret band at 432 nm and Q-bands at 562 and 610 nm.

Although we could not distinguish between the olefin and cyclized structures for the *575* species, we suspected that the two would have markedly different UV-vis features. Time-dependent DFT (TD-DFT) was used to investigate the predicted spectral differences between the olefin and cyclized *575* species structures as well as how they compare to the calculated spectra for MPE and PChlide. Orca 5.0 was used to optimize structures of MPE, PChlide, and the olefin-*575* and cyclized-*575* species.(38, 39) Bacteriochlorophyll and related porphyrins are known to prefer five-coordinate Mg geometries.(40, 41) The mass spectra of MPE, PChlide, and the *575* species show an acetic acid adduct as the most prevalent species for the HPLC method used to obtain UV-vis spectra. Therefore, we retained a five-coordinate magnesium geometry by coordinating an acetic acid carbonyl to the magnesium in each structure. Optimized coordinates for each species are given in the *Supporting Information*. TD-DFT was used to generate theoretical absorption spectra for each species. After including a shift of −0.19 eV to each transition energy, the Soret bands of MPE and PChlide accurately reproduce the experimental values, with λ_max_ values at 417 and 437 nm, respectively. These two matching data points help validate the computational method’s ability to predict the UV-vis features of these porphyrins. The predicted absorption spectra for the ring and olefin structures are overlaid in **Figure 6D**, *III*. The spectrum corresponding to the ring shows minimal change from that of MPE, with a λ_max_ of 419 nm. The olefin structure shows a predicted λ_max_ of 433 nm, matching the experimental value for the *575* species. The ring structure features no changes in conjugation compared to MPE, consistent with the negligible change in calculated absorbance. Conversely, the olefin extends conjugation from the corrin ring to the C^13^ ester group, which qualitatively explains the lower energy excitations similar to PChlide. Similar trends are shown for the calculated Q-bands (**Fig. S20**).

Assuming that the structure of the *575* species is an α,β-unsaturated ester, it should be susceptible to Michael addition. BME was used as a Michael donor to determine if the thiol would influence the production of the *575* species and overall PChlide formation. Assays were performed by adding BME to the BchE reaction during incubation, prior to initiation with Ti(III)citrate. BchE was run through a prepacked G-25 resin column to remove residual BME from the purification before adding 0, 0.5, 2, or 5 mM BME. **Figure 6E** shows that the presence of BME greatly inhibits the formation of the *575* species. The formation of the *575* species is reduced by almost 50% with only 0.5 mM BME. 2 mM BME shows further attenuation to ∼20%, while 5 mM BME essentially removes all of the *575* species. Conversely, PChlide formation is aided by adding 0.5 mM BME. **Figure 6F** shows that reactions lacking BME plateau in product formation at 10 min, while 0.5 mM BME extends the reaction to 30 min. Higher concentrations of BME result in slower PChlide formation but ultimately yield similar endpoint yields. The expected product of Michael addition between the olefin-*575* species and BME is shown in **Fig. S21**. An m/z signal corresponding to this species can be detected by HRMS. However, it appears to be unstable, as it decays after initial formation (**Fig. S21**).

Reactions were also performed with corresponding concentrations of TCEP instead of BME. TCEP is a reductant frequently used in place of DTT or BME but lacks a thiol group for Michael addition. Unlike BME, incubation with TCEP does not result in a drastic inhibition of the formation of the *575* species relative to PChlide (**Fig. S22**). Increasing TCEP concentrations had either a positive or no effect on the formation of the *575* species and PChlide until 5 mM, at which point both were inhibited to a similar degree. From this data, we conclude that while excessive reductant can be detrimental to the overall reaction, the *575* species is selectively sensitive to thiols. Both its inhibition by BME and its UV-vis spectrum, validated by TD-DFT, support the olefin structure. Therefore, we feel confident assigning the *575* species as an olefin.

In our attempts to obtain soluble BchE constructs, we also purified BchE from *R. capsulatus* and *R. gelatinosus*. Neither of these enzymes produced the PChlide product and could only form the *575* species using Ti(III)citrate with no activity using ferredoxins. We first speculated that the *575* species was an intermediate and that BchE required a partner protein to complete its reaction, as observed with OxsB/OxsA.(10) This hypothesis was ultimately wrong given our finding that *Rp*BchE is fully active alone, and that the *575* species is not an intermediate in the reaction. Nevertheless, we replicated an experiment from Gough et al.’s seminal work that showed BchE’s Cbl dependence.(20) For this experiment, freeze-thawed cells of *R. capsulatus* were incubated with MPE, nicotinamide (NAM), and MgCl_2_ to induce the cell’s native BchE activity. *R. capsulatus* does not utilize AcsF, so all production of PChlide must originate through the action of BchE. To replicate the results of Gough et al., we compared *R. capsulatus* cells grown aerobically in the dark with those grown anaerobically in the presence of light. Fluorescence spectroscopy showed that cells grown anaerobically preferentially accumulated a species distinct from PChlide. As with Gough et al., aerobically grown cells showed that PChlide (emission 633 nm) formation was dependent on both MPE and NAM (**Fig. S23**). The anaerobically grown cells showed a different species with an emission peak at 620 nm. A similar signal was observed during preliminary assays of *Rg*BchE, which could only produce the olefin off-pathway product. Analysis of cell extracts by HRMS not only detected the olefin species but also showed elevated levels in anaerobically grown samples, consistent with trends observed by fluorescence. More details are provided in the *Supporting Information*, but the key takeaway is that the olefin species is produced *in vivo* in *R. capsulatus*, especially when grown anaerobically.

## Discussion

In this work, we report the successful reconstitution of the in vitro activity of purified BchE. Although BchE has been annotated as a Cbl-dependent RS enzyme for over two decades, mechanistic studies have been severely limited by its notorious insolubility.(22, 23) To address this challenge, we attempted heterologous expression and purification of BchE homologs from eight different organisms; only three yielded any soluble protein, and of these, only BchE from *Rubrivivax pictum* catalyzed the formation of PChlide. Protein yield and stability were substantially improved by the introduction of a C-terminal MBP tag in combination with optimized expression and purification conditions.

Only a small number of known Cbl-dependent RS enzymes do not catalyze methylation. In addition to BchE, these include OxsB, HpnJ, PbsB, and ladderane synthase, with OxsB being the only member of this group to be isolated and characterized mechanistically.(4, 10, 12–14, 42) These enzymes act on chemically diverse substrates, including peptides, hopanoids, nucleotides, lipids, and porphyrins, but all catalyze reactions involving ring formation or a rearrangement. For each case, the precise role of Cbl remains speculative. Mechanistic studies of OxsB have led to a proposal in which bond formation between two *sp*³-hybridized carbons proceeds via the generation of a substrate radical that is stabilized by cob(II)alamin, forming a putative Co–alkyl intermediate.(43, 44) This stabilization would enable a second H• abstraction, generating a new carbon-centered radical and ultimately leading to C–C bond formation. However, the proposed Co-alkyl intermediate has not been directly observed. BchE’s successful reconstitution now provides an additional system to interrogate the mechanistic roles of Cbl in nonmethylase Cbl-RS enzymes.

Multiple reducing systems supported product formation by purified BchE. In addition to the proposed hydroxy- and keto-MPE intermediates, we detected unexpected species, including MeCbl and a previously uncharacterized compound absorbing at *575* nm. Because BchE contains multiple redox-active cofactors, we could not rule out the possibility that the *575* nm species arose from indiscriminate reduction of these cofactors by small-molecule reductants, leading to off-pathway or shunt products.(32) Moreover, the catalytically relevant oxidation state of Cbl in the BchE reaction remained unclear. To better approximate physiological conditions and preserve native redox states, we also tested biological reducing systems. Two ferredoxins were found to support BchE activity. Notably, both *Rc*FdxA and *Rg*FdxA are known to contain an [Fe_7_S_8_] cluster. Ferredoxins of this class have been reported to exhibit lower reduction potentials than [Fe₄S₄] or [Fe₂S₂] single cluster-containing ferredoxins.(45) Using these ferredoxins in BchE reactions, the proposed hydroxy- and keto-MPE intermediates accumulated and subsequently decayed in concert with PChlide formation, consistent with their assignment as intermediate species. In addition, MeCbl formation was negligible in reactions containing protein reductants as compared to reactions containing chemical reductants. *Rc*FdxA was originally identified by Wiesselmann et al. as a potential low-molecular-weight partner in the BchE reaction, although those authors were unable to reconstitute activity using isolated *Rc*FdxA and natively expressed BchE from *R. capsulatus*.(23) Similarly, despite our ability to purify soluble *Rc*BchE, we did not observe PChlide formation with *Rc*FdxA. Thus, it remains unclear why the *Ap*BchE construct was fully active, whereas the *Rg* and *Rc* homologs predominantly produced the *575* species.

The off-pathway *575* species was a major byproduct in the *in vitro* reactions described here. Based on spectroscopic analysis, we assign this compound as an olefin, consistent with an intermediate previously proposed by Yokoyama and Lilla. Importantly, the *575* species was observed both *in vitro* using chemical or biological reducing systems and *in vivo* in *R. capsulatus* cells. The isolated *575* species was not transformed into PChlide by BchE, suggesting that this olefin is a shunt product. A similar olefin off-pathway product has been reported for DarE, another enzyme that employs multiple redox cofactors.(32) To our knowledge, the *575* species has not been previously reported for BchE.

As noted above, the initial mechanism proposed by Gough et al. predates the identification of BchE as an RS enzyme and the exclusion of AdoCbl as a catalytically relevant cofactor, as demonstrated herein. With that mechanism eliminated, two prominent mechanistic models from the literature remain (Figure 7). Path A, proposed by Booker, begins with H• abstraction from carbon 13^1^ of MPE to generate a substrate-centered radical. This radical is subsequently quenched by the transfer of a hydroxyl group from OHCbl to form hydroxy-MPE. Although mechanistically analogous to methyl transfer in Cbl-dependent RS methylases, a hydroxyl-transfer role for OHCbl has not been reported in biological systems. Repetition of this process would generate a geminal diol, which could eliminate water to form the keto-MPE intermediate.

**Figure 7.**
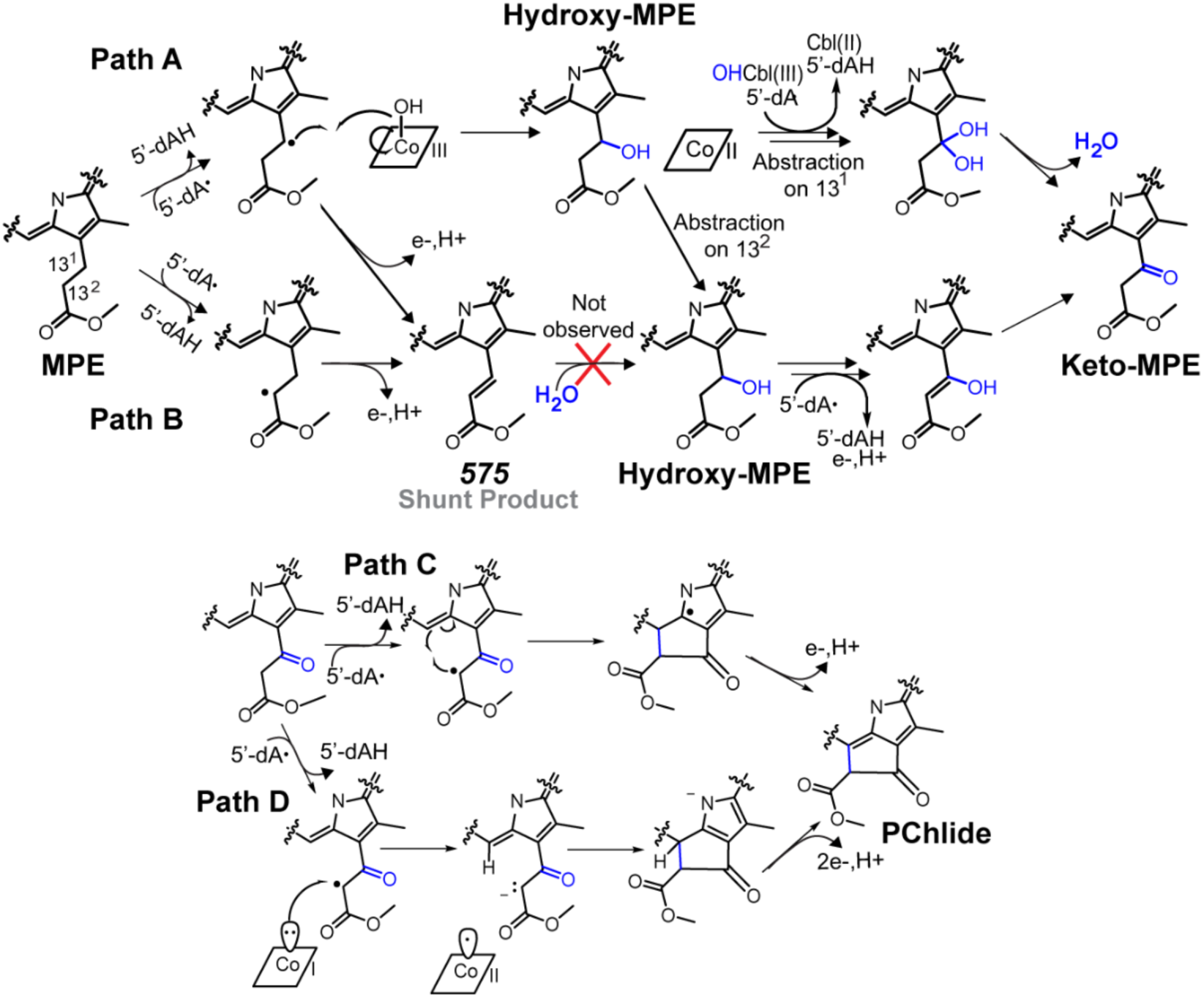
Proposed mechanisms for the BchE reaction.(16, 17)

Path B, proposed by Yokoyama and Lilla, diverges from Path A by initiating H• abstraction at carbon 13^2^.(17) Subsequent loss of an electron and a proton would yield an olefinic intermediate. In this work, we observed the accumulation of such a species, designated *575*. In principle, this α,β-unsaturated ester could undergo 1,4-addition of water to produce hydroxy-MPE. However, we were unable to detect the transformation of isolated *575* to PChlide, hydroxy-MPE, or keto-MPE. One possibility is that catalysis requires the enzyme to adopt a distinct state, such as a specific cofactor oxidation state or active site conformation, which is not fully accessed when starting with the *575* species as a substrate.

Another possibility is that a specific isomer (*i.e. E* or *Z*) of the *575* olefin is needed for the reaction to progress forward. In such a case, the *575* species we isolated would be the incorrect isomer and incapable of acting as a substrate for the BchE reaction. Unfortunately, we currently lack NMR or other characterization of the *575* species to indicate which isomer accumulates. Notably, formation of the *575* species could occur regardless of whether the initial H• abstraction takes place at carbon 13^1^ or carbon 13^2^ (Figure 7). An appealing feature of Path B + Path C/D is that all H• abstractions occur at a single position, whereas in Path A + Path C/D, two abstractions at carbon 13^1^ followed by an abstraction at carbon 13^2^ are required to proceed into Path C/D. Alternatively, the first H• abstraction may occur at carbon 13^1^ to yield hydroxy-MPE, as in Path A, followed by abstraction at carbon 13^2^ to converge with Path B. In this scenario, formation of the alcohol could induce a conformational change that repositions carbon 13^2^ closer to the RS cluster.

Following keto-MPE formation, both proposed mechanisms converge on a final H• abstraction at carbon 13^2^ (Path C) to generate a carbon-centered radical that attacks carbon 15, forming a five-membered ring. Aromaticity would then be restored through the loss of a proton and an electron, yielding PChlide. An alternative non-radical cyclization mechanism is depicted in Path D. In this model, the hydrogens at carbon 13^2^ of keto-MPE—being β to both a methyl ester and a ketone—are predicted to have pK_a_ values of 9–13. Following H• abstraction, cob(I)alamin could donate an electron to the radical, forming a carbanion (Path D) stabilized by the adjacent ketone and methyl ester functional groups. This enolate would then attack the C15 position to form the fifth ring. Subsequent loss of two electrons and a proton would yield PChlide. Zhu and Silverman showed that the *meso*-carbons of protoporphyrin IX become more electrophilic when the vinyl moieties are replaced with electron-withdrawing acetal groups.(46, 47) Therefore, the ketone of keto-MPE may produce a similar effect to facilitate nucleophilic attack on C15.

Support for a pathway involving cob(I)alamin is consistent with our observation that hydroxy- and keto-MPE intermediates accumulate when using the weakest reducing system (MV– FMN–NADPH), which may be insufficient to reduce cobalamin to the cob(I)alamin state. This interpretation assumes that the Cbl (II/I) redox couple is lower than that of the +2/+1 redox couple of the RS-cluster, which is generally the case but has not yet been established for BchE.(26) Furthermore, our EPR data suggest the Cbl in BchE is four-coordinate, which would stabilize cob(I)alamin.

In summary, we have purified and characterized BchE in vitro for the first time and established that it generates hydroxy- and keto-MPE intermediates, as well as the PChlide product, using both chemical and protein-based reducing systems. Under the low-potential conditions provided by Ti(III)citrate, we observe substantial MeCbl formation, which suppresses overall turnover but does not prevent accumulation of the intermediates, suggesting that OHCbl is unlikely to serve as the direct hydroxylating species. The appearance of a *575* species in both in vivo and in vitro contexts raises the possibility of an olefinic intermediate, although the isolated species is not converted to product when reintroduced into a reaction. Together, these findings establish a robust experimental platform for dissecting this unusual transformation and defining the mechanistic role of BchE in greater detail.

## ASSOCIATED CONTENT

### Supporting Information

Materials and Methods, Table S1, supporting figures (S1-S23), protein, peptide, and DNA sequences, and optimized DFT structure coordinates. The following files are available free of charge.

## AUTHOR INFORMATION

### Author Contributions

The manuscript was written through the contributions of all authors. All authors have given approval to the final version of the manuscript.

## ABBREVIATIONS

5’-dA•: 5’-deoxyadenosyl 5’-radical
5’-dAH: 5’-deoxyadenosine
FeS: iron-sulfur
SAM: *S*-adenosylmethionine
RS: Radical SAM
MPE: Mg-protoporphyrin IX monomethylester
PChlide: protochlorophyllide
Cbl: cobalamin
AdoCbl: adenosylcobalamin
MeCbl: methylcobalamin
OHCbl: hydroxocobalamin
MBP: maltose binding protein
MV: methylviologen
BME: β-mercaptoethanol
DTT: Dithiothreitol
DT: dithionite
LC-HRMS: Liquid chromatography-High resolution mass spectrometry
EPR: electron paramagnetic resonance
IMAC: immobilized metal affinity chromatography

