## Supporting Information for "Isolation and In vitro Characterization of BchE, the Cobalamin-Dependent Anaerobic Magnesium Protoporphyrin IX Monomethylester Cyclase"

**Supplemental contents**

1. **Materials and Methods**
2. **Supplemental Tables and Figures**
3. **Protein DNA and amino acid sequences**
4. **DFT Optimized Structure Coordinates**

#### Materials and Methods

##### Chemicals

Chemicals were used as received. *N*-(2-Hydroxyethyl)-piperazine-*N*'-(2-ethanesulfonic acid) (HEPES), potassium chloride, glycerol, yeast extract, magnesium sulfate, methanol, acetone, potassium phosphate monobasic, sodium phosphate dibasic, and dextrose were received from Fisher Scientific. Triton X-100, β-mercaptoethanol (BME), tryptone, sodium chloride, calcium chloride, tween-20, hydroxocobalamin acetate (OHCbl), dimethyl sulfoxide (DMSO), L-cysteine hydrochloride, phenylmethylsulfonyl fluoride (PMSF), ammonium acetate, *N*-[Tris(hydroxymethyl)methyl]-3-aminopropanesulfonic acid (TAPS), sodium hydrosulfite (dithionite, DT), L-tryptophan, N,N-dimethylformamide (DMF), dichloromethane (DCM), and sulfuric acid were ordered from Sigma Aldrich. L-Rhamnose and *N*-dodecyl-β-D-maltoside (DDM) were purchased from Chem-Impex. Lysozyme, DNase, isopropyl β-D-1-thiogalactopyranoside (IPTG), L-arabinose, and tris(2-carboxyethyl)phosphine (TCEP) were purchased from Gold Biotechnology. Iron(III) chloride, imidazole, and formic acid were from Thermo Scientific. LC/MS grade water, methanol, and acetonitrile were purchased from Honeywell. 1-Ethyl-3-(3-dimethylaminopropyl)carbodiimide (EDC) was from Oakwood Chemical. Peptone was from Beantown Chemical. 4-Dimethylaminopyridine (DMAP) was purchased from AstaTech Inc. Deuterated methanol was obtained from Acros Organics.

##### Synthesis of MPE

MPE was prepared from commercially available protoporphyrin IX (Sigma). Protoporphyrin IX (1.0 equiv.) was dissolved in a 1:1 (v/v) mixture of methanol and DMF (0.2 M), and EDC (1.0 equiv.) and DMAP (0.2 equiv.) were added to the stirred solution. After stirring at room temperature overnight, the reaction mixture was washed with water and brine. The organic layer was dried over anhydrous $Na_2SO_4$, filtered, and concentrated *in vacuo*. The crude material was purified by flash chromatography on silica gel to afford a 1:1 isomeric mixture of the monomethyl ester of protoporphyrin IX.

The isomeric mixture was then metalated by refluxing in pyridine at 120 °C with $Mg(ClO_4)_2$ (1.0 equiv.) for 1 hour to afford magnesium protoporphyrin monomethylester (MPE). The reaction mixture was concentrated under reduced pressure, and the crude product was purified by flash chromatography on silica gel to give MPE. After solvent removal, the product was redissolved in DMSO for storage and subsequent use. The concentration of the MPE solution was determined spectroscopically using an extinction coefficient of 18.5 $mM^{-1}$ $cm^{-1}$ at 552 nm.[1] Deuterated $CD_3$-MPE was prepared using the same procedure, substituting deuterated methanol for methanol in the esterification step.

##### Isolation of PChlide from *C. sphaeroides V3/ΔbciA*

*C. sphaeroides* V3/ *ΔbciA* is a strain that features both an unmapped mutant in a DPOR subunit-encoding gene and knockout of the *bciA* gene, encoding the native 8-vinyl (P)Chlide reductase.[2] Therefore, this strain accumulates 8V-PChlide, the direct product of the BchE reaction, when grown with Tween-20 in the media. The strain was grown in 4 L of PYS+ media (3 g/L peptone, 3 g/L yeast extract, 2 g/L NaCl, 2 mM $MgSO_4$, and 2 mM $CaCl_2$) from a single colony grown on a PYS+ agar plate. The culture grew for a total of three days in the dark at 30 °C and 150 RPM shaking. After 48 hours, 0.2% Tween-20 was added to the culture. After the final 24 hours of growth, the culture turned a vibrant green color. Cells were pelleted at 10,000 ×g for 10 min. The supernatant was loaded onto 4 parallel Waters Sep-Pak C18 cartridges with 2 g of stationary phase. After loading, a large green band was visible on the cartridges. The columns were washed with 20 mL of 10% acetone in water. Elution was performed using a stepwise gradient of increasing acetone by 10% every 10 mL. Fractions were collected by color and absorbance at 632 nm. Ultimately, 50-70% acetone fractions were found to elute PChlide. The pooled fractions were dried down under nitrogen to remove acetone, then lyophilized to remove water. The remaining solid was resuspended in DMSO. The concentration was determined by diluting an aliquot in 4:1 acetone:water and measuring the absorbance at 625 nm with an extinction coefficient of 28.0 $mM^{-1}$ $cm^{-1}$.[3,4] HPLC at 440 nm showed a single peak, which was identified as 8-vinyl PChlide by LC-MS. No 8-ethyl PChlide was detected.

<u>*Rp*BchE Expression and Purification</u>

Two plasmid constructs encoding *Rp*BchE were used, each with a distinct expression protocol. The first employed a pET-28a(+) vector induced with isopropyl β-D-1-thiogalactopyranoside (IPTG), and the second used a pRham vector induced with L-rhamnose. The amino acid sequence for *Rubrivivax pictus* (also known as *Pseudaquabacterium pictum*) BchE was obtained from Uniprot ID: A0A480ASV0. To improve solubility, a maltose-binding protein (MBP) fusion tag was appended to the C-terminus, linked via a Tobacco Etch Virus (TEV) protease cleavage site, followed by a C-terminal 10×His tag for IMAC purification. The entire sequence was codon-optimized using the IDT codon-optimization tool. The sequence was synthesized from Twist Bioscience and cloned into a pET-28a(+) plasmid at the *Nco*I and *Xho*I restriction sites. A glycine residue was introduced between Met1 and Arg2 to compensate for a frameshift introduced by the *Nco*I restriction site. The full insert sequence is given below. The resulting plasmid was used to transform *E. coli* BL21(DE3) cells, along with the pDB1282 (Fe-S cluster biosynthesis) and pBAD42-BtuCEDFB (cobalamin uptake) plasmids.[5,6] A 300 mL starter culture of LB media with 50 μg/mL kanamycin, 100 μg/mL ampicillin, and 50 μg/mL spectinomycin was inoculated from a single colony and grown overnight with shaking at 37 °C and 225 RPM. The following day, 10 mL was used to inoculate each 6 L flask containing 4 L of LB media with the same antibiotic concentrations. These cultures were grown with shaking at 180 RPM and 37 °C. At an $OD_{600}$ of 0.3, arabinose was added at a final concentration of 0.2% to initiate induction of genes on the pDB1282 and pBAD-BtuCEDFB plasmids. At this time, the media was supplemented with 1.3 μM hydroxocobalamin (OHCbl), 50 μM $FeCl_3$, 150 μM cysteine, and 0.25% glycerol. At an $OD_{600}$ of 0.8, the flasks were put on ice for 1 h before adding 50 uM IPTG. The cultures were shaken at 120 RPM overnight at 18 °C. The next day, the cells were harvested by centrifugation at 6,000 *g*, flash-frozen in liquid $N_2$, and stored at -80 °C.

It has previously been shown that deletion of the *btuR* gene affects the T7 expression system in *E. coli,* resulting in no expression from pET vectors.[6] Consistent with this observation, we did not detect induction of the *Rp*BchE–MBP pET-28a(+) construct in the Δ*btuR E. coli* BL21(DE3) strain. Therefore, we switched to a C-His pRham kanamycin-resistant expression vector (Lucigen), which uses L-rhamnose as the inducer. Using the pET-28a(+)-*Rp*BchE-MBP plasmid as a template, forward (5'-GAA GGA GAT ATA CAT ATG CGT GTA CTT CTG ATA CAT CCA-3') and reverse (5'-GTG ATG GTG GTG ATG ATG TTA ATG GTG ATG GTG ATG GTG-3') primers were designed based on the manufacturer's guidelines. Notably, the forward primer removes the Gly2 residue previously introduced in the pET-28a(+) vector. The reverse primer includes a stop codon to eliminate the vector-encoded 6×His tag, thereby allowing use of the existing C-terminal 10×His tag fused to MBP. PCR amplification was performed using Phusion DNA polymerase (New England Biolabs) following the manufacturer's protocol. The PCR products were purified by gel electrophoresis, extracted, and used to transform *E. coli* 10G competent cells along with the preprocessed C-His pRham vector. The final sequence was verified at the Penn State Genomics core facility.

The pRham-*Rp*BchE-MBP plasmid was used to transform competent *ΔbtuR E. coli* BL21(DE3) cells along with the pDB1282 and pBAD42-BtuCEDFB plasmids. For expression, a starter culture of 45 mL of LB media containing 100 μg/mL kanamycin, 200 μg/mL ampicillin, and 100 μg/mL spectinomycin was grown to an $OD_{600}$ of ~0.5 before being stored at 4 °C overnight. The next day, 10 mL was used to inoculate four 6 L flasks, each with 4 L of autoinduction media. The media consisted of 4 g/L tryptone, 2 g/L yeast extract, 1 mM $MgSO_4$, 25 mM $(NH_4)_2SO_4$, 50 mM $KH_2PO_4$, 50 mM $Na_2HPO_4$, 0.4% glycerol, 0.2% L-rhamnose, and 0.05% glucose. At an $OD_{600}$ of 0.3, 0.05% arabinose was added to induce expression of the genes on the pDB1282 and pBAD42-BtuCEDFB plasmids. At an $OD_{600}$ of 0.8, the flasks were placed on ice for 1 h before incubating them overnight (~18 hrs) at 18 °C with shaking at 120 RPM. The following day, cells were harvested by centrifugation at 6,000 ×*g* for 10 min at 4 °C. The resulting cell paste (~40 g) was frozen in liquid $N_2$ and stored at -80 °C.

All following protein preparation and handling were performed in a Coy anaerobic glovebox. Both proteins encoded on the pET-28a(+)-*Rp*BchE and pRham-*Rp*BchE plasmids were purified in the same manner. Cells (~40 g) were resuspended in 300 mL of lysis buffer (50 mM HEPES, pH 7.5, 300 mM KCl, 10% glycerol, 5 mM imidazole, 10 mM β-mercaptoethanol (BME), and 1% Triton X-100) with gentle stirring on ice for 30 min. While stirring, the suspension was supplemented with 1 mg/mL lysozyme, 0.1 mg/mL DNase, 0.2 mg/mL phenylmethylsulfonyl fluoride (PMSF), two tablets of Pierce protease inhibitor cocktail (EDTA-free), and 50 μM OHCbl. The cells were lysed by sonication at 70% amplitude for 15 sec on, 45 sec off for a total of 5 min on. Insoluble material was removed by centrifugation at 40,000 ×*g* for 1 h. The supernatant was run over a Ni-NTA IMAC column pre-equilibrated with lysis buffer. The retained protein was washed with 200 mL of 50 mM HEPES, pH 7.5, 300 KCl, 10% glycerol, 10 mM BME, and 50 mM imidazole. The same buffer contents were used for elution, but with imidazole increased to 300 mM. Protein was collected based on dark-brown color and concentrated to ~5 mL with an Amicon 50 kDa MWCO ultrafiltration centrifugation filter (EMD Millipore). The protein was buffer-exchanged on a G-25 resin column pre-equilibrated in protease buffer (50 mM HEPES, pH 7.5, 300 mM KCl, 15% glycerol, and 10 mM BME). His-tagged TEV protease was added at 2 mg/mL and incubated at room temperature for 1 h, followed by overnight incubation on ice. Chemical reconstitution of Cbl and the Fe-S clusters was performed simultaneously as described previously.[7] The following day, the protein was centrifuged at 10,000 ×*g* for 10 min to remove insoluble particulates. The protein was run over a Ni-NTA column pre-equilibrated with protease buffer. The flow-through was collected and concentrated to >5 mL. For size-exclusion chromatography, the protein was loaded onto a HiPrep 16/60 S-300 HR column (Cytiva) using an AKTA fast protein liquid chromatography (FPLC) system housed in an anaerobic chamber. The column was equilibrated with 50 mM HEPES, pH 7.5, 300 mM KCl, 10% glycerol, and 1 mM dithiothreitol (DTT). Fractions were collected based on absorbance at 280 nm and brown color. A majority of the protein elutes in the void volume (40 mL), consistent with aggregates, with a smaller peak at 65 mL. We found the aggregate fraction to be active, but all the work presented was performed on fractions collected from the later-eluting peak. The fractions were concentrated to 5 mL and protein concentration was determined by the Bradford assay using a bovine serum albumin (fraction V) standard. Iron and cobalamin content were determined as previously described.[7, 8] UV-vis spectra were recorded with an Agilent Cary 50 UV-vis spectrometer.

*Rubrivax gelatinosus* (Uniprot Q7X2C7) and *Rhodobacter capsulatus* (Uniprot P26168) BchE were also expressed and purified during this work. Both constructs were cloned into a pET-26b(+) plasmid at the *Nde*I and *Xho*I restriction sites to utilize the C-terminal 6×His-tag. A TEV protease site was included in the sequence immediately before the *Xho*I site so the affinity tag could be removed during purification. With this modification, the expression and purification procedures remained the same as with protein produced from the pET-28a(+)-*Rp*BchE-MBP plasmid. These enzymes were not found to be active with any tested biological reductants. Ti citrate was the only reductant found to afford any activity, but only the *575* species off-pathway product (see main text) was observed. Due to the lack of activity, further studies of these proteins were not pursued. Additional BchE sequences that were not found to be soluble upon expression and purification include the following: *Chlorobaculum tepidum* (Uniprot H2VFK1), *Chloracidobacterium thermophilum* (Uniprot G2LJY1), *Rhodoplanes roseus* (Uniprot A0A327KNI8), and *Rubrivivax albus* (Uniprot A0A437JY33).

<u>*Rc*FdxA, *Rg*FdxA, and *Rg*FNR Expression and Purification</u>

The amino acid sequences for each ferredoxin and ferredoxin reductase were obtained from Uniprot. These include *Rhodobacter capsulatus* FdxA (Uniprot ID: D5AP15), *Rubrivivax gelatinosus* FdxA (Uniprot ID: I0HRP6), and *Rubrivivax gelatinosus* ferredoxin reductase (*Rg*FNR) (Uniprot ID: I0HR46). The DNA sequences were optimized for expression in *E. coli B* using the IDT Codon Optimization Tool. The resulting sequences were ordered from Gene Universal Inc. and cloned into pET-26b(+) plasmids at the *Nde*I and *Xho*I restriction sites such that the encoded proteins would contain a C-terminal 6×His affinity tag. The plasmids were used to transform *E. coli* BL21(DE3) cells, along with pDB1282. For expression, an overnight culture of 300 mL of LB media with 50 μg/mL kanamycin and 100 μg/mL ampicillin was grown from a single colony. The next day, 20 mL was used to inoculate 8 L of LB media (2 × 4 L) with the same antibiotic concentrations, and the culture was grown at 37 °C and 180 RPM shaking. At an $OD_{600}$ of 0.3, $FeCl_3$, cysteine, and arabinose were added at concentrations of 50 μM, 300 μM, and 0.2 %, respectively. At an $OD_{600}$ of 0.6, IPTG was added to 50 μM, and the temperature was lowered to 18 °C with overnight shaking at 120 RPM (~18 h). The cells were harvested by centrifugation at 6,000 ×*g* and 4 °C for 10 min. The cell paste was frozen in liquid nitrogen and stored in -80 °C. Expression of *Rg*FNR followed the same procedure but excluded the pDB1282 plasmid and the addition of $FeCl_3$, cysteine, arabinose, and ampicillin.

Purification was performed in a Coy anaerobic chamber. All buffers consisted of 50 mM HEPES, pH 7.5, 200 mM KCl, 5% glycerol, 10 mM BME but varied in imidazole content. Lysis, wash, elution, and storage buffer contained 5 , 50 , 300 , and 0 mM imidazole, respectively. Frozen cell paste (~25 g) was resuspended in 200 mL of lysis buffer and stirred on ice for 30 min. During this time, 1 mg/mL lysozyme, 0.1 mg/mL DNAse, and 0.2 mg/mL PMSF were added to the suspension. The cells were lysed by sonication at 70% amplitude, 15 sec on, 45 sec off for a total of 5 min of sonication time. The mixture was then centrifuged at 40,000 ×*g* for 1 h at 4 °C to remove insoluble particulates. The supernatant was loaded onto an IMAC column and washed with 100 mL of wash buffer containing 50 mM imidazole. The protein was eluted with 50 mL of elution buffer and concentrated to ~4 mL using an Amicon ultracentrifugal filter with a 3 kDa cutoff. The remaining protein was buffer-exchanged into storage buffer using two preequilibrated PD-10 columns in parallel. Notably, the storage buffer did not contain BME, as it was found to react with a product of the BchE reaction, as addressed in the main text. The protein was aliquoted before freezing and stored in liquid nitrogen. *Rg*FNR was purified in the same way as the ferredoxins, but with 1 mM flavin adenine dinucleotide (FAD) added to the lysis buffer. Protein concentration was determined by Bradford assay with a bovine serum albumin (fraction V) standard.

Electron Paramagnetic Resonance Spectroscopy

Electron paramagnetic resonance (EPR) measurements were performed on a Magnettech 5000 X-band ESR spectrometer with an ER 4102ST resonator. An ER4112-HV Oxford Instruments variable-temperature helium-flow cryostat was used to control the temperature to 10 K or 70 K for $[Fe_4S_4]^{1+}$ cluster and cob(II)alamin analysis, respectively. Measurements at 10 K were performed with a modulation amplitude of 1 mT and a microwave power of 0.315 mW (25 dB). 70 K measurements were performed with a modulation amplitude of 0.8 mT and a microwave power of 10 mW (10 dB). All spectra consisted of 5 scans and were collected with a 100 kHz modulation frequency.

Samples were prepared in 4 mm O.D. quartz tubes (Wilmad) and contained 300 μM BchE, with SAM and MPE added at 1 mM and 0.6 mM, respectively, when applicable. BchE was diluted into storage buffer for cobalamin analysis. For Fe-S cluster analysis, the protein was buffer-exchanged into 50 mM TAPS (pH 8.5), 350 mM KCl, and 10% glycerol, and reduced with 1 mM dithionite. Simulations of EPR spectra were performed with Spincount (ver. 7.3.8832.24157) developed by Professor Michael Hendrich at Carnegie Mellon University.

Activity assays

A typical reaction was performed in reaction buffer consisting of 50 mM HEPES, pH 7.5, 200 mM KCl, 5% glycerol, 1 mM $MgCl_2$, and 0.01% N-dodecyl-β-D-maltoside (DDM). BchE (10 μM by Cbl quantification) was incubated with 1 mM SAM, 50 μM MPE, and 20 μM tryptophan (internal standard) for 10 min before initiating the reaction with reductant (Ti citrate or dithionite each added at 1 mM). The MV-FMN-NADPH reduction system consisted of 50 μM methyl viologen (MV) and 500 μM flavin mononucleotide (FMN), and 2 mM NADPH. When a ferredoxin reduction system was used, *Rc* or *Rg*FdxA (50 μM) and *Rg*FNR (25 μM) were incubated with BchE in the reaction mixture for 10 min before initiating with NADPH. All incubations were performed at room temperature (23 °C), and after initiation, reaction tubes were placed in a water bath equilibrated to 35 °C. When quantification of MeCbl or AdoCbl was desired, reactions were performed in the dark or under red light. Sample prep for LC-MS analysis was also performed under low light. When applicable, photolytic cleavage of the Co-C bond of BchE-bound AdoCbl was performed with a uvBeast V3 385-395nm UV Flashlight. The flashlight was placed ~5 cm from the enzyme solution in a glass tube. Light exposure totaled 45 min on ice with gentle mixing every 5 min. Aliquots were taken at specified timepoints and quenched 1:1 in either 150 mM $H_2SO_4$ in isopropyl alcohol for LC-HRMS or 0.2% ammonia in methanol for HPLC analysis using a photodiode array spectrometer (HPLC-PDA). In both cases, samples were centrifuged at 10,000 ×*g* for 10 min at 4 °C to remove insoluble particulates.

Acid-quenched samples were injected into a Thermo Scientific Vanquish UHPLC system coupled to a Thermo Scientific Q Exactive HF-X mass spectrometer with an H-ESI ion source.

Chromatographic separation was performed with a Restek Ultra C8 column (50 mm × 2.1 mm) with a 3 μm particle size. A gradient method was used with solvents A (0.1% aqueous formic acid) and B (0.1% formic acid in acetonitrile). The chromatographic gradient began at 1% solvent B from 0–1.5 min. Solvent B was then increased linearly from 1% to 80% between 1.5 and 6 min, followed by an increase to 100% from 6–9 min. The column was held at 100% B from 9–10.5 min, after which B was decreased to 1% from 10.5–11 min. The column was equilibrated at 1% B from 11–13 min, resulting in a total method time of 13 min. The column oven was maintained at 30 °C, and the flow rate was 0.3 mL/min. A 5 μL injection volume was used, and chromatograms were monitored at 440 nm.

Mass spectra were collected in positive full-scan mode from 100–1000 *m/z* with a resolution of 120,000 and an AGC target of $3 \times 10^6$ for the entire 13-min run. Data acquisition and analysis were performed using Thermo Scientific Xcalibur 4.2.47. When absolute quantification was not feasible, relative analyte abundance was calculated as the raw peak area normalized to the peak area of the internal standard, L-tryptophan.

Ammoniated methanol–quenched samples were analyzed on an Agilent 1290 Infinity II UHPLC system coupled to a 1290 Infinity II Diode Array Detector (DAD). Chromatographic separation was performed using the same C8 HPLC column described previously, with solvent A consisting of 40 mM ammonium acetate in water and solvent B consisting of methanol. The gradient program was as follows: 70% B from 0–1 min, increased to 100% B from 1–7 min, held at 100% B from 7–9 min, decreased to 70% B from 9–10 min, and equilibrated at 70% B from 10–12 min, giving a total run time of 12 min. UV detection was performed at 440 nm, and full UV–vis spectra were extracted from peaks of interest. This method maintained analytes at neutral to basic pH, allowing the porphyrins to be detected with their central magnesium ions bound. Although initially developed for activity assays coupled to mass spectrometry, MS analysis proved challenging because multiple *m/z* species with comparable intensities were observed for each porphyrin. For example, MPE formed a methanol adduct in positive mode and an acetic acid adduct in negative mode. Under acidic conditions, the magnesium ion was released, but the resulting [M+1] signals were more intense and free of adducts.

Ferredoxin screening

Multiple reduction systems were tested for compatibility with isolated BchE. These included spinach ferredoxin and ferredoxin reductase (purchased from Sigma), *Homo sapien* FDX1 and FDXR, and *E. coli* flavodoxin (flv) and flavodoxin reductase (flr), which were previously used in our lab.[5, 9] Three ferredoxins (Uniprot ID: I0HRP6, I0HVB1, and I0HPF2) and a ferredoxin reductase (Uniprot ID: I0HR46) were selected from *R. gelatinosus.* These were expressed and purified as stated above. BchE activity was determined using the assay conditions outlined above. Timepoints were quenched with 0.2% ammonia in methanol (20 μL reaction mixture into 200 μL methanol) and centrifuged to remove insoluble particulates. The supernatant was analyzed by fluorescence spectroscopy on a Varian Cary Eclipse Fluorescence Spectrophotometer. Spectra were collected at emissions of 550-700 nm with excitation at 440 nm. PChlide shows a peak at 633 nm and the 575 species at 620 nm under these conditions. Only the I0HRP6 ferredoxin and I0HR46 ferredoxin reductase together showed accumulation at these wavelengths, which were later validated by LC-MS.

Ferredoxins from *R. capsulatus* (Uniprot ID: D5ARY6, D5AP15, D5ARX7, and D5AM35) were also tested. These were chosen based on annotation as [$Fe_4S_4$] cluster-containing ferredoxins.

Each gene was cloned into pET-26b(+) vectors using the *Nde*I and *Xho*I restriction sites and expressed in *E. coli* BL21 (DE3) cells along with the pDB1282 plasmid. 40 mL cultures were pelleted and resuspended anaerobically in 1 mL assay buffer. The cells were lysed by vortexing with 0.5 g of 0.1 mm glass beads for 3 min. The lysed cells were centrifuged, and the supernatant was used to add BchE, NADPH, *Rg*FDR, MPE, and $MgCl_2$ at the typical assay concentrations. Aliquots were quenched at 1 h and 2 h timepoints and analyzed by HPLC-PDA. Only D5AP15 supported the production of peaks corresponding to PChlide and the *575* species.

Isolation of *575* species

The 575 species (described in the main text) was produced in appropriate amounts for isolation by scaling up a typical BchE reaction to 10 mL and using Ti citrate as the reductant. After 30 min of reaction time in a 35 °C water bath, the reaction was quenched 1:1 with 0.2% ammonia in methanol. The quenched reaction was centrifuged at 10,000 ×*g* for 30 min before purification by preparative HPLC. Chromatographic separation of the 575 species from MPE, PChlide, hydroxy-MPE, and keto-MPE was performed on an Agilent 1260 Infinity II Preparative HPLC with an Agilent 5 Prep-C18 column (50 × 21.2 mm) with a 5 µm particle size. Solvents A (aqueous 40 mM ammonium acetate) and B (methanol) were used in a gradient method.

The method used a constant flow rate of 20 mL/min with a 0.5 mL injection volume. From 0–2 min, the mobile phase consisted of 75% solvent B. Solvent B was then increased from 75% to 100% over 2–12 min, held at 100% from 12–14 min, and decreased back to 75% from 14–15 min. An additional 2 min of equilibration at 75% B was included before the next run. HPLC traces were monitored at 440 nm.

Fractions containing the 575 nm-absorbing species, identifiable by their green color, were pooled. Methanol was removed under a stream of nitrogen, and the remaining aqueous phase was lyophilized. To remove ammonium acetate, the dried material was resuspended in 3 mL of 50:50 water/methanol and loaded onto a C18 Sep-Pak cartridge (Waters). The cartridge was washed with 10 mL of 50:50 water/methanol, and the compound was eluted with acetone. The eluate was dried and subsequently resuspended in anaerobic DMSO inside a glovebox. The absence of MPE, PChlide, hydroxy-MPE, and keto-MPE was confirmed by LC-MS.

Computational methods

All calculations were performed using ORCA version 5.0.4. Initial structural models were generated in Chemcraft (v1.8). A molecule of acetic acid was included to coordinate the central magnesium ion, producing a five-coordinate complex consistent with the adduct observed by mass spectrometry under the HPLC conditions used to obtain the experimental UV–vis spectra.

Geometry optimizations were carried out using the B3LYP hybrid functional with the def2-TZVP basis set, the RIJCOSX approximation, Grimme's D3 dispersion correction, and a methanol CPCM solvation model.[10-15] Time-dependent DFT was performed on these optimized structures using the double-hybrid B2PYLP functional with the def2-TZVP basis set.[16] Additional settings included the RIJCOSX approximation, TightSCF convergence criteria, Grimme's D3 dispersion correction, and a methanol CPCM solvent model. Ten roots were calculated for the TD-DFT portion of the calculation using the Tamm-Dancoff approximation.[17] The theoretical absorbance spectra were generated with the orca_mapspc tool using a a full width at half maximum (FWHM) of 750 $cm^{-1}$.

To achieve optimal agreement with the experimental spectra, transition energies were uniformly shifted by −0.19 eV.

*R. capsulatus* growth and native BchE activity assays

This procedure was adapted from that of Gough et al.[18] *Rhodobacter capsulatus* strain SB1003 was grown from a single colony overnight in 5 mL PYS+ media at 30 ºC with shaking at 225 RPM. The following day, 50 µL was used to inoculate 5 mL of PYS+ media for aerobic growth and 10 mL of media for anaerobic growth. Anaerobic growth was achieved by using a stopper and vinyl tape to seal a sterile glass tube, leaving it filled near the top. Stoppered growths were continuously exposed to light with a Philips 60W white LED bulb. Five cultures were grown under both aerobic and anaerobic conditions at 30 ºC with shaking at 150 RPM for 72 h. Aerobic growths appeared red in color, whereas anaerobic growths were brown. The $OD_{700}$ of each culture was measured before pelleting the cells by centrifugation. The cells were resuspended in 50 mM potassium phosphate buffer (pH 7.3) to an effective $OD_{700}$ of 1.0 for each culture. The cells were then freeze-thawed twice with liq $N_2$.

The native BchE reaction was performed by incubating 200 µL of the resuspended cells with 10 µM MPE, 10 mM of nicotinamide (NAM), and 30 mM $MgCl_2$ for 1 h at 30 ºC with shaking at 150 RPM. Dependence on MPE, NAM, and Mg was determined by excluding each individually in separate reactions. The reaction mixture was centrifuged at 10,000 ×g for 5 min, and the supernatant was removed. The cells were resuspended in 0.3 mL acetone/0.35 M ammonia (4:1 v/v ratio) and 0.4 mL hexanes was used to extract the porphyrin pigments. Fluorescence was performed on the acetone layer at 440 nm excitation, with a 550-700 nm emission scan. To perform LC-MS, 100 µL of the ammoniated acetone layer was dried off and resuspended in 50/50 water/isopropyl alcohol containing 150 mM $H_2SO_4$. Prior to injection, samples were spiked with 20 µM L-tryptophan to use as an internal standard. HRMS was performed using the method outlined for assays.

**Supplemental Table & Figure List**

**Table S1.** EPR simulation parameters for Cobalamin and Fe-S cluster of *Rp*BchE

**Figure S1.** SDS-PAGE gels of typical *Rp*BchE expression and purification procedures

**Figure S2**. EPR spectra of reduced BchE incubated with aza-SAM, SAH, and SAM with MPE

**Figure S3**. Formation of methionine, 5'-dAH, and SAH with different chemical reductants

**Figure S4**. Optimization of initiation for BchE reaction

**Figure S5**. Formation of SAH, methionine, 5'-dAH for MeCbl dependence assay

**Figure S6**. Formation of methionine and 5'-dAH for AdoCbl dependence assay

**Figure S7**. EPR spectroscopy for *Rc*FdxA and *Rg*FdxA

**Figure S8**. Formation of 5'dAH, SAH, and MeCbl for ferredoxin activity assays

**Figure S9**. Formation of adenine and hypoxanthine for ferredoxin activity assays

**Figure S10**. MS/MS fragmentation of Hydroxy-MPE

**Figure S11**. MS/MS fragmentation of Keto-MPE

**Figure S12**. Overlain plots of PChlide, Keto-MPE, and Hydroxy-MPE formation with both *Rg* and *Rc*FdxA reducing systems

**Figure S13**. Formation of Hydroxy-MPE and Keto-MPE using chemical reductants for the BchE reaction.

**Figure S14**. Formation of Hydroxy-MPE, Keto-MPE, and PChlide with and without pre-methylation of BchE Cobalamin.

**Figure S15**. Formation of PChlide in anaerobic versus aerobic conditions

**Figure S16**. Mass spectrum of MPE, hydroxy-MPE, and keto-MPE after reactions in natural abundance versus $^{18}O$-water enriched buffer

**Figure S17**. Formation of *575* species with chemical reductants

**Figure S18**. MS/MS fragmentation of *575* species with natural abundance and $D_3$-methyl ester

**Figure S19**. Mass spectrum of MPE, PChlide, and *575* species under neutral mobile phase conditions

**Figure S20.** Calculated TD-DFT absorbance spectra of MPE, PChlide, *575*-ring, and *575*-olefin structures highlighting the porphyrin Q-bands.

**Figure S21**. Formation of *575* + BME Michael addition product with 5 mM BME

**Figure S22**. Formation of PChlide and 575 influenced by incubation with TCEP

**Figure S23**. Analysis of *R. capsulatus in vivo* BchE activity assays

**Supplemental Tables**

**Table S1.** EPR simulation parameters for Cobalamin and Fe-S cluster of *Rp*BchE**.** *g*-values and Hyperfine couplings are shown with *g*-strain (σ*g*) and hyperfine strain (σ*A*) shown in parentheses.

| | $g_1$ ($\sigma g_1$) | $g_2$ ($\sigma g_2$) | $g_3$ ($\sigma g_3$) | $A_1$ ($\sigma A_1$)* | $A_1$ ($\sigma A_1$)* | $A_1$ ($\sigma A_1$)* | LW # |
|---|---|---|---|---|---|---|---|
| Fe-S cluster | 2.041 (0.022) | 1.926 (0.020) | 1.875 (0.039) | | | | 1.0 |
| Cob(II)alamin | 2.391 (0.016) | 2.303 (0.018) | 2.003 (0.007) | 215 (23) | 190 (16) | 395 (5) | 0.8 |

* Hyperfine Coupling (MHz)

### Linewidth (mT)

**Supplemental Figures.**

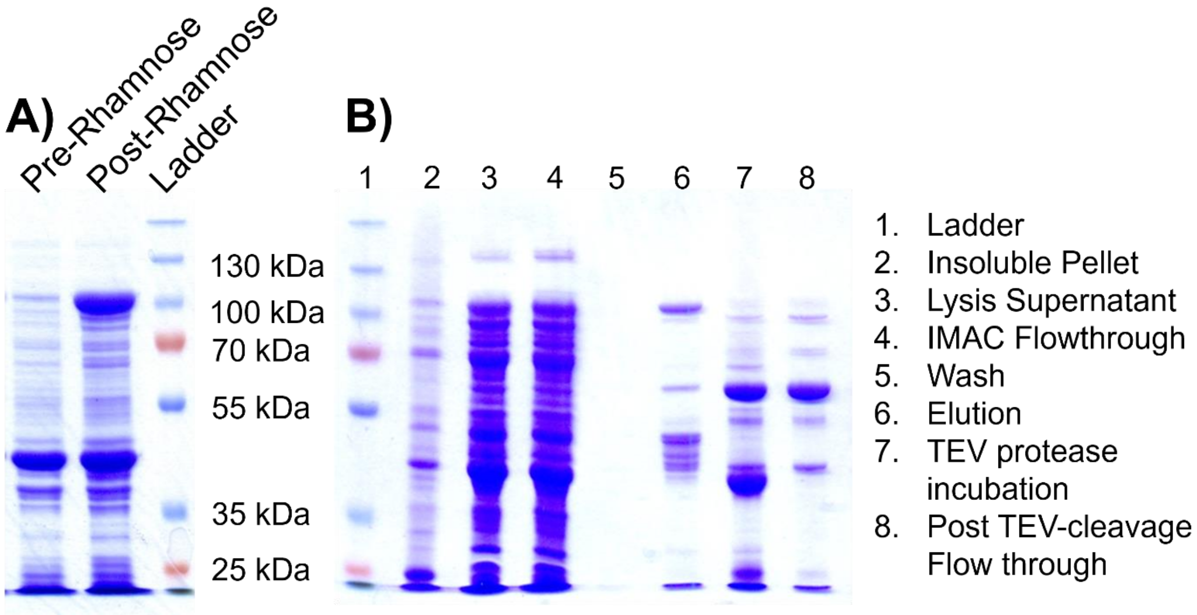


**Figure S1.** SDS-PAGE gels of expression and purification of *Rp*BchE. The heterologous expression of *Rp*BchE in the strain of *ΔbtuR E. coli* BL21 (DE3) cells is shown in **A**) before and after addition of L-rhamnose with overnight incubation at 18 ºC. A prominent band at ~100 kDa (theoretical 106 kDa) is seen in the post-rhamnose lane, consistent with the expected mass of *Rp*BchE-MBP. B) shows the gel from a typical purification of *Rp*BchE. Notably, lane 6 shows *Rp*BchE-MBP immediately after elution from an IMAC column. Lane 7 shows that the addition of TEV-protease results in bands consistent with cleaved *Rp*BchE (theoretical 64 kDa) and MBP with a 10× His tag (theoretical 42 kDa). Upon a second flow through the IMAC column, the MBP fusion protein is removed (lane 8). The resulting MBP-free *Rp*BchE was further purified by gel filtration, for which the resulting SDS-PAGE is shown in the main text.

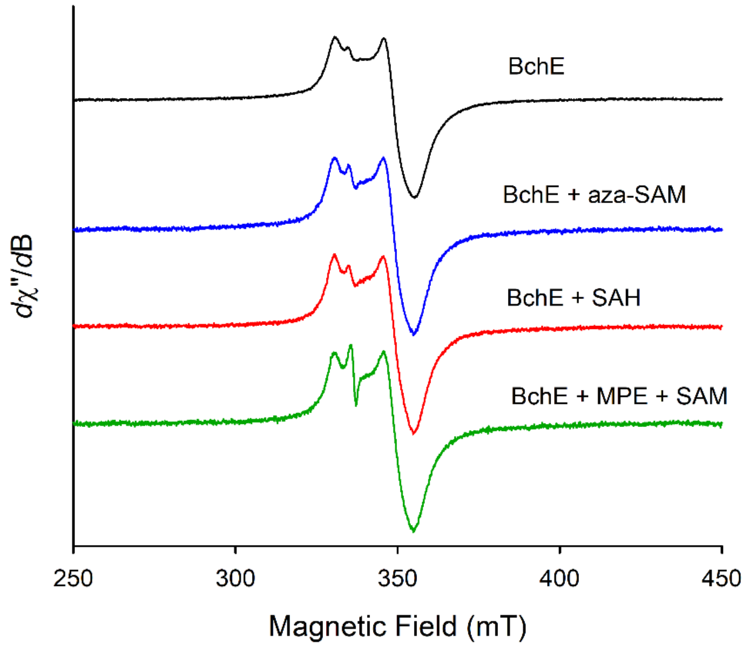


**Figure S2.** EPR spectra of *Rp*BchE reduced with 1 mM dithionite (black) and incubated with 1 mM aza-SAM (blue), SAH (red), or 0.6 mM MPE and 1 mM SAM (green).

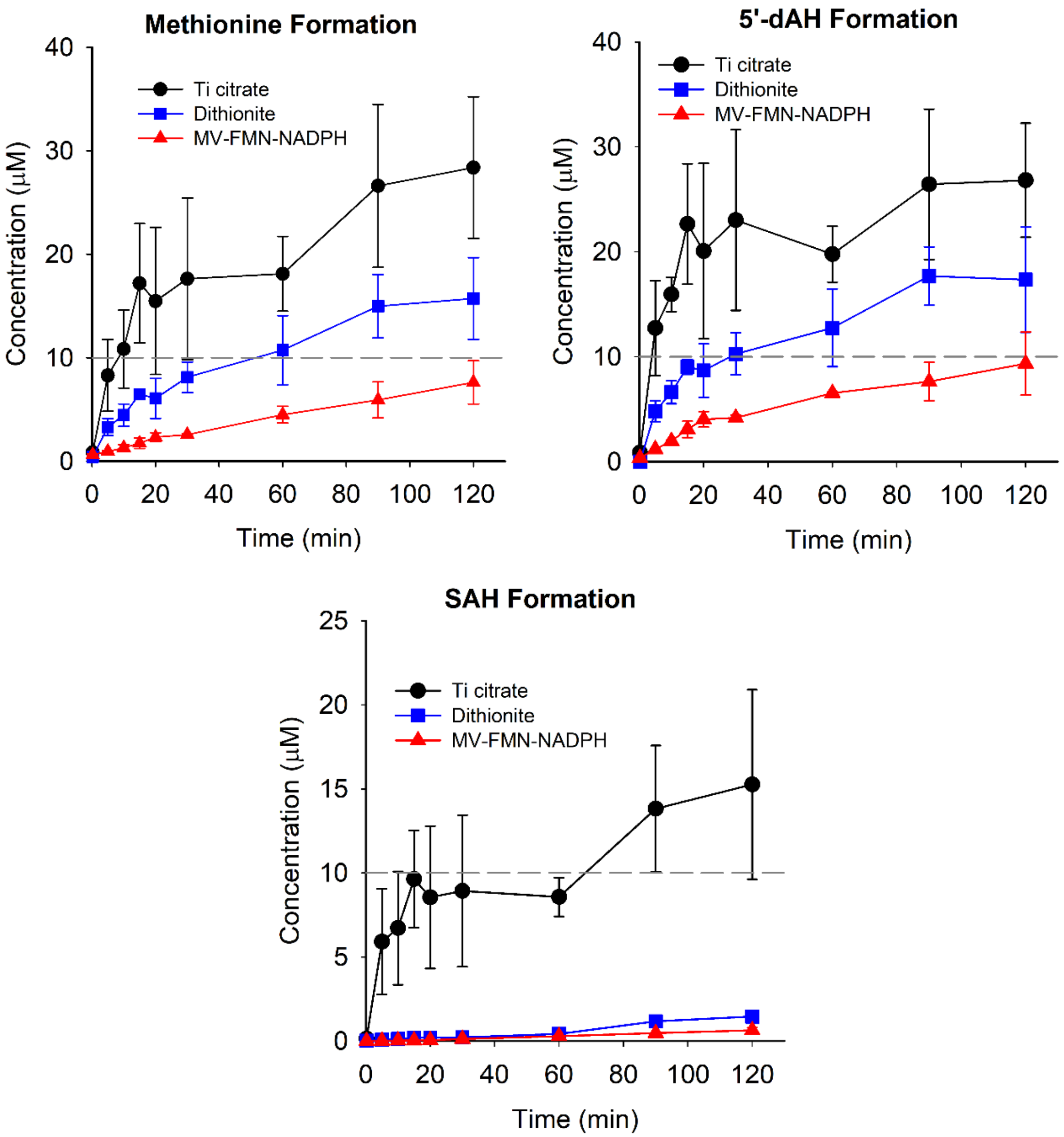


**Figure S3**. Methionine, 5’-dAH, and SAH formation from enzymatic reactions of BchE using either Ti citrate (*black*), dithionite (*blue*), or methyl viologen/FMN/NADPH (*red*) reducing systems. For reference, protein concentration is indicated by a gray dashed line at 10 µM. Reactions were run in triplicate with error bars representing the standard deviation for each timepoint.

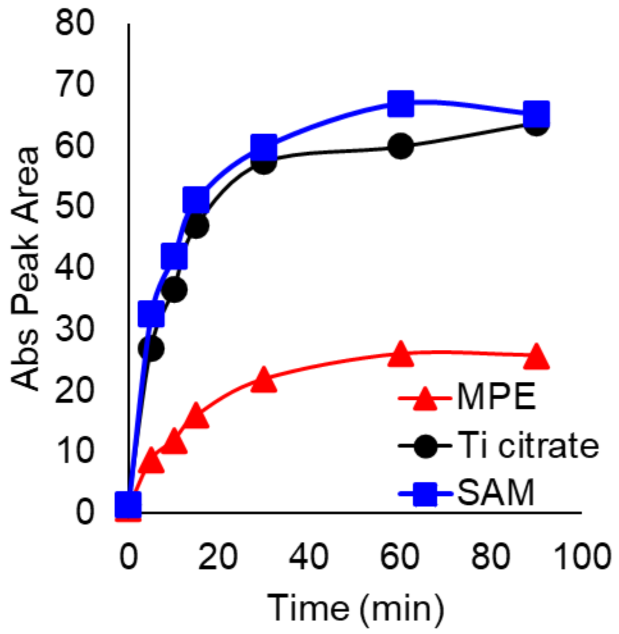


**Figure S4**. The BchE reaction was optimized by running three different reactions in parallel where either MPE, SAM, or Ti citrate (reductant), would be used to initiate the reaction. The figure displays the HPLC peak area associated with PChlide formation at various time points. Initiating with SAM and Ti citrate yielded similar results, whereas initiating with MPE resulted in much less PChlide formation. Each reaction was run in singlet.

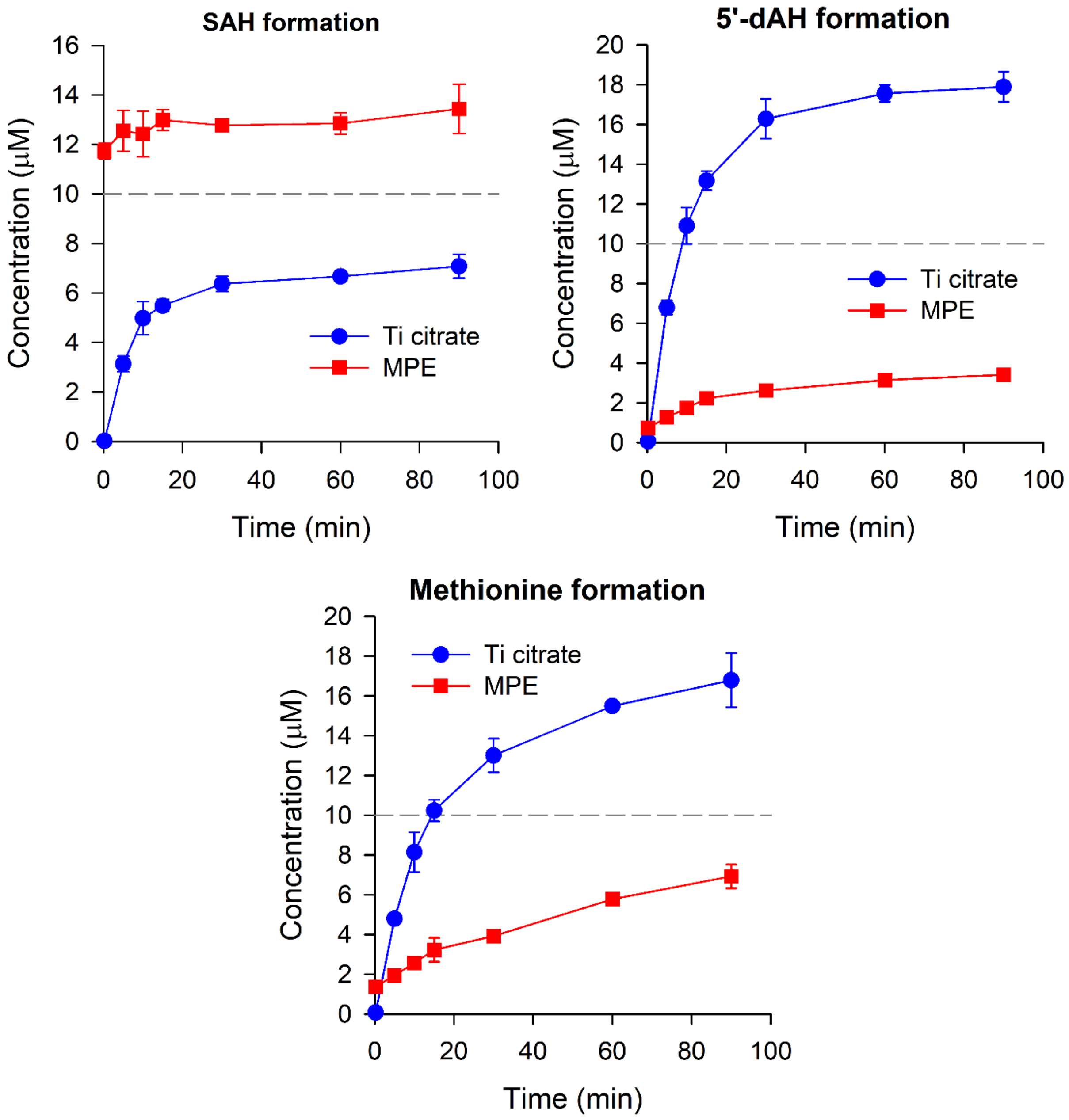


**Figure S5**. SAH, 5'-dAH, and methionine formation from enzymatic reactions of BchE Initiated with Ti citrate (*blue*) or MPE (*red*). For reference, protein concentration is indicated by a gray dashed line at 10 µM. Reactions were run in triplicate with error bars representing the standard deviation for each timepoint.

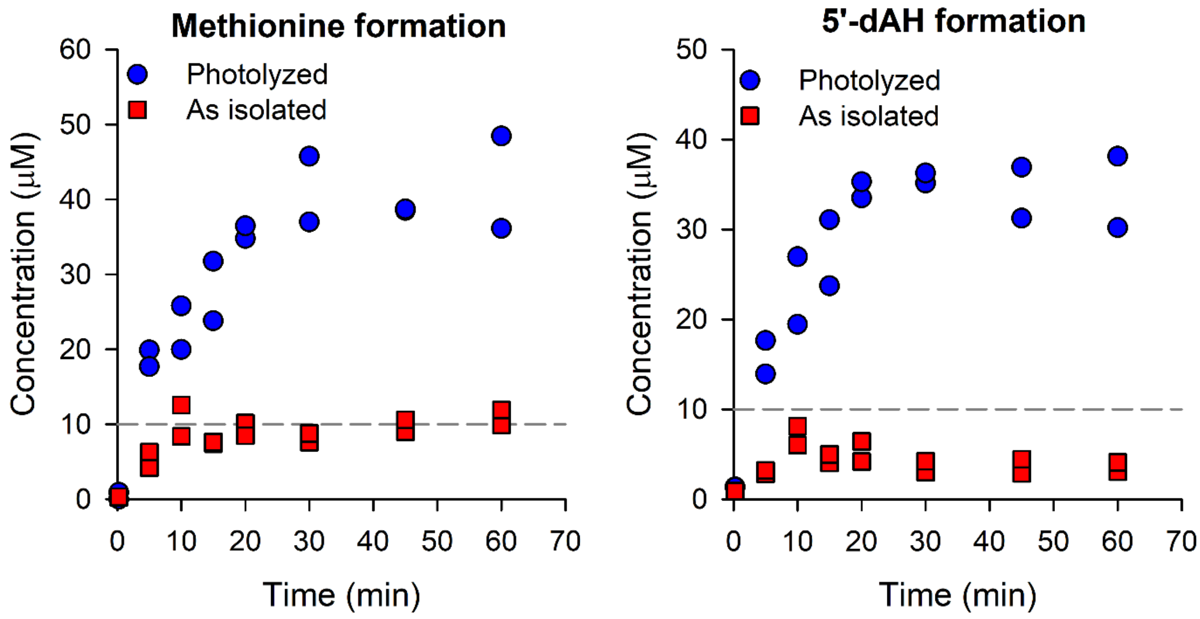


**Figure S6**. Methionine and 5'dAH formation from enzymatic reactions of BchE purified with contaminating AdoCbl. Each reaction was conducted either with as-isolated protein (*red squares*) or after photolysis to remove the Ado ligand (*blue circles*). For reference, protein concentration is indicated by a gray dashed line at 10 µM. Reactions were run in duplicate.

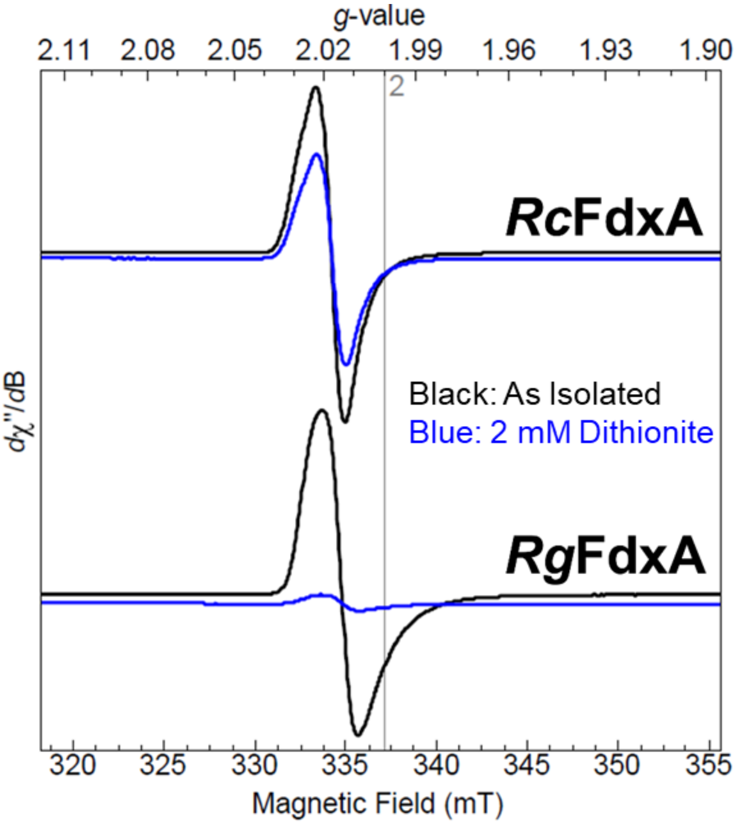


**Figure S7**. EPR spectra of *Rc*FdxA and *Rg*FdxA, as-isolated (*black*) or as-isolated and incubated with 2 mM dithionite (*blue*). Each spectrum consists of three scans at 10 K. Microwave frequency: 9.435 GHz; modulation amplitude: 10 G; microwave power: 0.1 mW.

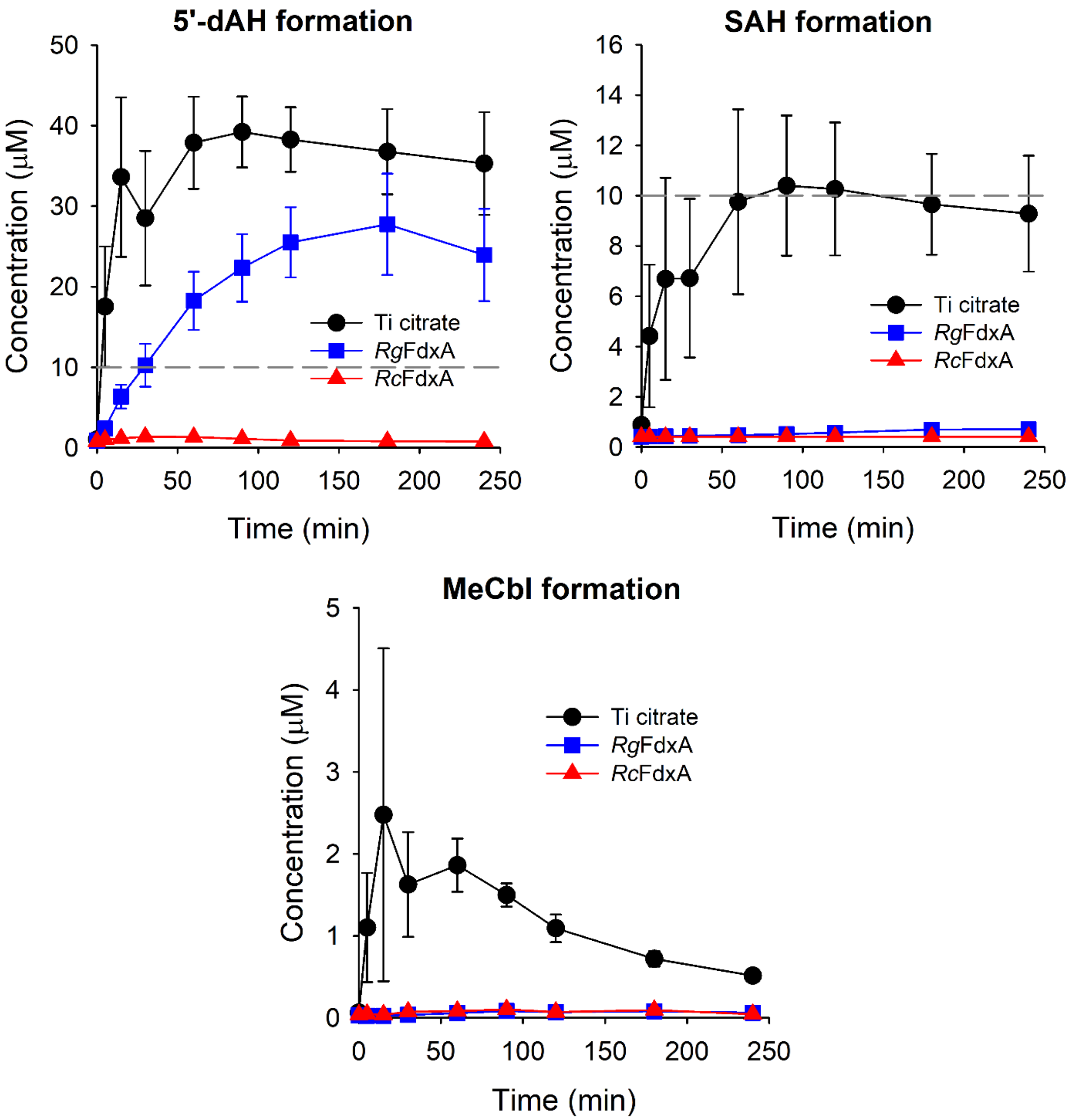


**Figure S8**. 5’-dAH, SAH, and MeCbl formation from BchE reactions using different reducing systems. Ti citrate is shown in *black circles*, *Rg*FdxA with *blue squares.* and *Rc*FdxA with *red triangles*. Both ferredoxin systems also utilize *Rg*FNR and NADPH. For reference, protein concentration is indicated by a gray dashed line at 10 µM. Reactions were run in triplicate with error bars representing the standard deviation for each timepoint.

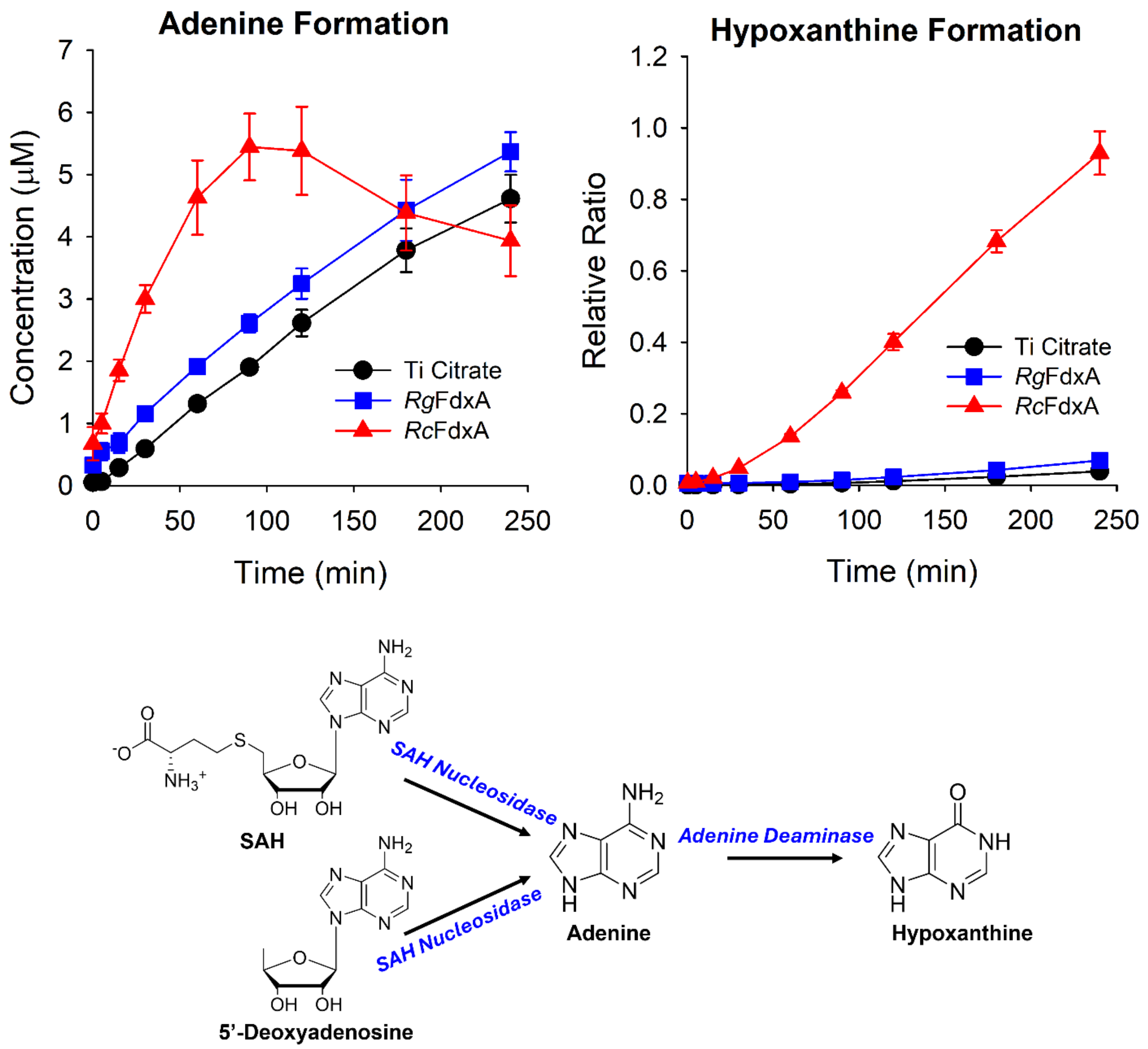


**Figure S9**. Adenine and hypoxanthine formation from BchE reactions detected by HRMS. Ti citrate is shown in *black circles*, *Rg*FdxA with *blue squares.* and *Rc*FdxA with *red triangles*. Both ferredoxin systems also utilize *Rg*FNR and NADPH. Reactions were performed in triplicate with error bars representing the standard deviation for each timepoint. A scheme is provided to depict how adenine and hypoxanthine can be formed from SAH and 5'-dAH in the presence of SAH nucleosidase and adenine deaminase.

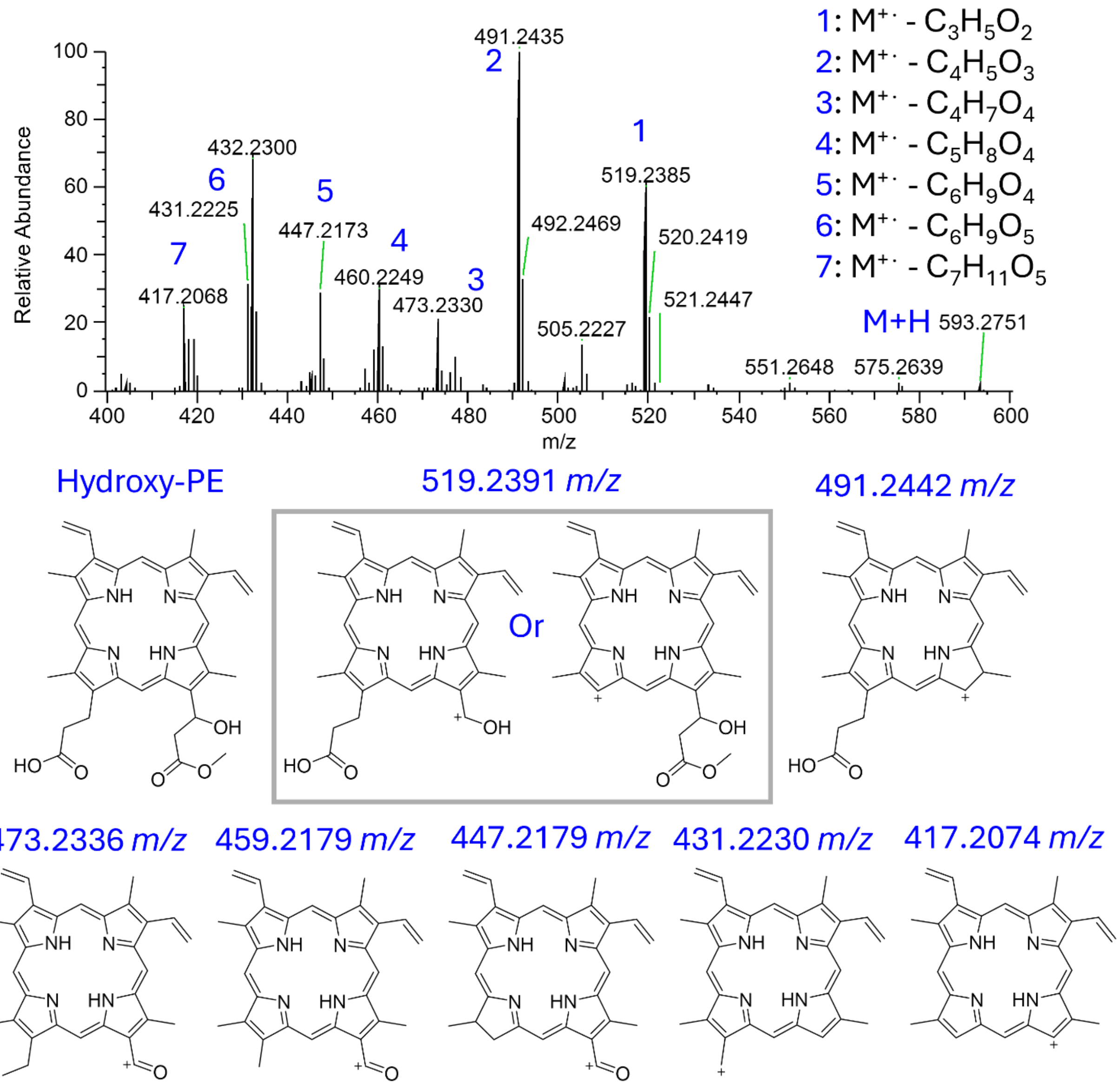


**Figure S10**. MS/MS fragmentation of hydroxy-protoporphyrin IX monomethylester (hydroxy-PE) by HRMS with a higher-energy collisional dissociation (HCD) of 50 eV in positive mode. The higher-abundance fragments are assigned proposed structures, which correspond to those previously reported for magnesium-bound porphyrins.[19]

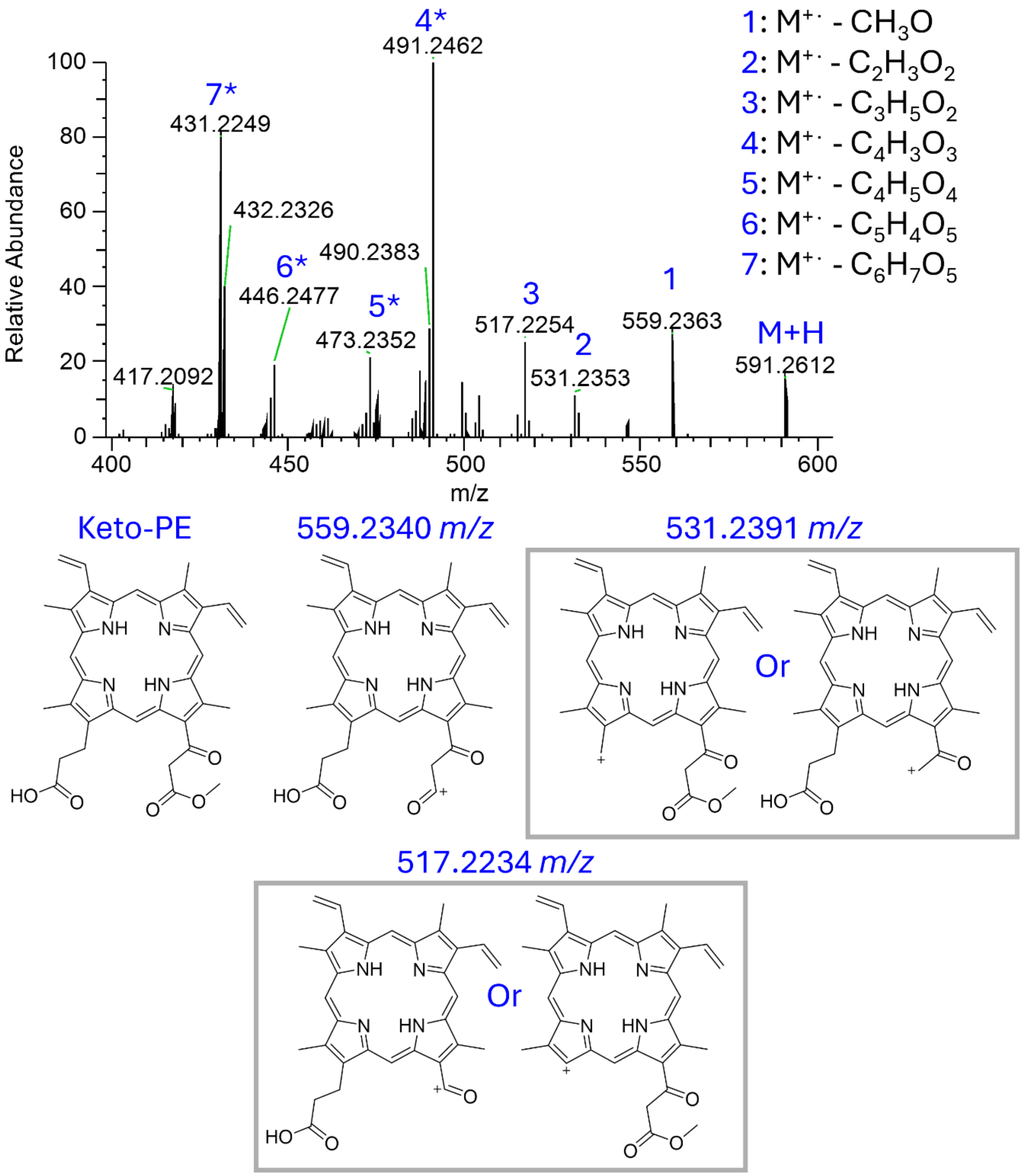


**Figure S11**. MS/MS fragmentation of keto-protoporphyrin IX monomethylester (Keto-PE) by HRMS with an HCD of 50 eV in positive mode. The higher abundance fragments are assigned proposed structures. Peaks marked with an "*" are assumed to be the same as proposed in **Figure S11** for peaks of the same *m/z*. These mirror those observed in previous work on magnesium-bound porphyrin.[19]

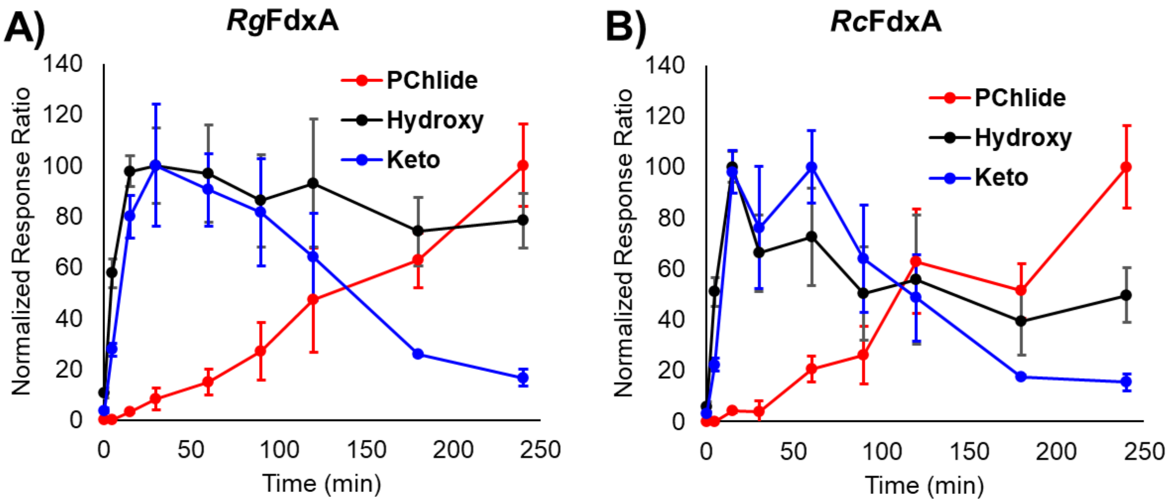


**Figure S12**. Overlaid plots of PChlide, Keto-MPE, and Hydroxy-MPE formation with both *Rg* and *Rc*FdxA reducing systems

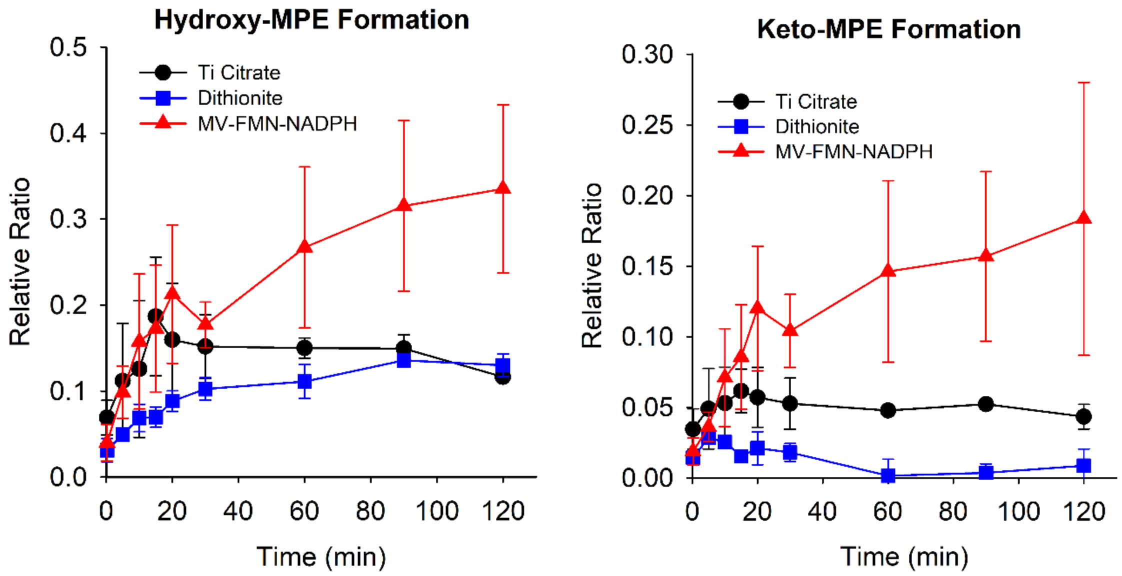


**Figure S13**. Time-dependent formation of Hydroxy-MPE and Keto-MPE for the BchE reactions using either Ti citrate, dithionite, or methyl viologen/flavin mononucleotide/NADPH reducing systems.

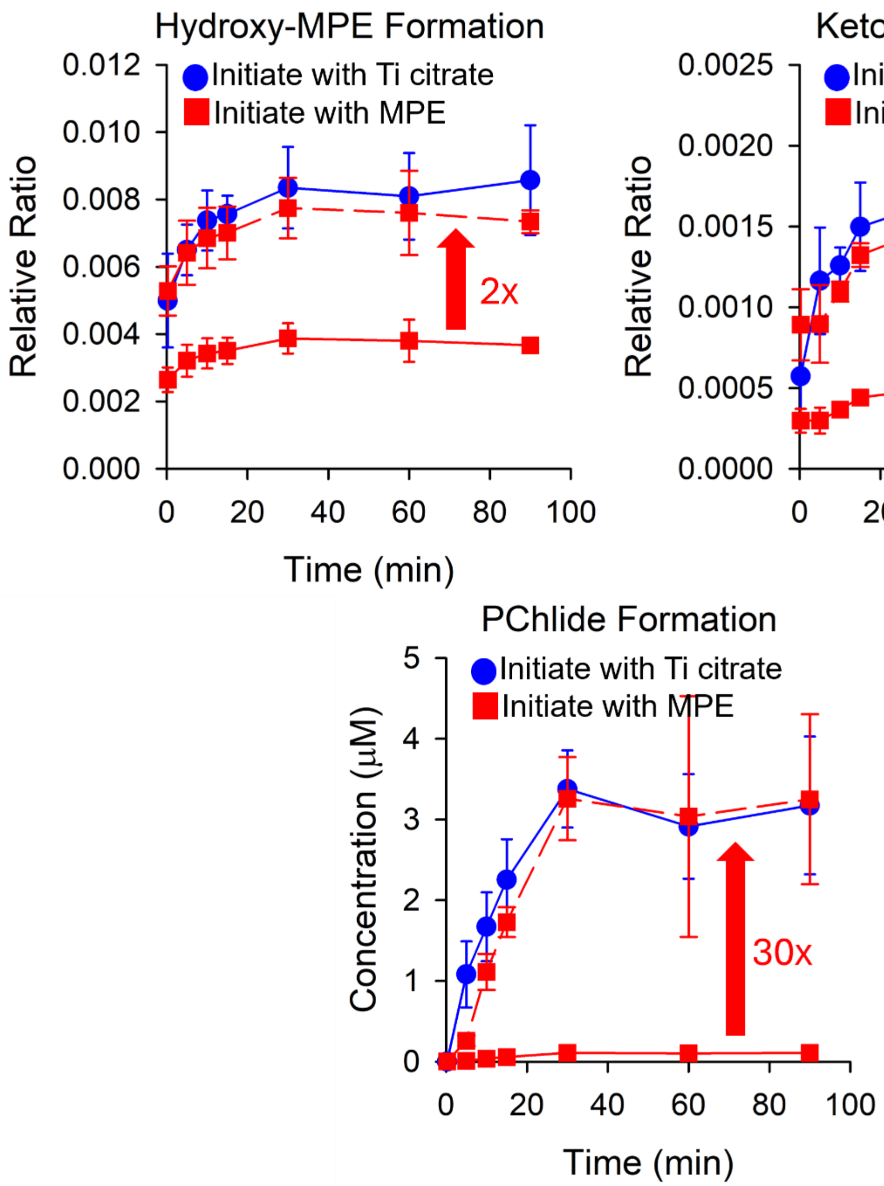


**Figure S14**. Time-dependent formation of Hydroxy-MPE, Keto-MPE, and PChlide for the BchE reactions using Ti citrate reductant. *Blue circles* indicate timepoints where the reaction was incubated with MPE and SAM and initiated with Ti citrate. *Red squares* correspond to parallel reactions incubated with Ti citrate and SAM and initiated with MPE. The *red dashed* traces correspond to the values after scaling by the factor indicated in each plot for the traces to overlay.

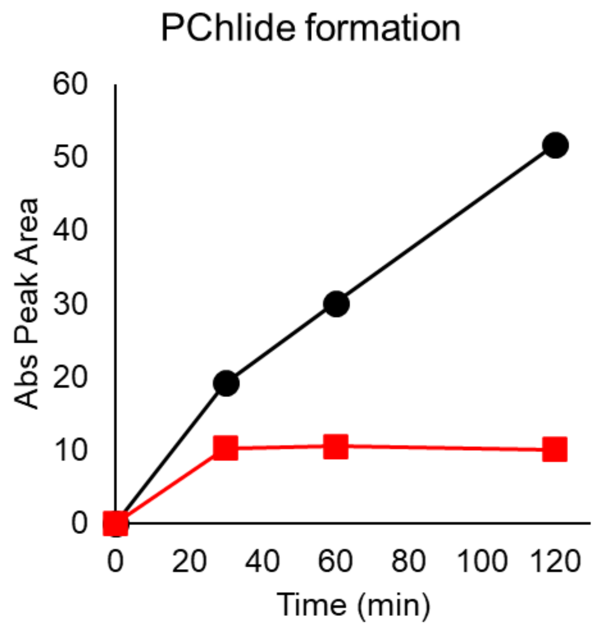


**Figure S15**. PChlide formation from a BchE reaction under anaerobic conditions (*black circles*) and when exposed to an oxygen-containing atmosphere (*red squares*). Each point represents the HPLC peak area associated with PChlide formation at various time points. Each reaction was run in singlet.

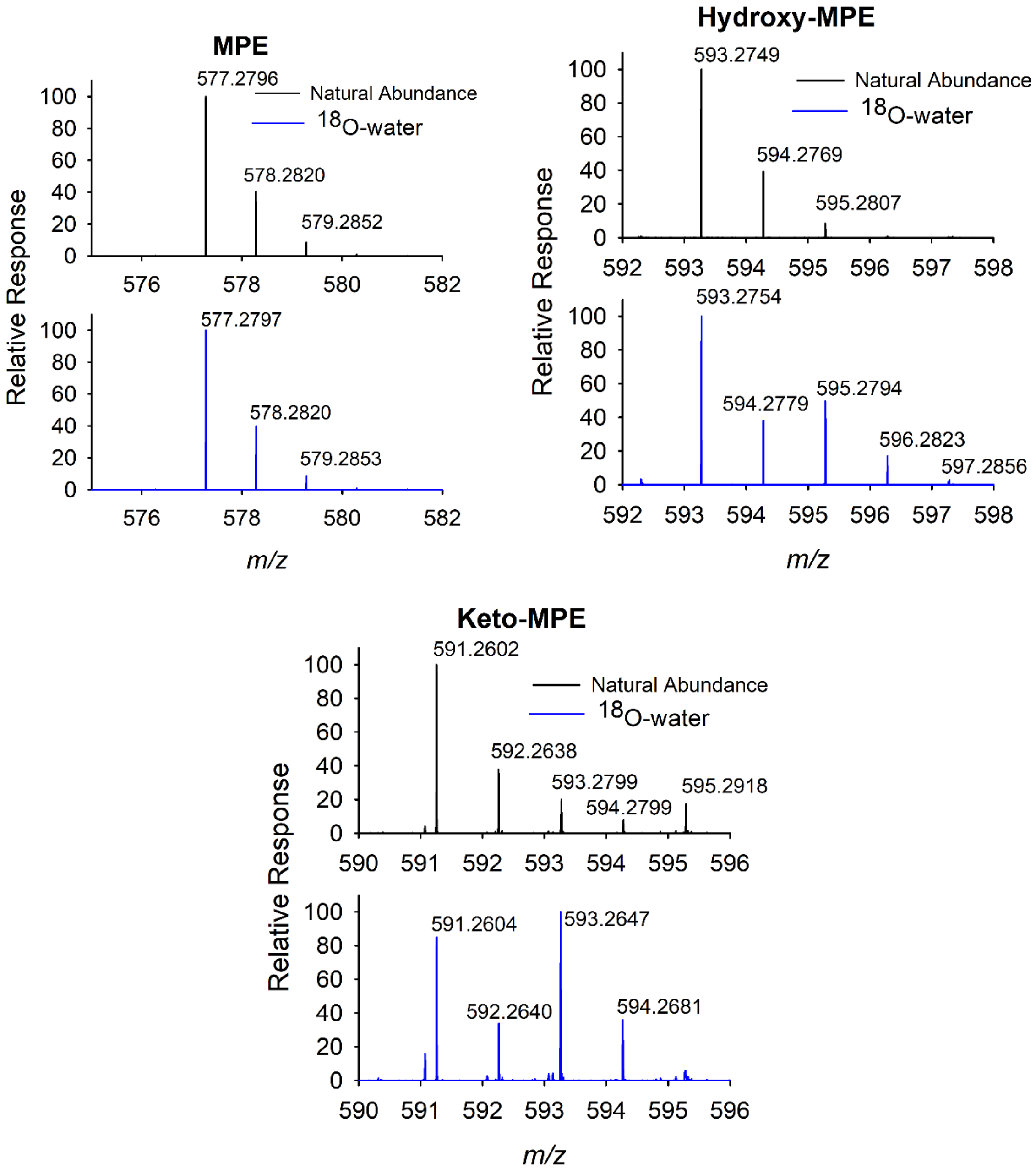


**Figure S16**. Mass spectra of MPE, Hydroxy-MPE, and Keto-MPE when a BchE reaction was conducted in either natural abundance aqueous buffer (black) or $^{18}O$-enriched water. Both reactions were performed anaerobically and quenched with methanol. The spectra of MPE remain unchanged between reactions prepared with natural abundance water and isotopically enriched ($H_2{}^{18}O$) water. Hydroxy-MPE shows an increase in *m/z* consistent with the +2 Da expected for $^{18}O$ incorporation, but does not yield ~70% of the signal expected, as seen for Keto-MPE and PChlide (main text, **Figure 5G**).

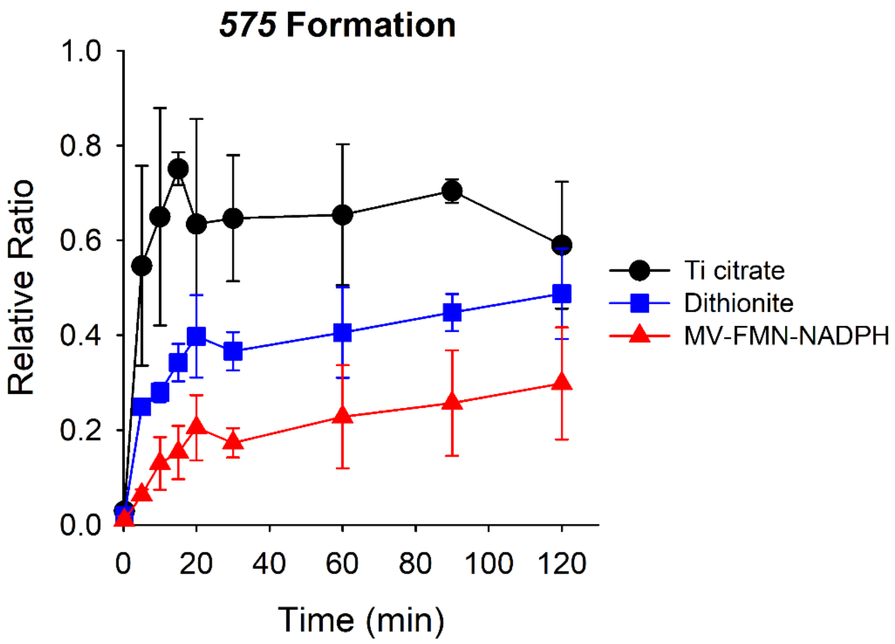


**Figure S17**. *575* formation from enzymatic reactions of BchE using either Ti citrate (*black*), dithionite (*blue*), or methyl viologen/FMN/NADPH (*red*) reducing systems. Values are reported as the relative ratio of the HRMS *575* peak area to the peak area of tryptophan (internal standard). Reactions were conducted in triplicate with error bars representing the standard deviation for each timepoint.

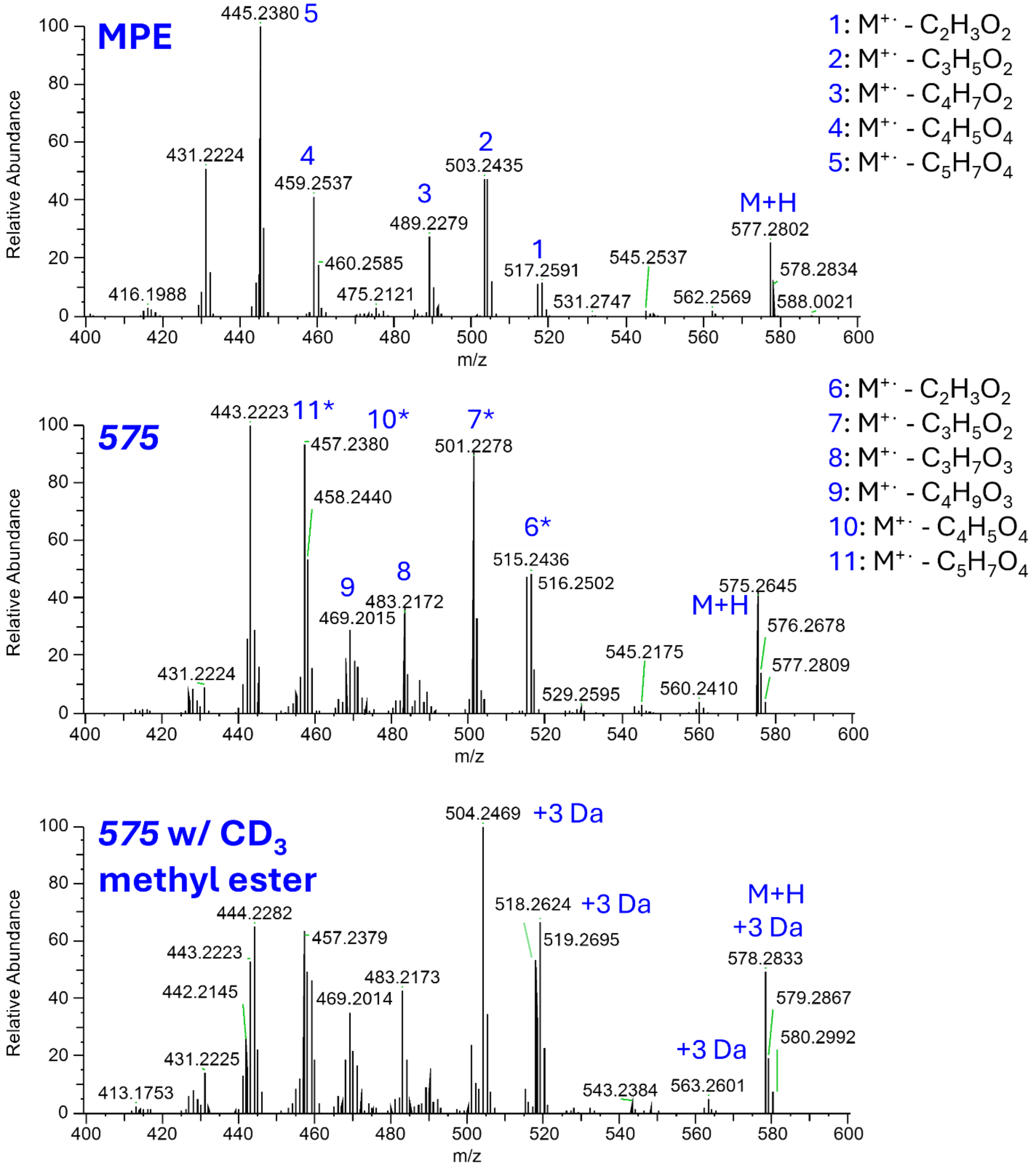


**Continued on next page.**

### Continued...

Unsaturation on ester or carboxylate chain?

#### MPE is 1:1 isomeric mixture $D_3$-MPE

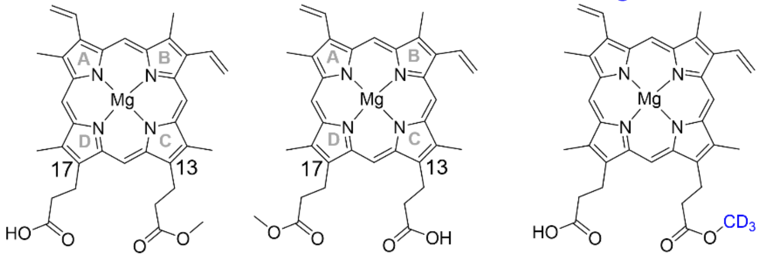


#### *Olefin*

518.2630 *m/z* 504.2473 *m/z*

**Ester**

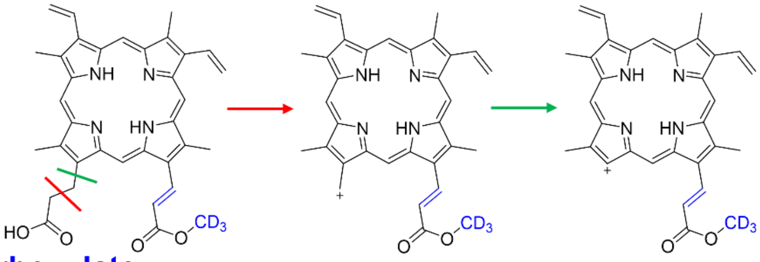

**Carboxylate**

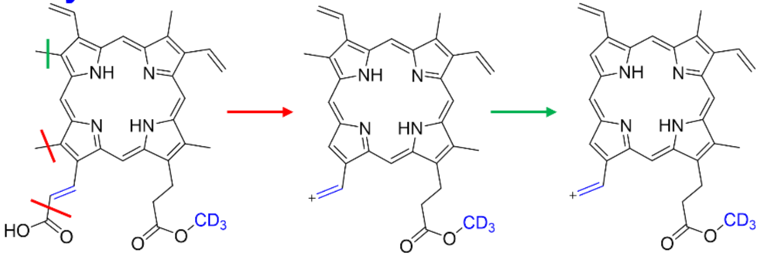

#### *Ring*

**Ester**

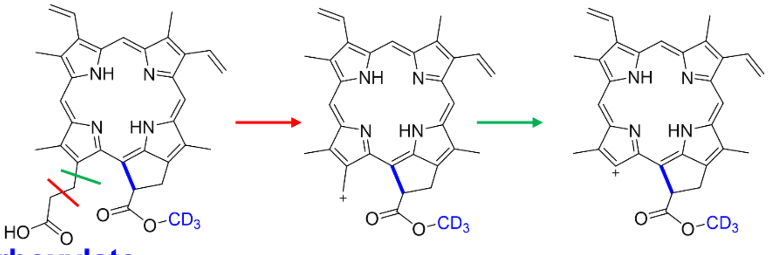

**Carboxylate**

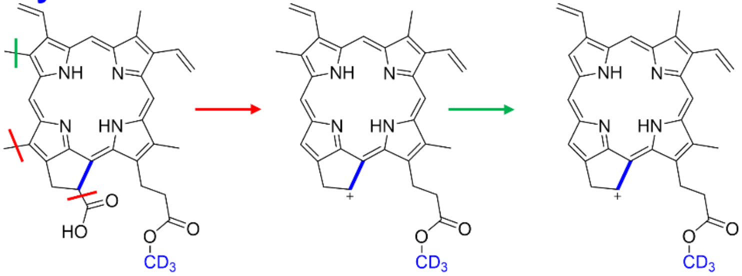

#### Other Fragments

483.2179 *m/z*

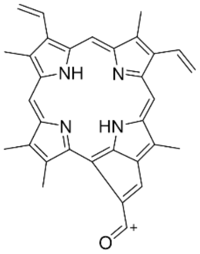

469.2023 *m/z*

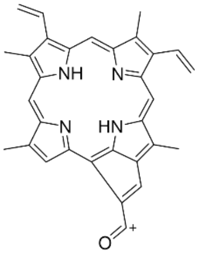

457.2387 *m/z*

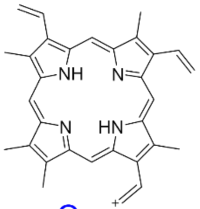

Or

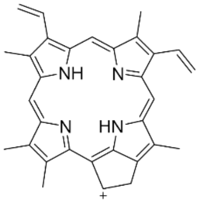

443.2230 *m/z*

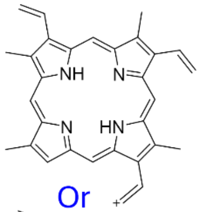

Or

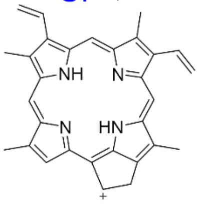

**Figure S18**. The MPE and *575* species were subjected to LC-MS/MS fragmentation on a high-resolution mass spectrometer. The intent of this experiment was to determine if the unsaturation was due to olefin or ring formation and whether it occurred on the alkyl chain with the ester or with the carboxylic acid. Each species was fragmented with an HCD of 50 eV. Parent ions were selected in the method via an inclusion list. The fragmentation spectra correlate with the retention times of each species, as observed with a parallel-scan method. The top spectrum shows fragmentation for the starting material, MPE. Directly below is the fragmentation spectrum for the *575* species. For clarity, *575* differs from the mass of MPE by the loss of two protons, or in other words, one additional degree of unsaturation. The spectra for MPE and *575* share common fragmentation events, i.e. the same neutral loss to produce said fragment. These peaks are noted with an "*" for the *575* spectrum. Notably, these peaks all feature the loss of two protons relative to the MPE signals, indicating they contain the unsaturated moiety. The *575* species was also prepared from a BchE reaction using MPE with a deuterated methyl ester group. The fragmentation spectrum of $CD_3$-*575* is shown in the bottom spectrum. Many fragments are of the same *m/z* as with the natural abundance spectrum, but some have a shift of +3 Da. This indicates the +3 Da fragments still contain the deuterated methyl ester group, and the remaining fragments do not. By referencing which fragments contain the additional unsaturation (by comparing MPE to *575*) and the methyl ester (by comparing *575* to $CD_3$-*575*), we can determine that the 504.2469 and 518.2624 *m/z* peaks observed in the deuterated spectrum must feature both the unsaturation moiety and methyl ester group. We predict that the unsaturation is due to either an olefin or ring formation. The native reaction of MPE to PChlide features ketone and ring formation on the chain with the methyl ester instead of the chain with the carboxylate. However, the *575* species may be an off-pathway product, as opposed to a true intermediate. In this instance, it is possible that *575* results from the incorrect positioning of MPE in the BchE active site, where hydrogen abstraction occurs on the carboxylate chain. With this in mind, we assigned possible structures to the peaks correlating to 518 and 504 *m/z* as seen in the $CD_3$-*575* spectrum. These structures were assigned with the postulate of containing both the deuterated methyl ester and unsaturation moiety. Notably, the MPE and $CD_3$-MPE used in this work are isomeric mixtures in which the methyl ester could be on the 13- or 17-carbon chain. These compounds are symmetric except for the relative locations of the carbon 3 and 8 vinyl groups. This prevents us from explicitly assigning whether the unsaturation occurs on the 13- or 17-carbon chain.

Cleavages were predicted for the olefin and ring structures on both the ester and carboxylate chains. We show fragments with the physiological form of MPE with the ester on the 13-carbon chain. Olefin and ring structures on the ester chain can both easily reproduce the 518 and 504 *m/z* peaks with a single cleavage indicated by the red and green lines, respectively. If the olefin/ring forms on the carboxylate chain, multiple cleavages would be needed. The 518 *m/z* peak could be produced by cleavage between carbons $17^2$ and $17^3$ in addition to a methyl group on the porphyrin ring (*red lines*). We illustrate the methyl group on carbon 18, but carbons 2, 7, and 12 are also possible. Furthermore, to produce the 504 *m/z* peak, yet another methyl group would need to be cleaved (*green*). The 504 and 518 peaks can be accounted for with straightforward single cleavages when the unsaturation occurs on the ester chain. However, two or three cleavages would be needed to achieve these masses if the unsaturation is on the carboxylate chain. We believe this suggests the ester chain is the site of the modification. Assuming this assignment is correct, we also assigned structures for the other peaks, which are likely to feature the unsaturation moiety. Based on these proposed structures, neither an olefin nor a ring structure can be ruled out. Notably, each of these structures did not require cleavage of one of the carbon 2, 7, 12, or 18 methyl or 3, 8 vinyl groups. Therefore, the predicted fragments are also applicable to the isomeric form of MPE with the ester on the 17-carbon chain.

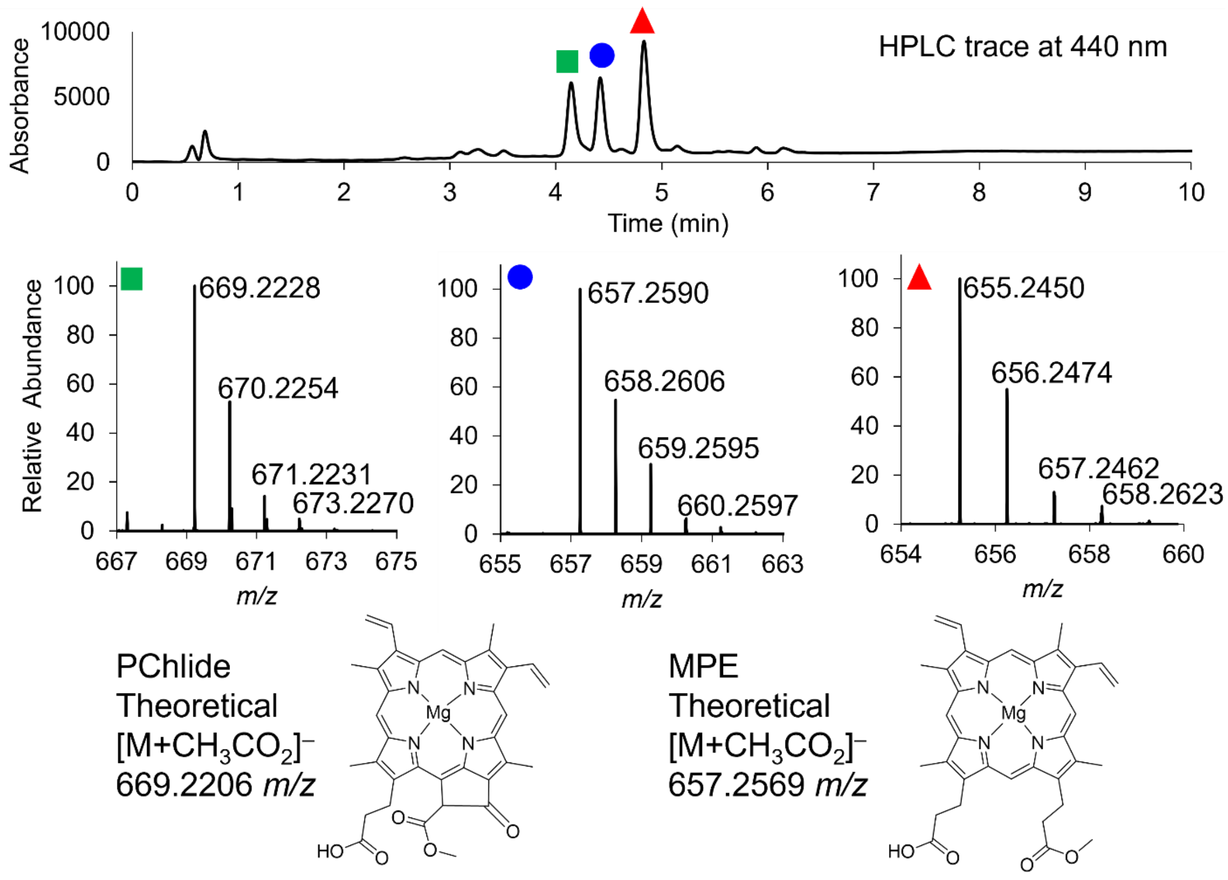


**Figure S19**. The HPLC method for analyzing the quenched BchE reaction at neutral pH was performed on the HRMS instrument to verify the identity of each peak. PChlide (*green square*) and MPE (*blue circle*) were both detected predominantly with acetate adducts in negative mode. The HPLC peak presumed to be the 575 species (*red triangle*) showed a spectrum consistent with MPE, with the loss of two protons. This reproduces the observed mass when run in acidic conditions in positive mode.

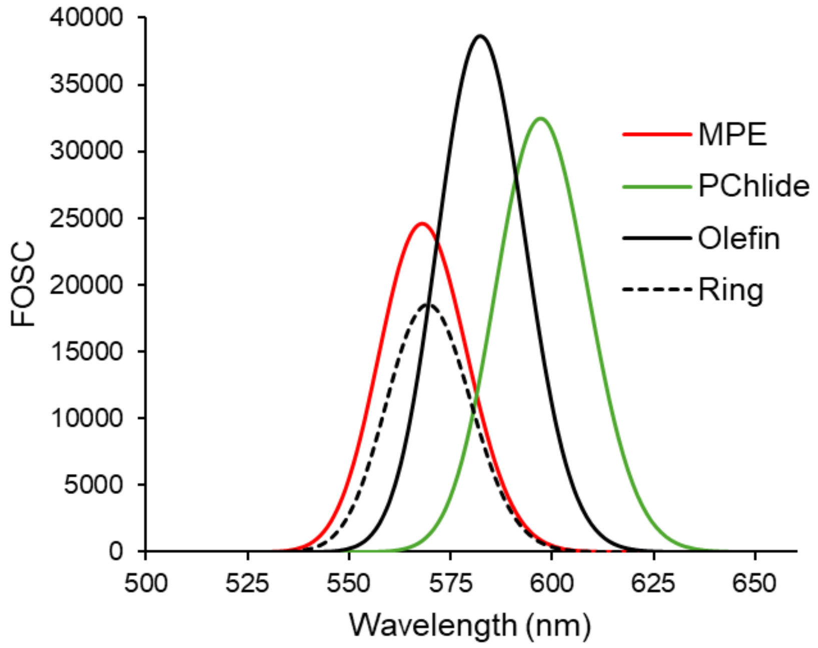


**Figure S20**. Calculated TD-DFT spectra for the Q-band region of optimized MPE, PChlide, *575*-ring, and *575*-olefin structures. As with the Soret bands (main text), a -0.19 eV shift was applied to each transition.

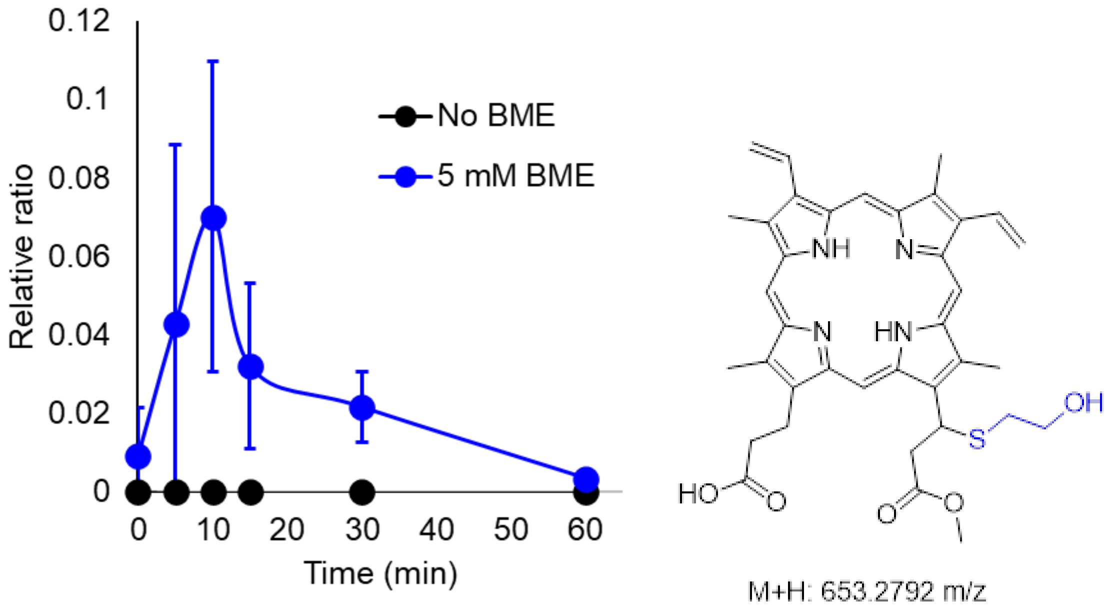


**Figure S21**. Formation of the *575* + BME Michael addition product with time during a BchE reaction. A peak with *m/z* correlating to the M+H *m/z* signal of the Michael addition product shown on the left is plotted relative to an internal standard of tryptophan. When no BME is present (*black*), no Michael addition product is observed. Reactions were conducted in triplicate with error bars representing the standard deviation for each timepoint.

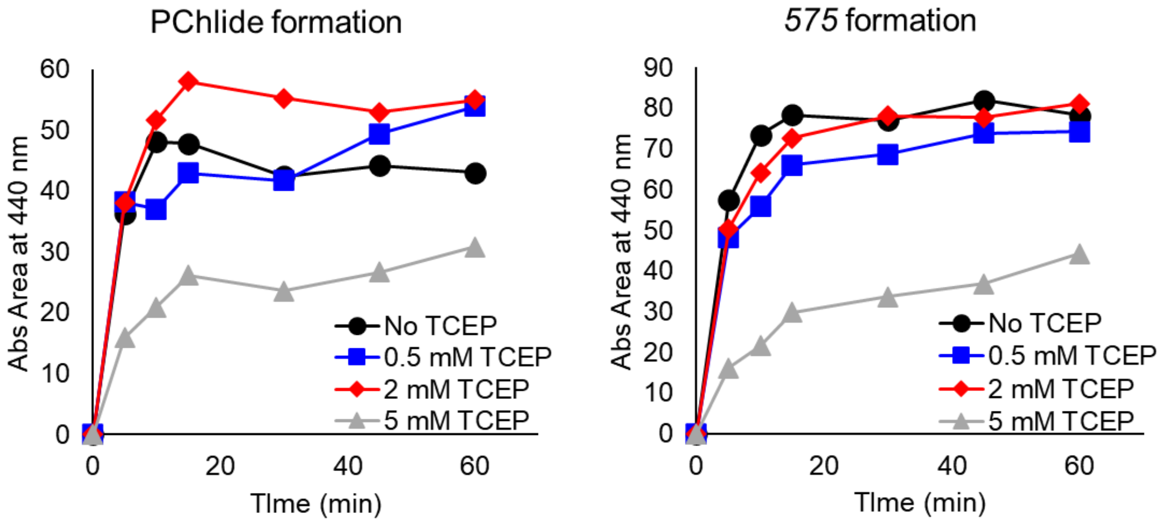


**Figure S22**. Formation of PChlide and 575 during the BchE reaction when incubated with varying amounts of reductant, TCEP. The concentrations 0, 0.5, 2, and 5 mM TCEP were chosen to replicate that used for BME shown in the main text.

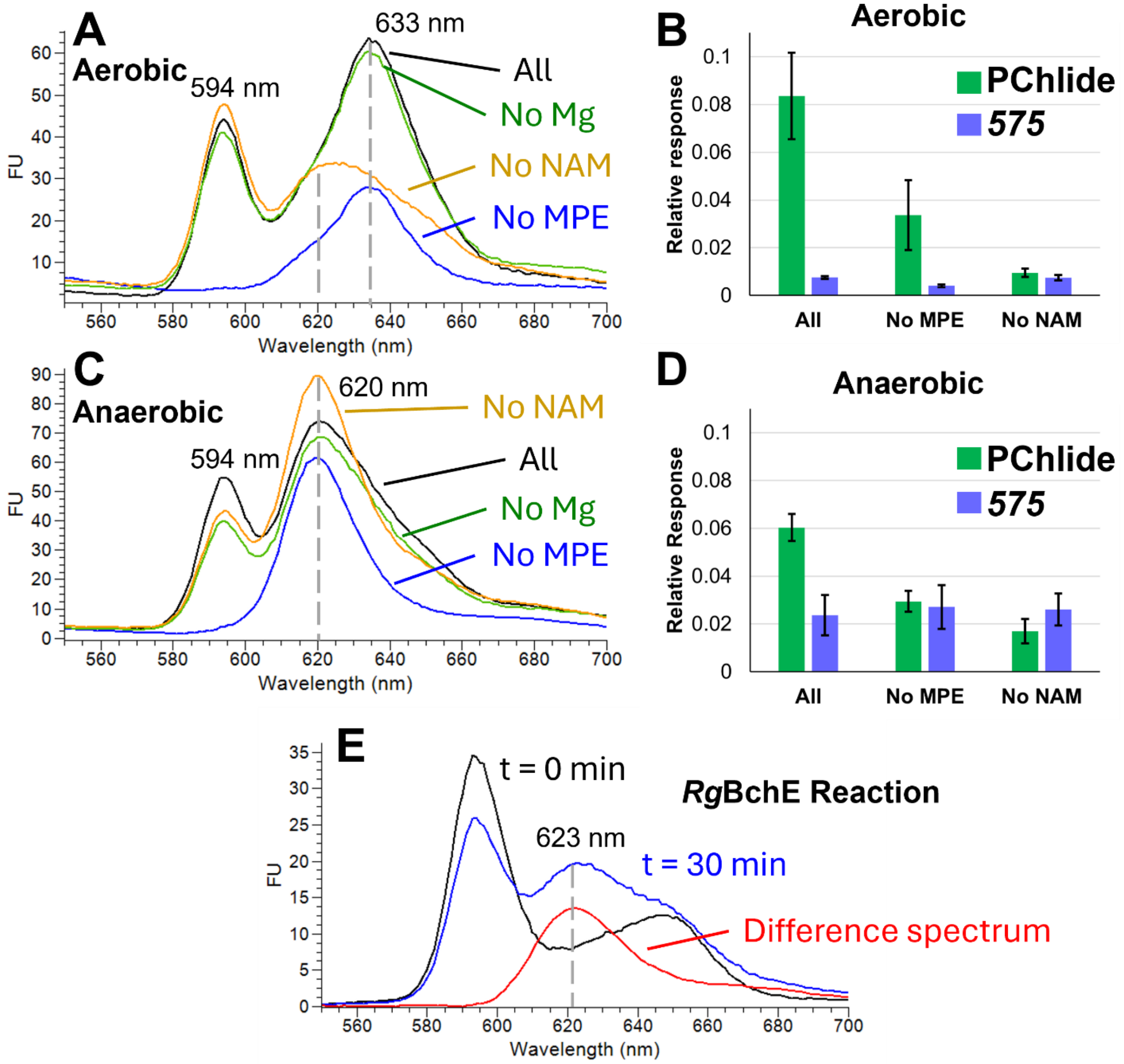


**Figure S23.** Analysis of *R. capsulatus* cells used to perform the BchE reaction *in vivo*. A full description of the experiment is given in Methods; briefly, *R. capsulatus* cells were grown aerobically in the dark or anaerobically under illumination. These cells were freeze-thawed and incubated with MPE, nicotinamide (NAM), and magnesium chloride to induce native BchE activity. The cells were washed and resuspended in ammoniacal acetone and hexanes. The acetone phase was analyzed either by fluorescence spectroscopy with excitation at 440 nm (A & C) or by high-resolution LC-MS after addition of sulfuric acid and tryptophan (internal standard) (B,D). Fluorescence spectra from aerobically grown cells (A) show a peak at 633 nm corresponding to PChlide, and a peak at 594 arising from the substrate MPE, which is not present when MPE is omitted. The overall trends mirror those reported by Gough et al., with maximal PChlide accumulation when all components (MPE, NAM, and Mg) are present, with a clear dependence on each.[18] Notable differences include a reduced dependence on Mg and detectable, though lower, levels of PChlide even without added MPE. However, the NAM dependence and blue-shift behavior were reproduced.

Anaerobically grown, illuminated cells yielded markedly different spectra (C), with a dominant emission at 620 nm rather than 633 nm. This 620 nm species increased in the absence of NAM and showed a similar dependence on MPE and Mg as the aerobic PChlide signal. In earlier work with the *R. gelatinosus* BchE homolog, we observed only the 575 species and were unable to achieve full conversion to PChlide; fluorescence monitoring of those reactions revealed a peak at 623 nm (E), closely resembling the anaerobic *R. capsulatus* spectra. The small difference in emission maxima likely reflects solvent effects, as ammoniacal acetone was used for *R. capsulatus* extractions and ammoniacal methanol for quenching the *R. gelatinosus* reactions.

To test whether the 575 species is present in vivo, we repeated the experiment for HRMS analysis. Panels B and D show the relative abundances of PChlide and the 575 species in extracts from *R. capsulatus* cells (n = 7; error bars represent one standard deviation). Aerobically grown cells displayed a dependence on MPE and NAM, with only low levels of the 575 species across conditions. In contrast, anaerobically grown cells accumulated substantially less PChlide and consistently higher levels of the 575 species, consistent with the fluorescence data for the 620 nm emitter.

Overall, these results demonstrate that the 575 species is formed in vivo and that its accumulation is strongly influenced by growth conditions. Based on our current understanding, we speculate that differences in intracellular reduction potential—driven primarily by the presence or absence of oxygen—underlie the distinct accumulation patterns.

**_Rp_BchE-MBP DNA sequence**

Synthesized and cloned into pET28a(+) vector at *Nco*I and *Xho*I sites (*Nco*I and *Xho*I sites bolded at 5'- and 3'- ends respectively)

**5'-**
**CCATGG**GCCGTGTACTTCTGATACATCCAAATTATCATAGCGGTGGGGCAGAGATCGC
GGGAAATTGGCCCCCTGCGTGGGCCGCTTATCTCACCGGCTATCTTAAGCATGGCGGC
TATACTGATGTCCATTTTGTCGACGCGATGACACACCACATAGACGAAGCCGGTGTTC
GAGCTAAAATAGCGGAACTTCAACCAGACATAGTTGGCTGTACAGCGATAACCCCCG
CAATCTACCAAGCGGAAGCGACGCTGCAATGGGCCAAGGAAATAAACCCGGACATAG
TTACTGTGCTGGGCGGTATTCATGGAACGTTCATGTATCCCCAGGTACTGGCAGAGGC
ACCTTGGATCGATGCCGTCGTACGAGGTGAAGGTGAGGCGGTCTTCCTGAACTTTGT
GCGAGCAGTTGATGATGGTAGCTGGGCAAGAGATAGACACTCTGTTCGGGGCATAGC
TTTCTTAGAAGACCAGGCCGCCGTAGACAAGAAGGTTCGCGCTACCGAGGCAGAAC
CCCCGATTGCAGACTTGGATACGATTCGTCCTGATTGGGGCATTCTTCAGTGGGACAG
TTACCTGTATATACCGATGAACACTCGCGTGGCAATCCCTAATTTTGCGCGTGGCTGCC
CGTTCACCTGCACTTTCTGCTCGCAATGGAAGTTTTGGCGCGACTATCGCATACGTGA
TCCAATTAAGGTAGTGGATGAAATCGAGGACCTGGTTAAGAATCACCAGGTAGGTTTC
TTCATACTTGCCGACGAAGAGCCGACAATACACCGCAAGAAATTTATAGCATTTTGTG
AAGAGCTGATAAAGCGCGACTTAGGGGTTCTGTGGGGTATTAATACCCGCGTTACGGA
CATTCTGAGAGACGAAAAGCTTCTGCCTTTGTTTCGCCAGGCGGGTCTGATCCATGTG
TCCCTCGGCACGGAAGCTGCTGCACAACTCAAACTGGAACGTTTTAATAAAGAGACA
ACTATCGCGCAGAACAAGCGCGCGATACAGCTTCTTAGAGAAGCGGGAATAGTAACT
GAGGCCCAATTCATTGTCGGACTGGAGAATGAGACCGCGGAAACCCTGGAAGAGAC
TTACCGCATGGCGCGTGACTGGAACCCTGATATGGCCAACTGGGCTATGTATACACCG
TGGCCTTTTAGCGATCTGTTTCAGGAATTGGGTGACAAGGTTGAGGTGTTTGATTTTG
CCAAGTACAATTTCGTTACGCCCATTATGAAACCGGATGCGATGGAGCGCGGCGAGTT
GTTGGATAGAGTTATGAACAATTATCGGCGGTTCTTCATGAACAAGAGTTTCTTCCAG
TACCCCTGGGAGAAAGATAAATTACGCCGGAAGTACCTCATGGGTTGTCTGAAGGCC
TTCCTCAAGTCGGGATTCCAGCGGACGTTTTACGACTTAGGCCGTGTAGGCTATTGGG
GTCCTCAGACAAAGAAGAAGGTAGACTTTGCGTTTGATACCACCCGCCAGTATTCTA
GACCCAGCGCTGATGCGGTTGCGGCGGCGGATGCAGGGTGGGTTACAATGCACGGCC
CCAAGATTGAGCATAAACGGCGTCAAGGTGAACTGAATGCAGCTCTGGCTTGCGGTG
GCGGCACCGAACAGTTAGCCGAAAGTGTTGCAGAGAGCGTGGCGGATACCCGGCAA
GCCACAGCGGAAAACCTTTACTTTCAAAGCGCTAGCAAGACTGAGGAAGGTAAATTA
GTGATATGGATTAACGGCGACAAGGGATATAATGGTCTGGCTGAAGTGGGTAAGAAAT
TCGAGAAAGATACGGGAATAAAAGTTACGGTGGAACATCCCGATAAGTTAGAAGAGA
AGTTCCCCCAAGTGGCCGCAACAGGTGACGGCCCTGATATAATCTTTTGGGCCCATGA
TCGCTTCGGCGGTTACGCACAAAGTGGGCTGCTGGCAGAGATCACCCCCGATAAGGC
ATTTCAAGACAAACTTTATCCATTTACCTGGGACGCCGTGAGATACAATGGTAAACTT
ATAGCATACCCAATCGCCGTGGAAGCGCTTAGTCTGATCTATAACAAAGACCTGCTTC
CGAACCCACCAAAGACATGGGAAGAAATTCCTGCGTTGGATAAAGAGCTGAAAGCC
AAGGGTAAAAGTGCGTTAATGTTTAACCTTCAAGAACCGTACTTCACCTGGCCGCTTA

TAGCCGCCGACGGTGGCTACGCATTCAAGTACGAGAACGGGAAGTACGATATCAAGG
ATGTTGGAGTGGATAATGCGGGTGCAAAAGCGGGACTGACATTCCTTGTAGACCTGA
TAAAGAACAAGCATATGAATGCCGATACAGACTATAGTATCGCTGAGGCGGCATTTAA
CAAGGGAGAAACGGCAATGACTATTAACGGCCCCTGGGCTTGGTCTAACATCGATAC
GTCCAAGGTAAATTACGGAGTGACAGTACTGCCCACCTTTAAAGGTCAGCCGTCAAA
GCCATTTGTAGGGGTTTTGTCAGCTGGAATTAACGCCGCAAGTCCCAACAAGGAGCT
TGCAAAAGAATTCTTAGAAAACTACCTTTTGACAGACGAGGGCTTAGAAGCCGTAAA
TAAGGATAAACCTCTGGGCGCAGTAGCGCTGAAATCCTATGAAGAGGAGTTGGTGAA
AGACCCGAGAATTGCTGCAACAATGGAGAACGCGCAAAAGGGCGAGATTATGCCGA
ATATCCCGCAGATGTCTGCATTTTGGTACGCGGTGCGCACGGCTGTCATTAACGCGGC
CAGCGGACGTCAAACGGTTGACGAAGCTCTGAAGGATGCTCAAACGAGTGCACATC
ACCATCATCACCATCACCATCACCATTAA**CTCGAG**-3’

**Amino acid sequence of *Rp*BchE-MBP** (TEV protease site and 10× His tag are bolded; C-terminal MBP-tag is underlined)

MGRVLLIHPNYHSGGAEIAGNWPPAWAAYLTGYLKHGGYTDVHFVDAMTHHIDEAGV
RAKIAELQPDIVGCTAITPAIYQAEATLQWAKEINPDIVTVLGGIHGTFMYPQVLAEAPWI
DAVVRGEGEAVFLNFVRAVDDGSWARDRHSVRGIAFLEDQAAVDKKVRATEAEPPIADL
DTIRPDWGILQWDSYLYIPMNTRVAIPNFARGCPFTCTFCSQWKFWRDYRIRDPIKVVDEI
EDLVKNHQVGFFILADEEPTIHRKKFIAFCEELIKRDLGVLWGINTRVTDILRDEKLLPLFR
QAGLIHVSLGTEAAAQLKLERFNKETTIAQNKRAIQLLREAGIVTEAQFIVGLENETAETL
EETYRMARDWNPDMANWAMYTPWPFSDLFQELGDKVEVFDFAKYNFVTPIMKPDAM
ERGELLDRVMNNYRRFFMNKSFFQYPWEKDKLRRKYLMGCLKAFLKSGFQRTFYDLG
RVGYWGPQTKKKVDFAFDTTRQYSRPSADAVAAADAGWVTMHGPKIEHKRRQGELNA
ALACGGGTEQLAESVAESVADTRQATA**ENLYFQS**ASKTEEGKLVIWINGDKGYNGLAEV
GKKFEKDTGIKVTVEHPDKLEEKFPQVAATGDGPDIIFWAHDRFGGYAQSGLLAEITPDK
AFQDKLYPFTWDAVRYNGKLIAYPIAVEALSLIYNKDLLPNPPKTWEEIPALDKELKAKG
KSALMFNLQEPYFTWPLIAADGGYAFKYENGKYDIKDVGVDNAGAKAGLTFLVDLIKN
KHMNADTDYSIAEAAFNKGETAMTINGPWAWSNIDTSKVNYGVTVLPTFKGQPSKPFV
GVLSAGINAASPNKELAKEFLENYLLTDEGLEAVNKDKPLGAVALKSYEEELVKDPRIAA
TMENAQKGEIMPNIPQMSAFWYAVRTAVINAASGRQTVDEALKDAQTSA**HHHHHHHH
HH***

***Rc*FdxA DNA sequence** cloned into pET26b(+) vector at *Nde*I and *Xho*I restriction sites (bolded)

**CATATG**ACCTACGTCGTCACGGATAACTGTATCGCCTGTAAATATACAGATTGCGTCG
AAGTATGTCCGGTCGATTGCTTTTACGAGGGTGAGAACACGTTAGTCATCCACCCTGA
CGAGTGCATCGATTGTGGTGTTTGCGAGCCGGAATGCCCTGCGGATGCGATCCGCCC
AGATACTGAACCAGGTATGGAAGATTGGGTGGAGTTTAACCGTACTTACGCCTCGCAG
TGGCCGGTAATCACGATCAAGAAAGACCCAATGCCGGACCACAAGAAGTACGATGGT
GAAACGGGTAAACGTGAAAAATACTTTTCACCGAATCCTGGCACGGGGGAT**CTCGA
G**

***Rc*FdxA amino acid sequence** C-terminal 6× His tag boldened

MTYVVTDNCIACKYTDCVEVCPVDCFYEGENTLVIHPDECIDCGVCEPECPADAIRPDTE
PGMEDWVEFNRTYASQWPVITIKKDPMPDHKKYDGETGKREKYFSPNPGTGDLE**HHH
HHH***

***Rg*FdxA DNA sequence** cloned into pET26b(+) vector at *Nde*I and *Xho*I restriction sites (bolded)

**CATATG**ACCCACGTGGTCCTTGACTCCTGCATTCGTTGTAAATACACGGACTGTGTGG
ATGTTTGTCCTGTTGACTGCTTTCGGGAAGGCCCGAACTTTTTGGTTATCGACCCGGA
GGAGTGCATCGATTGTGCTGTGTGTATCCCGGAGTGTCCTGCAAATGCAATCCTTCCT
GAGGAAGACGTACCAAGTGATCAATTACAATTCGTGCAACTCAACGCGGAACTCGCA
AAAGTTTGGCCGTCGATCACTAAGCGTAAAGGGTCCCTCCCTGATGCCGACGAATGG
AAAGACCGTAAAAATAAACTGCCGCATCTTCAACGTGAGAATCTCTACTTCCAATCT**C
TCGAG**

***Rg*FdxA amino acid sequence** C-terminal 6× His tag boldened

MTHVVLDSCIRCKYTDCVDVCPVDCFREGPNFLVIDPEECIDCAVCIPECPANAILPEEDV
PSDQLQFVQLNAELAKVWPSITKRKGSLPDADEWKDRKNKLPHLQRLE**HHHHHH***

***Rg*FNR (ferredoxin reductase) DNA sequence** cloned into pET26b(+) vector at *Nde*I and *Xho*I restriction sites (bolded)

**CATATG**TCTGCTTTTAACGAAGAGCGCGTATTGTCTGTCCATCATTGGACTGATCGCTT
ATTTACATTCACTACGACGCGGGATCAGTCGTTGCGGTTTTCTAACGGCCATTTCACA
ATGATTGGTTTGCGTGTCGAGGGCAAACCTTTACTTCGGGCTTACTCCATTGTGAGCC
CAAACTATGAAGAGCACCTCGAATTTCTGTCAATTAAAGTACCAAATGGTCCTCTCAC
CTCACGGTTGCAACATATCCAGGTTGGCGACTCAATTATCGTGGGTCGCAAGCCTACA
GGCACATTGTTAATTGATTACTTACTGCCAGGGAAACGGCTCTATCTGTTCTCTACTGG
GACTGGTTTGGCCCCATTTATGAGCATCATTCGTGACCCAGAAACGTATGAGAAATTT
GAAAAAGTTATTCTTGTTCATGGGGTCCGGCAGGTAGATGAGCTGGCGTACCACGAC
CTGCTTACTAAGAACCTCCCGGAACATGAGTTTCTTGGTGAAATGATTCAGAGTCAGT
TACTGTATTATCCGACCGTCACTCGGGAGAATTATCGGAACCGGGGCCGCATCACAGA
GCTCATCCAGAGCGGCAAAATGTTCGATGATTTAGACCTCCCGATGCTTGACCCAATC
CATGACCGGGTGATGATCTGTGGCAGCCCTGCCATGCTCCGTGACCTCAAGCACATGT
TGGAGGGCATGCGCTTCAAAGAGGGTAACACTACAACGCCTGGCGATTTCGTCATCG
AGCGTGCGTTTGCGGACCAAGAGAATCTCTACTTCCAATCT**CTCGAG**

***Rg*FNR (ferredoxin reductase) amino acid sequence** C-terminal 6× His tag boldened

MSAFNEERVLSVHHWTDRLFTFTTTRDQSLRFSNGHFTMIGLRVEGKPLLRAYSIVSPNY
EEHLEFLSIKVPNGPLTSRLQHIQVGDSIIVGRKPTGTLLIDYLLPGKRLYLFSTGTGLAPF
MSIIRDPETYEKFEKVILVHGVRQVDELAYHDLLTKNLPEHEFLGEMIQSQLLYYPTVTRE

NYRNRGRITELIQSGKMFDDLDLPMLDPIHDRVMICGSPAMLRDLKHMLEGMRFKEGNT
TTPGDFVIERAFADQENLYFQSLE**HHHHHH***

**DFT optimized coordinates for MPE, PChlide, olefin-*575*, and ring-*575* with acetic acid adduct**

**MPE**

| | | | |
|---|---|---|---|
| O | -6.15537219804414 | -1.25951475820968 | 1.04322468367421 |
| O | -6.19798842387878 | 2.78631521531166 | -2.11392042773606 |
| O | -6.54996841780911 | -3.44597748767992 | 1.38242349876292 |
| O | -6.38061971734844 | 2.32891230739904 | 0.07340125090128 |
| N | 2.83594810567577 | 1.39921468895467 | 0.42847385772985 |
| N | 2.81246173754539 | -1.52750975075837 | 0.22231447270242 |
| N | -0.09785471400738 | -1.47873619546671 | 0.23869399264967 |
| N | -0.08301475983355 | 1.42370930779260 | 0.50681391194559 |
| C | -2.15051118157413 | -2.51407037770126 | 0.03002850980245 |
| C | -2.12299748176813 | 2.51008220195224 | 0.58835182851516 |
| C | -1.44545892189689 | -1.25678290786860 | 0.22691502398825 |
| C | -1.19576191490733 | -3.48767831532844 | -0.08288685051884 |
| C | -1.43324036883971 | 1.23081787960475 | 0.47828861742789 |
| C | -1.14896228224581 | 3.46756550547098 | 0.67976382128946 |
| C | 0.09211915641471 | -2.81914249843920 | 0.04392730917758 |
| C | 0.12866707852531 | 2.76915211349765 | 0.62695534516402 |
| C | 2.63105939166942 | 2.74533583228170 | 0.56467716828814 |
| C | -3.63748546993006 | -2.64643700768990 | -0.02133369550475 |
| C | 2.59435637130038 | -2.86849058363977 | 0.00296062372097 |
| C | 4.16221892513418 | -1.29895955458153 | 0.06286983127112 |
| C | -3.60855942450910 | 2.67635256514085 | 0.55835565367089 |
| C | 4.18037370573438 | 1.19834431903810 | 0.28595591469640 |
| C | -2.05607723564433 | -0.00961237142547 | 0.35520330098133 |
| C | 3.89987973537570 | 3.44081302087505 | 0.53393222265214 |
| C | 4.83433372950372 | -2.54109061680747 | -0.22217852843713 |
| C | 3.86713954136131 | -3.52848144151827 | -0.23843981630642 |
| C | 4.87186342817594 | 2.48524135861634 | 0.33244949599075 |
| C | 1.38395536252268 | 3.36875433375809 | 0.67366542643379 |
| C | 1.32903462199558 | -3.45375378856832 | -0.05404652530565 |
| C | 4.78182013501961 | -0.05145699986159 | 0.14365631596602 |
| C | -4.27537792011009 | -2.62500427514238 | 1.38871407714921 |
| C | -1.38436522812454 | -4.95348833256570 | -0.29476964327235 |
| C | -1.31254964244984 | 4.94750363433796 | 0.79219814769043 |
| C | -4.18167060809685 | 2.62637009274120 | -0.86339723676869 |
| C | 4.06982397003603 | 4.92028146707950 | 0.63599655269980 |
| C | 6.30471273600825 | -2.69437639246372 | -0.42887438417056 |
| C | 4.11879535125261 | -4.94515946960321 | -0.46644278480598 |
| C | 6.29572482689327 | 2.75741744896046 | 0.18980927681153 |
| C | -5.77340056611436 | -2.51910034553072 | 1.29118594641687 |
| C | -5.72032640135831 | 2.58516305839490 | -0.96591374159202 |
| C | 3.45009804970988 | -5.98029046733582 | 0.04991453265894 |
| C | 7.18731933839785 | 2.09606743298067 | -0.55457336970763 |

| | | | |
|---|---|---|---|
| C | -7.56316401173518 | -1.02440466032059 | 0.83627517977075 |
| H | -4.05803119617671 | -1.82600436206800 | -0.60630786910347 |
| H | -3.92071232940522 | -3.57181267225414 | -0.52442832376515 |
| H | -4.08252035266707 | 1.89273517719306 | 1.15128604233182 |
| H | -3.88649370756375 | 3.62311963225048 | 1.02548073556192 |
| H | -3.13707841949657 | 0.00057074159251 | 0.33627811998457 |
| H | 1.39786718176791 | 4.44552315894704 | 0.77882039614075 |
| H | 1.29957886782799 | -4.51445238082935 | -0.25227280889299 |
| H | 5.85738922292431 | -0.05791081183544 | 0.05715540387898 |
| H | -4.02303970619789 | -3.53721492461490 | 1.92707048477734 |
| H | -3.89378511207336 | -1.76952618123680 | 1.94577196404345 |
| H | -0.86560564121294 | -5.29548840786778 | -1.19393822491946 |
| H | -2.43915662912406 | -5.20559225671014 | -0.39794198365717 |
| H | -0.98364784636818 | -5.52949197394587 | 0.54366979972299 |
| H | -0.78256649371975 | 5.34306033317388 | 1.66247822476957 |
| H | -2.36330060890818 | 5.22064328816317 | 0.88361636499138 |
| H | -0.91053266712903 | 5.45969904535302 | -0.08646098178277 |
| H | -3.80974899499701 | 1.73380517597130 | -1.37640029490067 |
| H | -3.81942169720664 | 3.47719992382325 | -1.44554200035969 |
| H | 5.11157479415518 | 5.19240683221760 | 0.80148565835520 |
| H | 3.74065372026537 | 5.41986481840935 | -0.28004371606715 |
| H | 3.47698084661224 | 5.33083927965114 | 1.45576624701925 |
| H | 6.69386033603345 | -1.92092197965162 | -1.09370595204735 |
| H | 6.54889543133734 | -3.66482926180552 | -0.85900695362533 |
| H | 6.84673553661444 | -2.60788562353244 | 0.51751251425007 |
| H | 4.97166956292272 | -5.16471263499249 | -1.10267147582870 |
| H | 6.64984447108572 | 3.62725458666731 | 0.73638765625921 |
| H | 3.73885386630301 | -6.99571115795007 | -0.19009148994543 |
| H | 2.62175653258088 | -5.85692978367459 | 0.73534707177127 |
| H | 8.22327964605637 | 2.40998005501484 | -0.57589169098910 |
| H | 6.92132734656472 | 1.25218893835585 | -1.17739036504693 |
| H | -7.64344143612601 | 0.03365228469063 | 0.60506758749276 |
| H | -8.12114999189970 | -1.26746593272086 | 1.73965458586551 |
| H | -7.92590277321724 | -1.62895113879918 | 0.00611199693700 |
| Mg | 1.35985420064342 | -0.05774316199560 | 0.66260403605877 |
| H | 2.93184529822208 | -1.76298516103664 | 2.10108360288366 |
| O | 2.89697028200291 | -1.80857514425632 | 3.09195207500915 |
| O | 1.37529694402014 | -0.20872689691987 | 2.76640579579622 |
| C | 2.01012330702829 | -0.93153809975708 | 3.52808566683015 |
| C | 1.85912951538021 | -0.90660398890888 | 5.01151498105560 |
| H | 1.11316602334236 | -0.17243506279664 | 5.30063175872378 |
| H | 1.56846623197162 | -1.89867413375932 | 5.36133813565939 |
| H | 2.82124552567658 | -0.66668869905660 | 5.46764432466770 |

**PChlide**

| | | | |
|---|---|---|---|
| O | -4.64476053329811 | -0.91789701733517 | -1.77763707685664 |

| | | | |
|---|---|---|---|
| O | -5.76621861596439 | 1.50145736809880 | 2.50447930921805 |
| O | -2.67730811201308 | -0.58593106326408 | -2.81411754983615 |
| O | -5.86841351577678 | 2.00730071148098 | 0.32212343630700 |
| N | 3.46267773898777 | 0.91251934961777 | 0.20998630411874 |
| N | 3.66161666224283 | -2.01619836105072 | 0.13625163711091 |
| N | 0.85552226885239 | -2.13133871580164 | -0.13568851113993 |
| N | 0.50338340358613 | 0.73932700714943 | 0.05866572360593 |
| C | -1.14032398856877 | -3.18217713114322 | -0.47276349686095 |
| C | -1.58991206869265 | 1.73836709430533 | -0.02224513722139 |
| C | -0.45915061531778 | -1.94700482010441 | -0.27698714588604 |
| C | -0.18088863374261 | -4.18499987056232 | -0.46455840017238 |
| C | -0.84611739627434 | 0.48256812070660 | -0.08112774276288 |
| C | -0.66039656594696 | 2.72955872644732 | 0.14000824578049 |
| C | 1.07374999230161 | -3.48941924014878 | -0.25631976945422 |
| C | 0.64586745298262 | 2.08654399690820 | 0.17615640441724 |
| C | 3.15227612537822 | 2.24238229950564 | 0.27441117993433 |
| C | -2.55645270279248 | -2.91333064298817 | -0.58568318483485 |
| C | 3.55407383787179 | -3.38259629378347 | -0.04330327709866 |
| C | 4.99505573105251 | -1.70198654806690 | 0.09541632599084 |
| C | -3.07508376766499 | 1.90789246745197 | -0.04214178328782 |
| C | 4.82327108914406 | 0.80009147012395 | 0.17886880949304 |
| C | -1.29690040406798 | -0.82078618994197 | -0.26483760292982 |
| C | 4.37324715959987 | 3.02812755973534 | 0.29825072584252 |
| C | 5.78021784293557 | -2.90523975790863 | -0.05899845079817 |
| C | 4.89132622095231 | -3.95781425765329 | -0.12272355484796 |
| C | 5.41822311904257 | 2.13934092390907 | 0.20899876315736 |
| C | 1.86214116239525 | 2.76977694364555 | 0.28601755661961 |
| C | 2.35056505326849 | -4.06165583557013 | -0.20736015352049 |
| C | 5.51927737714775 | -0.40586124253494 | 0.16016917428316 |
| C | -2.70693514526666 | -1.33682939591080 | -0.51348736589473 |
| C | -0.35632128602683 | -5.65368136784339 | -0.64085480748376 |
| C | -0.88967319472879 | 4.19815693764261 | 0.27586791219336 |
| C | -3.70960948609811 | 1.64144995538392 | 1.32681339983914 |
| C | 4.43113539942655 | 4.51794900493954 | 0.33996787937854 |
| C | 7.27095565375591 | -2.95476978546732 | -0.10466759167038 |
| C | 5.25795825338892 | -5.36201682866384 | -0.25178480713705 |
| C | 6.82738664405326 | 2.50210687565711 | 0.14354103958766 |
| C | -3.31204902095897 | -0.88785227046894 | -1.82892785806102 |
| C | -5.24872132976859 | 1.72595488461998 | 1.37839456976064 |
| C | 4.61169467853383 | -6.41343280140527 | 0.25922320954705 |
| C | 7.80019834146653 | 1.86910293206005 | -0.51855500787018 |
| C | -5.34701656680826 | -0.56880301329164 | -2.98915186166849 |
| H | -3.31937736738968 | 2.92427251342255 | -0.35181558063210 |
| H | -3.53861042040198 | 1.26189687662466 | -0.78412708085988 |
| H | 1.79190349619023 | 3.84666253341742 | 0.36002742409312 |
| H | 2.40895499759567 | -5.12842639259643 | -0.36336312299213 |

| | | | |
|---|---|---|---|
| H | 6.59613135932975 | -0.33664615573413 | 0.16876607248437 |
| H | -3.40886944526407 | -1.10803626280999 | 0.28834026638025 |
| H | 0.10267637265082 | -5.99192385183757 | -1.57416801350581 |
| H | -1.41182830519314 | -5.91868927320847 | -0.66580006350670 |
| H | 0.12198108916105 | -6.20935556085906 | 0.16906828934120 |
| H | -0.56580918628233 | 4.55952723681916 | 1.25578288671141 |
| H | -1.94361032095814 | 4.44738411871524 | 0.16215362196381 |
| H | -0.32697990152488 | 4.75997707581752 | -0.47402623997629 |
| H | -3.31273599021216 | 2.35235947976709 | 2.05731359372799 |
| H | -3.42080173349304 | 0.65329990783034 | 1.69538069787266 |
| H | 5.44172562114666 | 4.87298852726167 | 0.53709679216770 |
| H | 4.10854675964929 | 4.95027855962551 | -0.61177489697510 |
| H | 3.77237599982396 | 4.91794206683096 | 1.11344196147585 |
| H | 7.66903990539619 | -2.23844266201316 | -0.82662087176823 |
| H | 7.63006893296183 | -3.94603959062882 | -0.37700877256160 |
| H | 7.70126089897906 | -2.70191818626840 | 0.86863904346014 |
| H | 6.17960275835158 | -5.54981508416314 | -0.79508898131619 |
| H | 7.08991852056508 | 3.41496591565977 | 0.67077390915263 |
| H | 4.98630566511788 | -7.41724362381408 | 0.10436274845818 |
| H | 3.71681867618874 | -6.31273341933928 | 0.85945766856535 |
| H | 8.81442819450283 | 2.24704070556408 | -0.49638901606430 |
| H | 7.62195455348907 | 0.98479364779061 | -1.11608499537062 |
| H | -6.40143290497114 | -0.65592023510278 | -2.74618145955633 |
| H | -5.07756382613486 | -1.25568188693210 | -3.78978939495761 |
| H | -5.10397004232748 | 0.45228498176823 | -3.27895420296237 |
| Mg | 2.09170260550691 | -0.60437460065563 | 0.40096385552069 |
| O | -3.50055605877571 | -3.66815725701782 | -0.71282895328537 |
| H | 3.59744219472658 | -2.20480667375261 | 2.06493410148685 |
| O | 3.45400046404587 | -2.21209228294473 | 3.04409416319997 |
| O | 1.91265145712225 | -0.69728403509028 | 2.48537405559790 |
| C | 2.48996337461133 | -1.35876906757220 | 3.34234977716312 |
| C | 2.17702848685726 | -1.28054034919338 | 4.79794826450436 |
| H | 3.07594341252584 | -0.99095968093714 | 5.34496960773290 |
| H | 1.38184647466543 | -0.56218041309946 | 4.97276043818936 |
| H | 1.87884748443762 | -2.26768073365920 | 5.15561064970917 |

**Olefin-*575***

| | | | |
|---|---|---|---|
| O | -6.36661487740690 | -3.56771687647683 | -0.04317987730141 |
| O | -6.56894793777333 | 3.20314578983998 | -1.12136907894237 |
| O | -6.90781286417629 | -1.56407801542054 | -0.91397670346287 |
| O | -6.56957190051655 | 1.95756196645840 | 0.74301572278008 |
| N | 2.64277596295168 | 1.41662805645338 | 0.54121046281461 |
| N | 2.66960537063318 | -1.49626017484996 | 0.22829841962904 |
| N | -0.25051028646829 | -1.51781485422443 | 0.38322093795607 |
| N | -0.25991158679588 | 1.37054003996035 | 0.79054632594613 |
| C | -2.27916693295116 | -2.58610995627550 | 0.16470028600712 |

| | | | |
|---|---|---|---|
| C | -2.30967218395287 | 2.39457206673730 | 1.10839998975469 |
| C | -1.59745432919221 | -1.31866329160400 | 0.41274108248607 |
| C | -1.28953273002801 | -3.54185633469879 | 0.01980055265244 |
| C | -1.60397868605911 | 1.14298496684680 | 0.85887271735804 |
| C | -1.35248171841192 | 3.36945244169550 | 1.18206097523137 |
| C | -0.02869423169940 | -2.84787129383401 | 0.13565632281872 |
| C | -0.06890011237771 | 2.70890272511357 | 0.98303904278060 |
| C | 2.42062738394750 | 2.75046807445127 | 0.74722859291349 |
| C | -3.69231943856695 | -2.84232755519374 | 0.03605633302954 |
| C | 2.47212044140196 | -2.83957814292956 | -0.00191408067414 |
| C | 4.00278753124440 | -1.23504108970325 | 0.01166623695115 |
| C | -3.79546598068662 | 2.53108256457982 | 1.18990222871803 |
| C | 3.97879816128975 | 1.25345978966365 | 0.30360187789561 |
| C | -2.21343626866352 | -0.09596820218389 | 0.69348502691335 |
| C | 3.67062307739086 | 3.47801555924730 | 0.65781788515635 |
| C | 4.69113980627206 | -2.45737340844805 | -0.32284899021879 |
| C | 3.74861446239532 | -3.46680661429544 | -0.30827224046789 |
| C | 4.64477598037473 | 2.55544972781444 | 0.35128928651193 |
| C | 1.17379279496949 | 3.33782795244606 | 0.97535672051803 |
| C | 1.22180083979897 | -3.45021172499876 | -0.01990739371815 |
| C | 4.59920753728331 | 0.02549384154038 | 0.08866149730477 |
| C | -4.64898915898525 | -1.98087173627502 | -0.35666844701932 |
| C | -1.45353454718445 | -4.99451012382419 | -0.27485488299438 |
| C | -1.53798178107938 | 4.83482665891283 | 1.39946688779778 |
| C | -4.45574223869685 | 2.60076407870693 | -0.19407597948444 |
| C | 3.81545415753947 | 4.95563520319695 | 0.80793073108815 |
| C | 6.15405551012377 | -2.57034791603965 | -0.59630436816928 |
| C | 4.01968406820064 | -4.87533693299270 | -0.56313379797041 |
| C | 6.04826037128588 | 2.86546410700362 | 0.11616780890327 |
| C | -6.06701685532628 | -2.32791825931886 | -0.47073571343147 |
| C | -5.99707662972971 | 2.59406686197908 | -0.17904982315972 |
| C | 3.39815259837038 | -5.92726107590486 | -0.02271085101511 |
| C | 6.89292552753314 | 2.25209609794010 | -0.71832293697249 |
| C | -7.74383155132544 | -3.96924474844702 | -0.15237970691425 |
| H | -4.00169076396022 | -3.86037505227485 | 0.24612314691227 |
| H | -4.22443631753890 | 1.68950538273590 | 1.73527047465991 |
| H | -4.05349100037700 | 3.42902789010649 | 1.75579921138732 |
| H | -3.28780419998744 | -0.10790292719160 | 0.78706893094512 |
| H | 1.17263480412293 | 4.40700007105375 | 1.14023816955789 |
| H | 1.20923393034467 | -4.50726322864084 | -0.23713695851446 |
| H | 5.66813219288933 | 0.04787070612783 | -0.05771303026796 |
| H | -4.42617451197416 | -0.96910863041694 | -0.66245036334514 |
| H | -1.28884438711868 | -5.20117832219790 | -1.33661167081950 |
| H | -2.45548805945933 | -5.34318069611317 | -0.02640102011435 |
| H | -0.73567480658401 | -5.59531546856540 | 0.28579157801451 |
| H | -0.94215571295960 | 5.19252712554756 | 2.24311729109341 |

| | | | |
|---|---|---|---|
| H | -2.58214169418592 | 5.07268837309820 | 1.59932600128974 |
| H | -1.22714084508039 | 5.40903592194556 | 0.52219913084002 |
| H | -4.14277585308482 | 1.73351578478566 | -0.78599507722942 |
| H | -4.10827104097389 | 3.48439827036971 | -0.73245279407355 |
| H | 4.86049339197347 | 5.24594643925564 | 0.90874140733015 |
| H | 3.40982521847343 | 5.48247159276997 | -0.06093526153682 |
| H | 3.27489266706048 | 5.31875737390927 | 1.68427729540289 |
| H | 6.48632934498897 | -1.79995728591725 | -1.29505336519125 |
| H | 6.40786650162418 | -3.54224740734460 | -1.01706256032193 |
| H | 6.73670502018826 | -2.44528571455229 | 0.32103761517900 |
| H | 4.84387313005582 | -5.07143051131036 | -1.24310933400023 |
| H | 6.42430051837631 | 3.72263388348119 | 0.66792232615400 |
| H | 3.69441682780735 | -6.93497621465211 | -0.28492853891645 |
| H | 2.60352054230651 | -5.82539824123780 | 0.70493386086998 |
| H | 7.91791413439157 | 2.58947294734053 | -0.80621350062047 |
| H | 6.59605118294824 | 1.42552578401872 | -1.35046442303047 |
| H | -7.78154055846538 | -4.98381245795892 | 0.23384710541143 |
| H | -8.06520053855926 | -3.94588286511194 | -1.19309899744373 |
| H | -8.37891397060165 | -3.31232551220646 | 0.44068841475576 |
| Mg | 1.21279900809840 | -0.08380214827105 | 0.79756397818733 |
| H | 2.93609954353457 | -1.77680433928810 | 2.12518622485827 |
| O | 2.94621502281472 | -1.84273065078578 | 3.11349107936905 |
| O | 1.32881904976232 | -0.32230424427662 | 2.88234768680740 |
| C | 2.03307049148869 | -1.02330963483657 | 3.60244303086678 |
| C | 1.93077647013877 | -1.01288149226813 | 5.09079433990552 |
| H | 2.20145523367804 | -0.01735013652396 | 5.44781253353839 |
| H | 0.89296471913057 | -1.19300117830544 | 5.37281828393814 |
| H | 2.58045528500857 | -1.75652267096458 | 5.54347128799748 |

**Ring-*575***

| | | | |
|---|---|---|---|
| O | -4.47876421339391 | -0.87228056416099 | -1.59571556478243 |
| O | -5.64533594702990 | 0.89830713958106 | 2.53919698168061 |
| O | -2.51055860202481 | -0.51557941310204 | -2.61949540636807 |
| O | -5.70226846244618 | 2.09468186417763 | 0.64336874749985 |
| N | 3.63912725052433 | 0.91836490878687 | 0.29018243844171 |
| N | 3.84976129498932 | -2.00595146788344 | 0.06557231067404 |
| N | 1.03965256770756 | -2.12671791553377 | -0.09244276278890 |
| N | 0.66990189259753 | 0.72623521215589 | 0.23856097053178 |
| C | -0.97795655080978 | -3.18088481586492 | -0.39494202439446 |
| C | -1.42484008503971 | 1.71251392579690 | 0.24302100132422 |
| C | -0.29057057816850 | -1.95116145010370 | -0.16874968066793 |
| C | -0.02698857080982 | -4.16144961721763 | -0.48195048894270 |
| C | -0.67652663204769 | 0.46770275566345 | 0.12238136960938 |
| C | -0.49743753882551 | 2.70572422121317 | 0.42772504990620 |
| C | 1.25290730219787 | -3.46886631441665 | -0.29064965151689 |
| C | 0.81052195410450 | 2.07361848854028 | 0.40919029161093 |

| | | | |
|---|---|---|---|
| C | 3.32118594704453 | 2.24115276066076 | 0.43928754062268 |
| C | -2.44997909439968 | -2.92731628361882 | -0.42929811671256 |
| C | 3.74010191593518 | -3.35558196145543 | -0.19197834977762 |
| C | 5.18354373253806 | -1.67544956885037 | -0.02158833248036 |
| C | -2.91174639301338 | 1.87469567111951 | 0.23198692682090 |
| C | 4.99862939285816 | 0.81839914050802 | 0.21373578561301 |
| C | -1.12881829616757 | -0.84174354777601 | -0.09280890881134 |
| C | 4.53341055347388 | 3.03084491409395 | 0.47413628495647 |
| C | 5.96107874308511 | -2.86013796712435 | -0.29115304730652 |
| C | 5.07185415315543 | -3.91435564491130 | -0.37597109823825 |
| C | 5.58457302366382 | 2.15442154874207 | 0.30729209396044 |
| C | 2.02484443129425 | 2.75553151152244 | 0.51501591301312 |
| C | 2.52615120429769 | -4.03561976836255 | -0.33115992020856 |
| C | 5.69975268162378 | -0.38408033386148 | 0.09749103786400 |
| C | -2.54918848922678 | -1.34524136595001 | -0.33800088033083 |
| C | -0.19417598776478 | -5.62697923772981 | -0.71480692042049 |
| C | -0.73649657622978 | 4.16660541891248 | 0.62345621816398 |
| C | -3.56081480659223 | 1.51879009873448 | 1.57471246114817 |
| C | 4.58626809296339 | 4.51717963095256 | 0.59818088869640 |
| C | 7.44733865592784 | -2.90278784975277 | -0.42246632390703 |
| C | 5.44141255307649 | -5.30320394934450 | -0.61466332693700 |
| C | 6.98809032870933 | 2.53707956192185 | 0.23584745641128 |
| C | -3.14101488201658 | -0.84601630615992 | -1.64050819065430 |
| C | -5.10231286068794 | 1.50133956204573 | 1.57574288528666 |
| C | 4.82656356336686 | -6.39630968557988 | -0.15389321619446 |
| C | 7.96457368185056 | 1.94871222438944 | -0.46196159487096 |
| C | -5.17542493255563 | -0.48090720197889 | -2.79615744125847 |
| H | -3.16348128460202 | 2.90470922634485 | -0.01886394008227 |
| H | -3.36276161190264 | 1.27107262636937 | -0.55342323966503 |
| H | 1.95108073967369 | 3.82741355617807 | 0.64355044198056 |
| H | 2.58078738507656 | -5.09228542532512 | -0.54591248695603 |
| H | 6.77554825051256 | -0.30703198534979 | 0.06662643261880 |
| H | -3.23227572608602 | -1.07158918522460 | 0.46354977937206 |
| H | 0.32431446676491 | -5.94957338947132 | -1.62185704861820 |
| H | -1.24853782901607 | -5.88292104600973 | -0.81989666871379 |
| H | 0.21578335149191 | -6.21321774363974 | 0.11207874554566 |
| H | -0.40556349606379 | 4.49291381546633 | 1.61338849242864 |
| H | -1.79363891844012 | 4.41191456582819 | 0.53092507831510 |
| H | -0.18715458292873 | 4.76310599538695 | -0.10938316927925 |
| H | -3.24014463488235 | 2.24262034047465 | 2.33156527955511 |
| H | -3.20881348547192 | 0.54798440146060 | 1.92920901912155 |
| H | 5.59575700551781 | 4.86479263893260 | 0.81427290026059 |
| H | 4.26270729497826 | 5.00184345728342 | -0.32779514950106 |
| H | 3.92745067625923 | 4.87259315568883 | 1.39297334734487 |
| H | 7.81483448176623 | -2.07773586390622 | -1.03551640507546 |
| H | 7.78218108133251 | -3.83574096642016 | -0.87436543727087 |

| | | | |
|---|---|---|---|
| H | 7.93252148123102 | -2.81867845546196 | 0.55465674459277 |
| H | 6.34207744150580 | -5.44405586976116 | -1.20579808343752 |
| H | 7.24599823732267 | 3.42934904938413 | 0.79981130768682 |
| H | 5.20430640079547 | -7.38201136434510 | -0.39424840980042 |
| H | 3.95608011896100 | -6.34945182012825 | 0.48736220043121 |
| H | 8.97314083679363 | 2.34109586548975 | -0.43136945329039 |
| H | 7.79506472044789 | 1.08857982843621 | -1.09602942433264 |
| H | -6.23136142654671 | -0.57341389142023 | -2.56107452114976 |
| H | -4.90563583606363 | -1.14024526957867 | -3.61974412906937 |
| H | -4.92960566342252 | 0.54888568004752 | -3.05127738017939 |
| Mg | 2.28292048815558 | -0.61576833333736 | 0.45349250268621 |
| H | -2.96797372440180 | -3.37334734004319 | 0.42156415102336 |
| H | -2.93469140035957 | -3.30740370908644 | -1.33032085874205 |
| H | 3.82679117338567 | -2.29644386870194 | 1.93448450265993 |
| O | 3.72909545932934 | -2.37370563584715 | 2.91997451892367 |
| O | 2.21630895892705 | -0.77556511199362 | 2.54926326841782 |
| C | 2.81406589861533 | -1.51266099802322 | 3.32750308010592 |
| C | 2.58810800598021 | -1.51577609218041 | 4.80163668772535 |
| H | 3.49567819486103 | -1.16659005459530 | 5.29893905847964 |
| H | 1.75554287384109 | -0.86601491365253 | 5.05447155476501 |
| H | 2.39731678902694 | -2.53369273152052 | 5.14327322056612 |
